# Biological causes and impacts of rugged tree landscapes in phylodynamic inference

**DOI:** 10.1101/2025.06.10.657742

**Authors:** Jiansi Gao, Marius Brusselmans, Luiz M. Carvalho, Marc A. Suchard, Guy Baele, Frederick A. Matsen

## Abstract

Phylodynamic analysis has been instrumental in elucidating epidemiological and evolutionary dynamics of pathogens. Bayesian phylodynamics integrates out phylogenetic uncertainty, which is typically substantial in phylodynamic datasets due to limited genetic diversity. Phylodynamic inference does not, however, scale with modern datasets, partly due to difficulties in traversing tree space. Here, we characterize tree space and landscape in phylodynamic inference and assess its impacts on analysis difficulty and key biological estimates. By running extensive Bayesian analyses of 15 classic large phylodynamic datasets and carefully analyzing the posterior samples, we find that the posterior tree landscape is diffuse yet rugged, leading to widespread tree sampling problems that usually stem from sequences in a small part of the tree. We develop clade-specific diagnostics to show that a few sequences—including putative recombinants and recurrent mutants—frequently drive the ruggedness and sampling problems, although existing data-quality tests show limited power to detect them. The sampling problems can significantly impact phylodynamic inferences or distort major biological conclusions; the impact is usually stronger on “local” estimates (*e.g*., introduction history) associated with particular clades than on “global” parameters (*e.g*., demographic trajectory) governed by general tree shape. We evaluate existing and newly-developed MCMC diagnostics, and offer strategies for optimizing phylodynamic analysis settings and mitigating sampling problem impacts. Our findings highlight the need and directions to develop efficient traversal over rugged tree landscapes, ultimately advancing scalable and reliable phylodynamics.

**Significance Statement:** Bayesian phylodynamics is central to epidemiological studies, but exploring the vast and complex tree space is computationally challenging. Phylodynamic datasets comprise many highly similar sequences, sampled through time, creating a uniquely structured landscape of optimal trees. Here, we show that phylodynamic tree landscapes are often highly rugged, with multiple peaks separated by difficult-to-cross valleys. These features lead to widespread sampling problems which are often driven by a few sequences. These problems can significantly impact phylodynamic estimates, especially those associated with particular clades, distorting biological conclusions. We develop diagnostics to identify problematic sequences and provide solutions to mitigate their impacts. We offer strategies to optimize phylodynamic analysis workflows and to develop algorithms for navigating rugged landscapes, thereby advancing infectious disease investigation.

---

Phylodynamic inference combines genetic and epidemiological data via phylogenetic modeling. This phylodynamic approach is key to infectious disease studies for elucidating pathogen biology and transmission dynamics. During the COVID-19 pandemic, unprecedented SARS-CoV-2 genomic data proved invaluable for identifying variants of concern (1, 2) and for assessing intervention measures (3, 4). Phylodynamic datasets of densely sampled heterochronous genomic sequences from measurably evolving populations (pathogens, cells; 5) are increasingly available. However, despite sheer data volume, the limited genetic diversity leads to significant uncertainty in phylogenetic inference (6, 7).

Bayesian phylodynamic inference incorporates various epidemiological and biological information (8, 9) and accommodates phylogenetic uncertainty by integrating over the space of possible phylogenies (“tree space”). This integration is typically achieved via Markov chain Monte Carlo (MCMC), which approximates the joint posterior through stochastic simulation. Many key evolutionary and epidemiological inferences (*e.g*., pathogen spread history and demographic trajectory) depend on the underlying phylogeny, making efficient tree space exploration crucial for scalable phylodynamics. However, due to challenges in traversing the vast and complex tree space and costly likelihood evaluations, Bayesian phylodynamic analysis of modern datasets is often computationally prohibitive.

Tree space is especially challenging to navigate when the optimality score landscape (“tree landscape”; 10, 11) is rugged, comprising complex topographical structures—*e.g*., peaks (12), terraces (13), and ridges (14)—where high-scoring regions are separated by low-scoring regions (“valleys”; 15). We focus on the “operational ruggedness” of tree landscapes: ruggedness with respect to the tree rearrangement moves used in phylogenetic MCMC (*e.g*., nearest-neighbor interchange, NNI; subtree prune-and-regraft, SPR). These can lead to sampling problems in phylodynamic inference, including lack of convergence between replicate chains and/or inadequate mixing within each chain (Fig. 1).

**Fig. 1.**
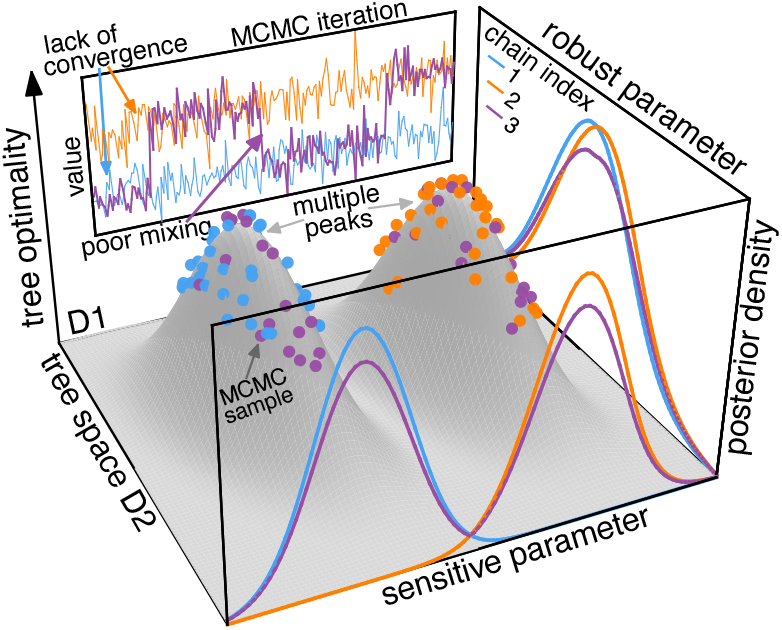
Conceptual illustration of a rugged tree landscape and consequent sampling problems. The *x* - and *y*-axes represent a two-dimensional tree space, while the *z*-axis indicates the optimality score (*e.g*., Bayesian posterior density). This tree landscape is multimodal, with two local peaks separated by a valley. Three MCMC replicate chains sample the posterior: chains 1 and 2 start from their respective peaks and remain stuck there, while chain 3 occasionally crosses the valley. If only one chain is run, chain 1 or 2 would suggest satisfactory sampling, whereas chain 3 would uncover both peaks but show inadequate mixing due to infrequent commutes between peaks. When multiple chains are run, comparing them reveals lack of convergence in sampling the tree. Tree landscape ruggedness may lead to poor sampling of tree-dependent parameters, manifested in MCMC trace plots (back panel) and discrepant posteriors among chains (front panel). Conversely, some parameters may be robust to tree sampling problems, with congruent estimates among chains (right panel) despite the ruggedness and poor tree sampling.

MCMC diagnosis is therefore critical in Bayesian phylodynamic analysis. Traditional diagnostics (*e.g*., effective sample size, ESS) focus on continuous parameters (16), while recent studies (17–21) advocate diagnosing tree topology given its central role. Tree landscape ruggedness and tree sampling problems have been a long-standing research focus (12, 15, 22–25), though primarily for topological inference with divergent species.

Phylodynamics differs from traditional phylogenetics by focusing on estimating epidemiological and evolutionary parameters rather than topologies. Bayesian phylodynamic inference typically marginalizes over time trees of heterochronous sequences with limited genetic diversity under coalescent tree priors. A comprehensive survey of phylodynamic tree space and landscape is thus much needed. Does limited genetic diversity lead to less rugged tree landscapes than phylogenomic datasets? How efficiently does MCMC explore phylodynamic tree space, and how much cost stems from the ruggedness? How robust are focal phylodynamic estimates to tree sampling problems, and which diagnostics best identify these problems and their causes? Can we identify biological factors contributing to the ruggedness before running costly phylodynamic analyses?

Here, to answer these critical questions, we comprehensively characterize phylodynamic tree space for a collection of landmark datasets and assess its impacts on inference difficulty and biological conclusions. Through exhaustively long and carefully diagnosed Bayesian phylodynamic analyses of these large datasets, we demonstrate that the tree space is frequently multimodal and sampling problems are widespread. By overlaying posterior densities over tree space and examining the unsampled trees in between peaks, we illustrate operationally rugged phylodynamic tree landscapes, with peaks of comparable height separated by deep valleys. We develop MCMC diagnostics to dissect tree sampling problems and show they often stem from a limited number of problematic sequences, whose removal dramatically improves sampling performance. We show that data-quality tests are of limited utility for detecting these problematic sequences. We further reveal that tree sampling problems can strongly impact key phylodynamic estimates, especially those associated with particular clades. Finally, we offer strategies to identify problematic sequences and optimize phylodynamic inference with modern datasets.

## Results

We collected 15 large phylodynamic datasets from representative high-impact empirical studies and performed comprehensive Bayesian phylodynamic analyses. For each dataset, we ran 10 independent MCMC chains using `BEAST` (39): five chains with 400 million iterations (“400M run”) and five with one billion iterations (“1B run”). We specified models, priors, and MCMC proposals following the original studies; the proposals were mostly identical to BEAUti defaults, reflecting standard empirical practices. We analyzed posterior outputs to visualize tree space and quantify sampling performance via existing and new MCMC diagnostics. Analysis details are in Materials and Methods and Suppl. Section S2.

### Biological Properties of Phylodynamic Datasets Underlie Tree Inference Difficulties

We quantified genetic diversity among sequences by performing mutation mapping over posterior trees. For most datasets, each branch typically experienced zero or one mutation, and most sites are invariant or parsimony-uninformative (Suppl. Fig. S1). Such limited genetic diversity leads to high uncertainty in tree inference. This is manifested in the large fraction of weakly supported clades subtended by short branches in summary trees (Fig. 2, Suppl. Fig. S3) and in diffuse posterior tree space (Fig. 3, Suppl. Fig. S4). Notable exceptions include the avian influenza (AIV) and Lassa datasets, where most branches experienced tens or more mutations, leading to little uncertainty in summary trees. In addition, due to fluctuating viral prevalence and temporal sampling biases, the number of active lineages varies greatly over time (Suppl. Fig. S2). Complex evolutionary and epidemiological processes likely further contribute to highly imbalanced tree shapes (Fig. 2). These biological properties distinguish viral phylodynamic tree inference from that in phylogenomic analysis of divergent species.

**Fig. 2.**
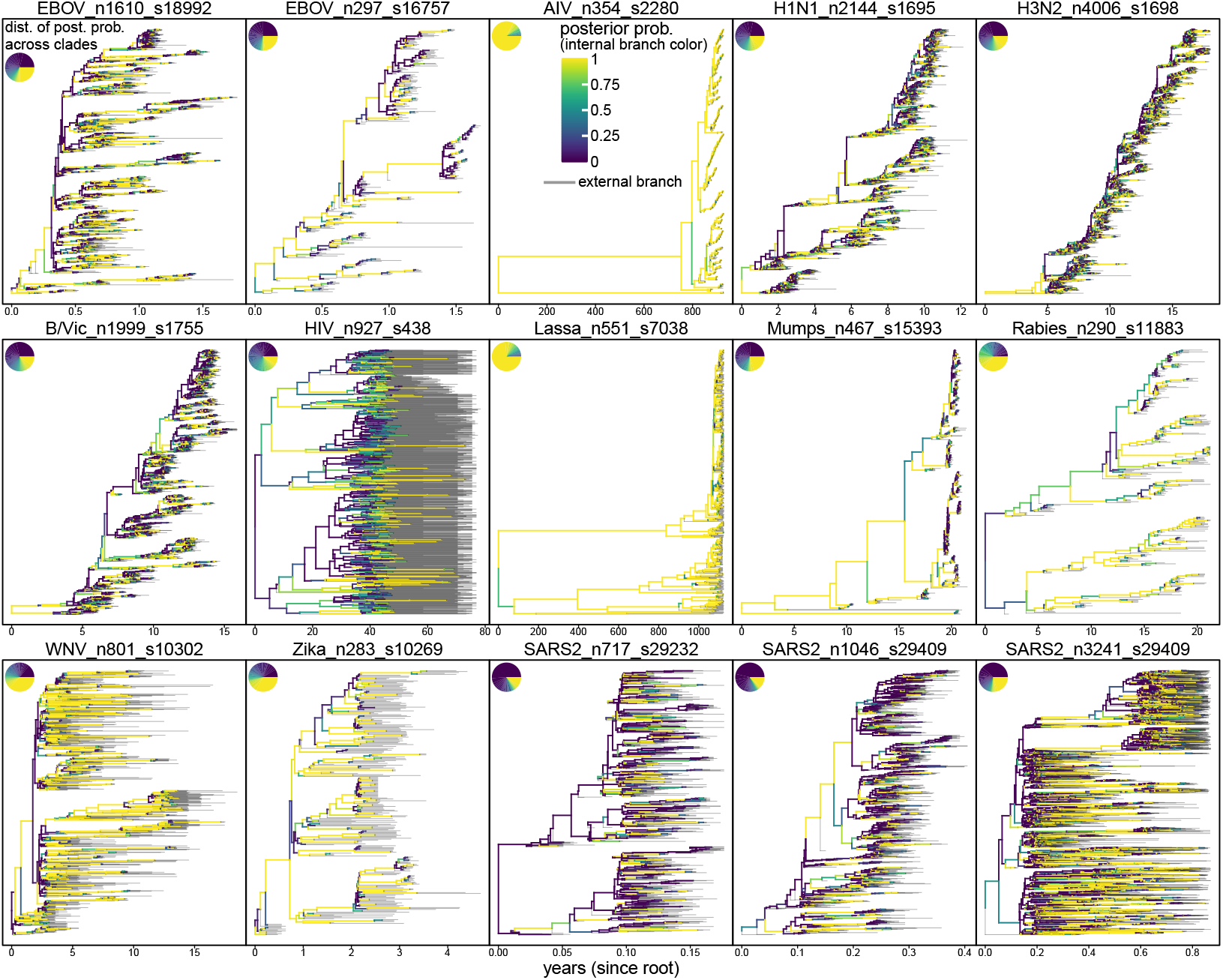
Dated viral phylogenies are typically highly imbalanced and inferred with high uncertainty. Each panel shows the summary MCC (maximum-clade credibility) time tree for a dataset. Branch colors indicate posterior probability of the subtending nodes (yellow: high; purple: low; gray: external branches). Inset pie chart (top-left) summarizes the clade posterior probability distribution in that tree. For nearly all datasets, most clades are weakly supported and subtended by short branches. Trees are highly imbalanced with heterochronous tips unevenly distributed through time. Dataset names at panel tops show the virus, number of sequences (n), and sites (s) in the alignment. Dataset sources (numbered by panel order): 1 (26); 2 (27); 3 (28); 4–6 (29); 7 (30); 8 (31); 9 (32); 10 (33); 11 (34); 12 (35); 13 (36); 14 (37); and 15 (38). See Suppl. Table S1 and Section S2 for dataset details.

**Fig. 3.**
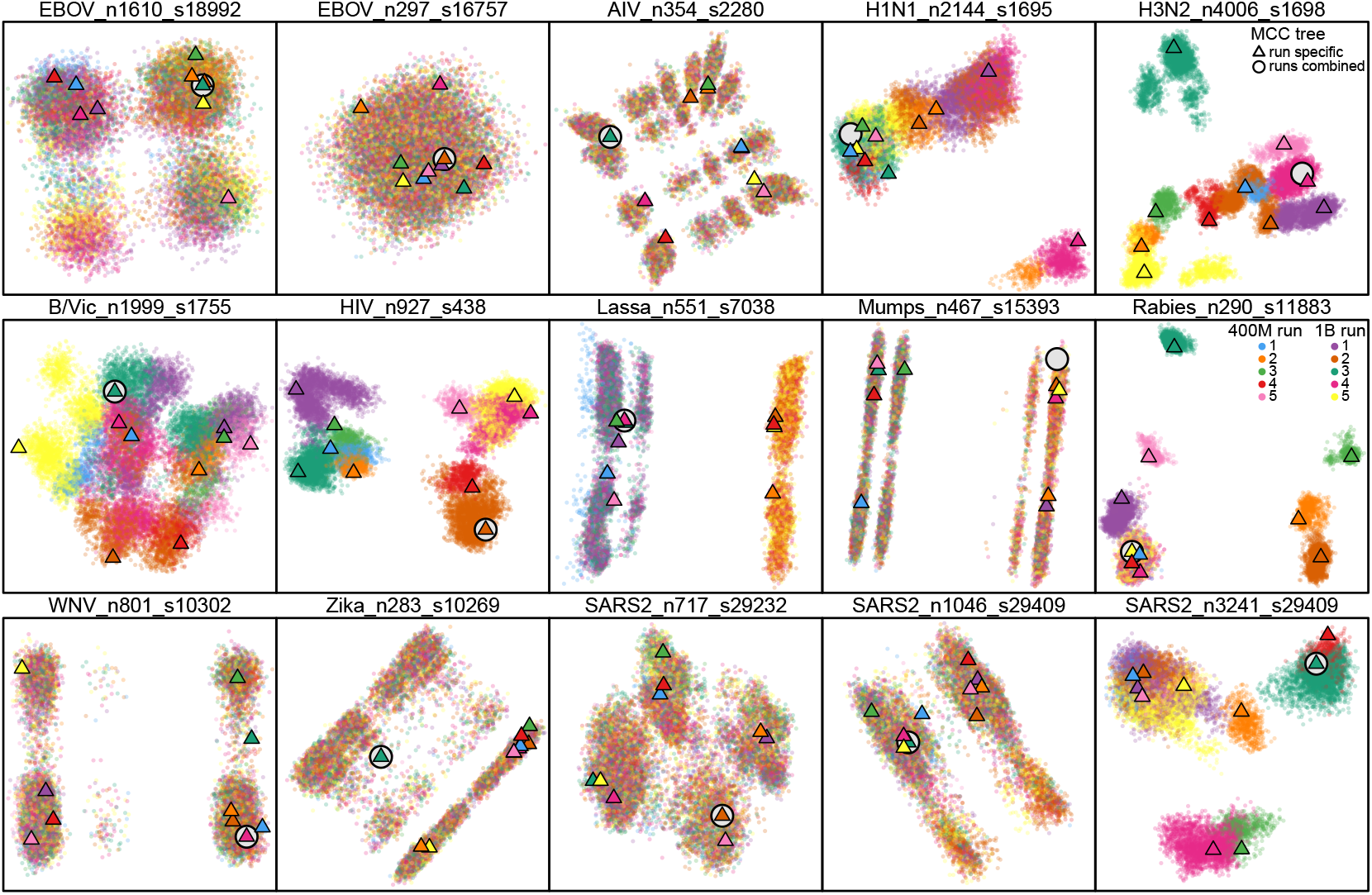
Phylodynamic tree space is frequently multimodal and tree sampling problems are widespread. Each panel shows MDS posterior tree space, based on pairwise RF distances between sampled trees. Each dot represents a sampled tree; color indicates the MCMC chain. A triangle represents the MCC tree of each chain, while an open circle indicates the MCC tree from all chains combined. For most datasets, tree space is multimodal. Seven datasets (*e.g*., HIV) show non-convergence even after one billion iterations, with clear color separation. Five other datasets (*e.g*., EBOV n1610) exhibit slow mixing within each chain, with unevenly mixed colors among peaks.

### Widespread Multimodality and Sampling Problems in Phylodynamic Tree Inference

We visualized posterior tree space via multi-dimensional scaling (MDS), a technique that projects tree space into lower dimensions based on pairwise tree distances (40). Focusing on discrete-topology space, we computed both Robinson-Foulds (RF; 41) and rooted SPR (rSPR; 42) distances between all pairs of sampled trees. For most datasets, posterior tree space was multimodal and sampling problems were widespread; RF-(Fig. 3) and rSPR-based (Suppl. Fig. S4) MDS plots showed qualitatively similar patterns. After months of computation, we obtained exact rSPR distances for only five datasets with smaller trees and less diffuse tree spaces, consistent with theory that rSPR computation difficulty scales with tree space diffuseness (42). We thus focused on results based on the commonly used RF distance.

MCC trees from individual chains resided on different peaks, while the MCC tree from all chains combined typically resided toward MDS plot corners (Fig. 3). In contrast, for some non-converging datasets, the combined summary trees generated by two alternative methods—HIPSTR (43) and CCD0-MAP (44)—lay between peaks (Suppl. Fig. S9). For datasets without convergence failure, chain-specific HIPSTR and CCD0-MAP trees were generally closer to their respective combined summary trees than those from MCC.

Beyond visual inspection, we quantified tree sampling performance via tree-based MCMC diagnostics. We evaluated within-chain mixing with tree ESS (17, 20) and between-chain convergence with tree potential scale reduction factor (tree PSRF; 12, 45) and average standard deviation of clade frequencies (ASDCF; 46). These diagnostics largely corroborated the MDS plots in revealing sampling problems (Suppl. Fig. S5), though showing more non-convergence (12 versus seven) using conventional cutoffs (PSRF above 1.1; ASDCF above 0.01). The non-convergence was primarily identified by ASDCF; raising its cutoff from 0.01 to 0.02 reduced the number of non-converging datasets from 12 to seven, matching the MDS plots.

Such tree-based diagnostics identify sampling problems but are computationally expensive to obtain. Alternatively, MCMC diagnosis of `BEAST` analyses is typically conducted by loading posterior log files into `Tracer` (16) and inspecting ESSs of logged continuous variables. Given this standard practice, we investigated whether diagnostics of variables logged by default, such as posterior density and evolutionary rate, could assess tree sampling performance. We also developed new parsimony-score diagnostics, computed from the minimum number of mutations required over each sampled tree. This score is model-independent, determined solely by topology given the alignment, and insensitive to noise in parameter sampling. While diagnostics of some continuous variables (*e.g*., substitution model parameters and root age) rarely indicated sampling problems, variable diagnostics overall exhibited higher rates of inadequate mixing but lower rates of non-convergence than tree-based diagnostics (Suppl. Fig. S5). In contrast, parsimony-score ESS and PSRF corroborated the MDS plots more closely, indicating inadequate mixing and non-convergence for 12 and seven datasets, respectively.

These datasets exhibited distinct sampling performance. Seven datasets showed non-convergence, with chains stuck in separate regions after billions of iterations. Among them, five (H1N1, H3N2, B/Vic, HIV, SARS2 n3241) exhibited widespread sampling problems (parsimony-score PSRFs > 1.2, ASDCFs > 0.025, parsimony-score ESSs < 100), with HIV showing the most extensive failure and unique problems in sampling substitution model parameters. The other two (Lassa, Rabies) showed adequate within-chain mixing despite non-convergence, suggesting that sampling problems can be concealed without sufficient replicates. Lassa—where chains divided between two peaks, with four chains stuck on the left peak, four on the right, and two that moved between the peaks once—illustrated the difficulty in identifying peaks. In that case, to reduce the chance of missing a second peak (and thus incorrectly assuming convergence) to below 5%, five or more chains would be required.

Three datasets (EBOV _n297, AIV, Zika) showed no sampling problems. The other five (EBOV_n1610, Mumps, WNV, SARS2_n717, SARS2_n1046) exhibited near-convergence yet slow mixing (ASDCFs ∈ (0.01, 0.02), parsimony-score ESSs < 200), with all chains finding the same peaks and commuting between them but requiring tens of millions of iterations per commute, suggesting convergence with longer runs.

### Phylodynamic Posterior Tree Landscapes are Often Operationally Rugged

We overlaid posterior density values of sampled trees on MDS tree space to characterize phylodynamic tree landscapes. As trees from non-converging datasets may not represent the true posterior, we focused on the datasets where chains found the same regions of tree space. Posterior density heatmaps across discretized tree space revealed multiple highdensity peaks separated by low-density valleys (Suppl. Fig. S6). 3D MDS plots with posterior density as the *z*-axis directly depicted isolated peaks at comparable heights (Suppl. Fig. S7). MDS plots retaining only the highest-density samples (1%) highlighted spatial separation of peak tops (Suppl. Fig. S8). These complementary visualizations demonstrate rugged tree landscapes, with multiple similarly high peaks separated by valleys and no clear global mode.

We are focused on characterizing operational ruggedness, which corresponds to how Bayesian phylodynamics is practiced: It depends on how trees are sampled (standard tree moves during MCMC), how tree space is defined (vector (RF) or tree-rearrangement (rSPR) metrics), and how trees are scored (phylodynamic model specification). For example, while RF- and rSPR-based MDS plots are similar overall, they diverge notably for the Lassa dataset. Although both show non-convergence, the large valley between peaks in the RF-based plot (Fig. 4E, Left) is inconspicuous in the rSPR-based plot (Suppl. Fig. S12A). The two peaks are driven by tip 091 wobbling between two positions six nodes apart in the tree (Fig. 4A and B); *i*.*e*., these two trees have an RF distance of 12 but are separated by a single SPR move, explaining the RF and rSPR divergence. Tip 091 is sister to 088 in the right peak; the SPR move crosses the valley from right to left by placing 091 as sister to the cherry (036,046).

**Fig. 4.**
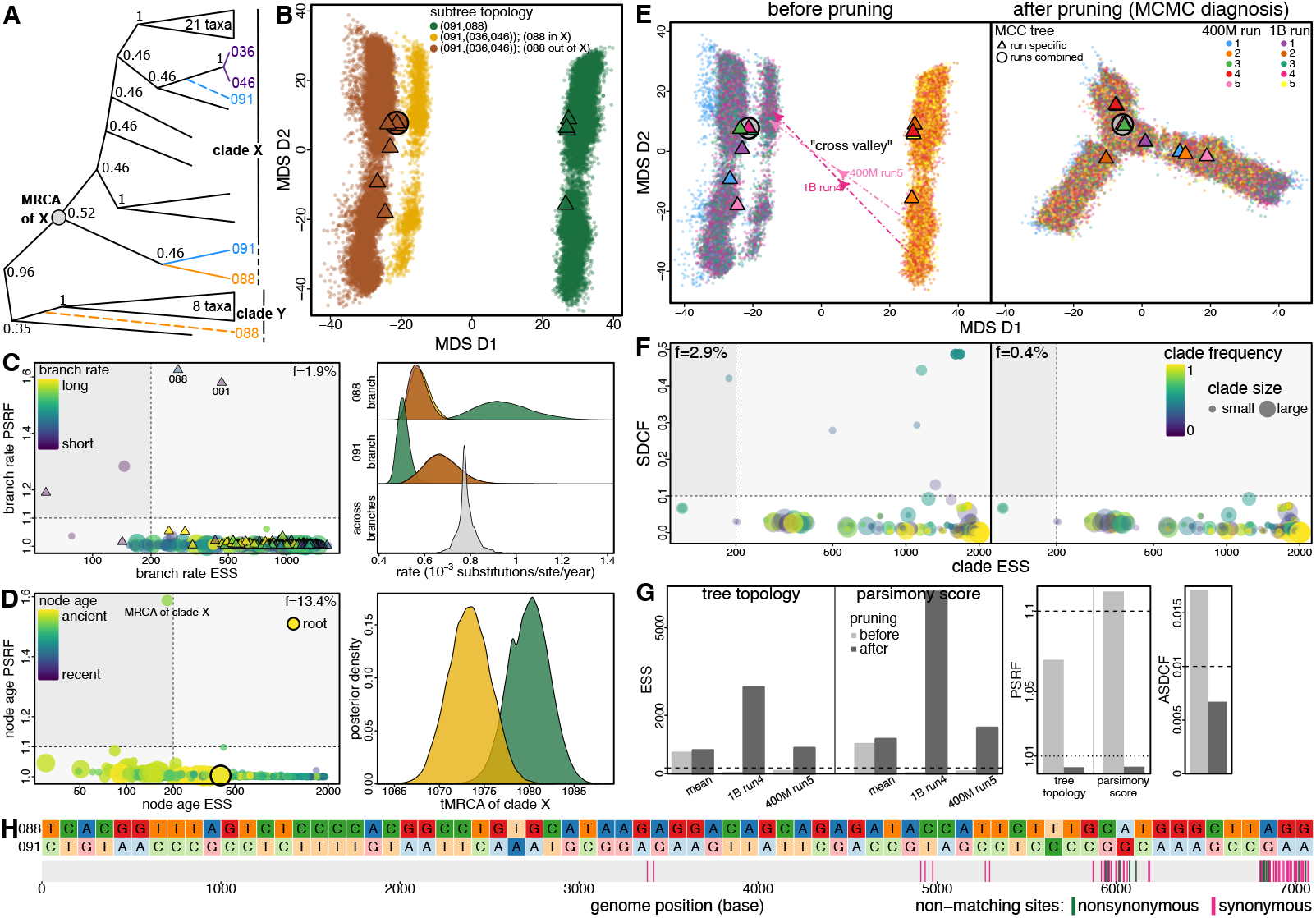
Tree sampling problems can be caused by just a few sequences. **A)** Subtree of the Lassa dataset (31) focusing on the identified problematic sequences (088 and 091). Dashed branches indicate alternative positions of 088 and 091 when they are not sisters. Sequence names (*e.g*., “088”) are shorthands following the original study (47); see Suppl. Table S2 for full names and GenBank IDs. **B)** MDS representation of the posterior tree space (based on RF distance) with each tree colored by the topology of its focal subtree. When 088 and 091 are sisters (green), they are always basal to clade X. When 091 is in turn sister to (036,046) (purple in **A**), 088 alternates between the base of clades X and Y (yellow and brown). **C)** Evolutionary rate estimates for branches subtending these sequences are topology-dependent. (Left) MCMC diagnostics for branch rates. PSRF (*y*-axis) and ESS (*x*-axis) quantify between-chain convergence and within-chain mixing, respectively; dashed lines show standard cutoffs (PSRF>1.1, ESS<200). Circles and triangles represent internal and external branches, respectively, with color indicating rate (yellow: high; purple: low) and size proportional to descendant tip number. A small fraction (*f* = 1.9%) falls outside acceptable ranges (bottom-right quadrant), indicating poorly sampled branch rates. (Right) Posterior distribution of the branch rate (top: 088; middle: 091) conditioning on the topology (colored as in **B**). **D)** tMRCA estimates are also affected by sampling problems. (Left) PSRF of clade X tMRCA reveals convergence failure. (Right) Clade X tMRCA is significantly younger when 088 and 091 are sisters than otherwise. **E)** Posterior tree space before (Left) and after (Right) pruning, with dots colored by chain index (Left panel dots are identical to **B** except for coloring). (Left) Before pruning, at least two peaks are present; the valley is crossed once in only two chains (1B run4 and 400M run5; dashed arrows), with MCC trees of six chains in the left peak and four in the right. (Right) The valley vanishes after pruning, and the combined MCC tree moves near the center. **F)** Clade occurrence diagnostics before (Left) and after (Right) pruning. SDCF (*y*-axis) and clade ESS (*x*-axis) quantify convergence and mixing, respectively. Each dot represents a clade, whose color indicates the mean clade frequency across all chains. **G)** Tree-based diagnostics quantify MCMC performance before (light gray) and after (dark gray) pruning, corroborating visual patterns in **E**. For tree and parsimony-score ESSs (Left), we show the mean over all chains and values from the two chains that crossed the valley. **H)** Incompatible sites between 088 and 091 and their positions in Lassa’s L segment. (Top) Nucleotide states at incompatible sites. Darker cells indicate mutations inferred on the subtending branches of 088 or 091 when they are sisters. (Bottom) Incompatible site positions marked by colored vertical bars (green: nonsynonymous; pink: synonymous).

Nevertheless, we find that the peaks and valley structures are not simply a feature of poor MCMC exploration. For Lassa, although the valley requires just one SPR (or six consecutive NNIs) to cross, it was only crossed twice over five billion iterations. The rare valley crosses result from two factors: (1) intermediate trees along the 6-NNI path between peaks were never sampled, and (2) the direct SPR move rarely succeeded. To validate the valley (factor 1), we performed additional phylodynamic analyses with topology fixed at each step along the 6-NNI path. These analyses revealed that intermediate trees had significantly lower posterior densities than trees at the peaks, with density decreasing when moving away from one peak and increasing after the midpoint (Suppl. Fig. S10A and B). This confirms the peaks-and-valley structure observed in the MCMC-sampled tree landscape.

To understand the rare direct moves between peaks (factor 2), we recorded all SPR moves attempted during the phylodynamic analyses. For Lassa, each specific SPR move was attempted fewer than twice on average per 100 million iterations (Suppl. Fig. S10C), consistent with theory that the attempt frequency of a specific SPR move decreases as the inverse square of tip number. Specifically, the SPR move from right to left was attempted 36 times in all five billion post-burnin iterations with only two acceptances, while the reverse move was attempted 43 times with none accepted.

This low acceptance proportion likely results from distinct evolutionary rates over focal branches under each topology. Branch 088’s rate differed markedly between peaks: the mean rate estimate in the right peak was approximately twice that in the left, decreasing from the 99.5th to 0.5th percentile of all branch rates (Fig. 4C, Right). Notably, the two acceptances (of the 36 direct move attempts from right to left) occurred when branch 088’s rate was near its minimum observed value in the right peak (Suppl. Fig. S10D).

The Mumps dataset provided a key comparison with the Lassa dataset. Like Lassa, the MDS plot revealed two main peaks separated by a single SPR, driven by tip 20.16/2 wobbling between two positions three nodes apart in the tree (Suppl. Fig. S16A and B). Fixed-topology analyses confirmed that intermediate trees had lower posterior densities than trees at peaks (Suppl. Fig. S11A and B), validating peaks-and-valley structure. Unlike Lassa, intermediate trees were occasionally sampled, and valley crosses were more frequent (roughly once per 50 million iterations), avoiding convergence failure but causing inadequate mixing. The more frequent valley crosses compared to Lassa likely result from the shallower valley, shorter NNI path, and higher acceptance proportion (22.3%) of the direct SPR moves. The phylodynamic model specified for Mumps is also simpler (strict-clock rather than relaxed-clock), avoiding tree move rejections due to mismatches in topology-dependent branch rates. This suggests that, in addition to infrequent proposals, low acceptance rates caused by strong parameter correlations under complex phylodynamic models contribute to operational ruggedness.

### Tree Sampling Problems Can Be Dissected by Clade-Specific MCMC Diagnostics

To quantify sampled tree differences and identify clades driving sampling problems, we explored various clade-specific diagnostics (Suppl. Fig. S5). We examined within-chain mixing and between-chain convergence for each clade using clade ESS and standard deviation of clade frequency (SDCF), respectively. Clade ESS is computed by treating each clade’s occurrence as a binary variable: one when the clade is present, zero otherwise (19). For most datasets, fewer than 10% of clades exhibited poor sampling, demonstrating that sampling problems likely arose from a small part of the tree. We also assessed sampling performance of clade-specific continuous variables—including node age, branch duration, and branch rate—by computing their ESSs and PSRFs. Similar to clade occurrence, in most datasets fewer than 20% of clades showed sampling problems for each clade-specific variable. Finally, we evaluated the association of alignment sites with sampling problems by treating each site’s log-likelihood as a continuous variable to compute its ESS and PSRF. For most datasets, a small fraction (below 10%) of sites showed sampling problems (Suppl. Fig. S5), similar to clade-specific patterns. An outlier was the HIV dataset, where most clades and sites exhibited poor sampling, likely caused by its substitution model parameter sampling problems.

### A Few Problematic Sequences Can Strongly Impact Phylodynamic Analysis; MCMC Diagnostics Can Identify Them

Tree sampling problems typically stemmed from a small fraction of sequences that each disrupted multiple clades. For five datasets in which clade-specific diagnostics identified a limited number of problematic sequences, we pruned these tips from each sampled tree and re-assessed MCMC performance by recomputing diagnostics with the pruned trees. Sampling problems were attenuated in all five datasets after pruning, with four requiring only 1–3 tips pruned (Fig. 4 and Suppl. Figs. S15–S18).

As shown for Lassa and Mumps, a problematic tip usually wobbled between topological positions 3–10 nodes apart, creating peaks in tree space with comparable height. Multiple such tips may lead to many peaks combinatorially; *e.g*., the three tips identified in the SARS2 n717 dataset gave rise to at least eight peaks (Suppl. Fig. S15). Such combinatorial explosion—*i*.*e*., 10 problematic tips could create thousands of peaks—may partially explain the widespread sampling problems observed in datasets where each chain found a unique peak. SARS2 n717 also suggested that, with extremely limited genetic diversity, a single recurrent mutation can make the sequence problematic, behaving like a recombinant.

We focus on Lassa to illustrate the impact of sequence removal, where pruning just two of all 551 sequences reveals a unimodal tree landscape and resolves sampling problems (Fig. 4). After removing the two problematic tips, the large valley between peaks and chain separation in tree space both vanish (Fig. 4E). For the two chains that previously crossed the valley once, tree and parsimony-score ESSs both increase by over an order of magnitude after pruning, while tree and parsimony-score PSRFs and ASDCF drop to well below the cutoffs (Fig. 4G). The marked diagnostic improvement corroborates the visual contrast in MDS plots. After pruning, non-converging clades (SDCF above 0.1) no longer exist and the fraction of poorly sampled clades decreases from 2.9% to 0.4% (Fig. 4F). The fraction of sites with sampling problems also decreases from 2.4% to 0.3%, with all site-specific PSRFs dropping below the cutoff (Suppl. Fig. S12D).

To validate the causal role of these sequences, we performed new phylodynamic analyses for Lassa after excluding the two problematic sequences. This prospective removal yields a unimodal tree landscape with satisfactory sampling performance (Suppl. Fig. S13), matching the post-hoc tree pruning results. The sampling diagnostics improve progressively from original analyses through post-hoc pruning to prospective removal. Similar reanalyses of the Rabies dataset, with 14 identified problematic sequences excluded, show comparable improvements in tree-space structure and sampling diagnostics (Suppl. Fig. S17). These results confirm that the identified sequences drive the sampling problems and validate the value of clade-specific diagnostics for identifying such sequences. Thus we find that post-hoc pruning provides an effective MCMC-free way to assess the impact of sequence removal.

### Parameter Estimation Impacts and Biological Causes of Problematic Sequences

Tree sampling problems can strongly impact key biological estimates, especially for parameters associated with nodes and branches near problematic tips. For Lassa, in addition to the previously shown marked differences in branch rates between peaks, the time spans of branches subtending the two problematic tips also appeared topology-dependent, with means varying by 2–4 times and 95% credible intervals (CIs) barely overlapping (Fig. 4C and Suppl. Fig. S12C). The age estimates of nodes (tMRCAs) near those tips were also affected, with the tMRCA of clade X (Fig. 4D) inferred to be significantly younger when 088 and 091 were sisters than otherwise. These strong discrepancies were revealed by the exceptionally high branch-rate PSRFs associated with the two tips and tMRCA PSRF of clade X (Fig. 4C and D). Therefore, branch-rate (and -length) diagnostics are particularly useful for detecting problematic sequences and are more interpretable than clade ESS or SDCF, as one problematic tip can disrupt many clades, rendering them poorly supported.

The prospective removal reanalyses additionally allowed us to validate impacts of problematic sequences on phylodynamic model parameters. For Lassa, similar to the tree sampling diagnostics, removing the two sequences improved parameter sampling, with the fractions of poorly sampled branch lengths, branch rates, and node ages all significantly decreasing (Suppl. Fig. S13). The previously shown topology-dependent rate and age estimates of clades near the wobbling sequences were particularly affected by sequence removal. For example, consider clade Z, the sister to cherry (088,091) in the MCC tree (Fig. 4A). The tMRCA estimates from reanalyses were virtually identical to those from original analyses when 091 was sister to cherry (036,046). However, in the original analyses, the tMRCA was significantly younger when conditioning on the MCC topology where the clade was sister to (088,091).

Problematic sequences can also have global impacts, compromising estimation of tree-wide parameters beyond clade-specific ones. For Rabies, removing the identified problematic sequences dramatically improved sampling performance and estimation reliability of the mean evolutionary rate, from clear convergence failure in original analyses (PSRF > 1.1) to perfectly overlapping posterior distributions among reanalysis chains (Suppl. Fig. S17). However, the converged mean rate estimates after sequence removal were significantly lower than those from any of the non-converging original chains. More generally, as sequence removal alters the data, quantitative parameter differences beyond sampling improvement are expected and can be biologically meaningful.

To understand biological causes of the peaks for Lassa, we examined incompatible sites between the two problematic sequences, locating them in the genome (L segment). The 71 incompatible sites were distributed extremely unevenly along the segment: only two occurred in the first two-thirds 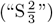 while 69 occurred in the final third 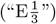, especially the last 5% (Fig. 4H). Furthermore, per-site Hamming distances between 088 and other clade X taxa (excluding 091) in 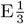 were 3 times those in 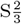 (Suppl. Fig. S14A). `GARD` (48) and `3SEQ` (49) analyses both detected recombination; `GARD` inferred a breakpoint at genomic position 2925, while 342 of the 346 recombinant triplets inferred by `3SEQ` involved 088. In addition, over 10% of sites in 088’s and 091’s genomes were missing (Suppl. Fig. S14B), predominantly at the third codon position. Notably, a sequence from clade Y was used as the reference when computing a consensus sequence for 088 (47). This evidence suggests that these two sequences likely represent consensus sequences from mixed viral samples due to superinfection or contamination.

### Limited Success of Root-to-Tip Regression and Recombination Tests in Identifying Problematic Sequences

We have demonstrated that clade-specific d iagnostics c an identify problematic sequences in datasets where few such sequences exist. However, for seven other datasets in which more clades (though still a small fraction, *≈* 10%) showed poor sampling, no stable backbone topology emerged from sampled trees, precluding identification of key problematic sequences via MCMC diagnostics. In addition, although shorter MCMC chains might suffice for detecting these sequences, hours or days of computation are still required for this preliminary check.

Accordingly, we investigated whether problematic sequences could be identified with simpler and faster data-quality tests rather than full phylodynamic analyses. We conducted root-to-tip (RTT) regression (50) and recombination tests on each dataset, comparing detected outlier and recombinant sequences with those identified by MCMC diagnosis. RTT regression assesses temporal signal and identifies outlier sequences with sampling dates or genetic composition incongruent with the rest. This test revealed strong temporal signal in 11 datasets (Suppl. Fig. S25), nine of which had identified tree sampling problems. RTT detected outliers in all nine datasets, seven of which had more than 10 outliers.

However, despite the large number of RTT outliers, pruning them from sampled trees had little impact on sampling performance (Suppl. Figs. S16–S24), considerably less than the MCMC-diagnosis approach (Suppl. Figs. S16–S18). This reflected minimal overlap between RTT outliers and MCMC-identified problematic sequences. The only exception was the Rabies dataset, where five of seven RTT outliers were also identified by MCMC diagnosis; both approaches improved sampling performance, though the MCMC-diagnosis approach showed more substantial improvement (Suppl. Fig. S17).

To explore whether recombination tests can identify problematic sequences, we analyzed each dataset with `3SEQ`, a quick phylogeny-free method that examines sequence triplets for recombinants. `3SEQ` detected recombination in six datasets, five with identified tree sampling problems. Three datasets showed pervasive recombination; for example, Lassa and Rabies had 346 and 271 putative recombinants, respectively, accounting for 62.8% and 93.4% of all sequences. Additional tests with `GARD`, a likelihood-based phylogenetic method, confirmed strong recombination signal in these two datasets. However, such high percentages of putative recombinants pose challenges for purging them to enable reliable phylodynamic inference. Notably, a small set of sequences typically served as putative parents for most recombinant triplets detected by `3SEQ`. For example, Lassa’s problematic sequence 088 was involved in 98.8% of detected recombinant triplets, and one MCMC-identified problematic Rabies sequence was involved in 72.7%. These results suggest that quick recombination tests may detect key recombinants, making such tests useful preliminary checks in phylodynamic analysis.

### Impact of Tree Sampling on Key Phylodynamic Conclusions

To assess the biological impacts of tree sampling problems, we focused on three phylodynamic inferences: (1) effective population size over time (demographic trajectory), (2) geographic dispersal dynamics, and (3) tMRCAs and evolutionary rates. Many phylodynamic analyses use the skygrid coalescent model (51) to estimate demographic trajectories for inferring outbreak scale and timing. The inferred demographic trajectories were barely distinguishable among chains, except for a few datasets where the curve was shifted slightly (Suppl. Fig. S26). Moreover, the demographic trajectory—including both its accuracy and precision—summarized from the first 10 million post-burnin iterations (2% of the full analysis) was virtually identical to that from the full analysis (Suppl. Fig. S27). Consequently, if demographic trajectory is the primary interest, MCMC chains orders of magnitude shorter than those needed for adequate tree sampling would suffice.

In contrast, geographic phylodynamic estimates can be sensitive to tree sampling problems. For eight datasets with sampling locations, we assessed the impact on the inferred number of dispersal events across all areas and between area pairs by reconstructing a parsimonious dispersal history conditioning on each posterior tree. Although most event numbers were robust to tree sampling problems (Suppl. Fig. S28), event numbers for several biologically important dispersal routes differed significantly among chains (Fig. 5 and Suppl. Figs. S29–S34). For example, the HIV dataset revealed large discrepancies among chains in inferred dispersal events between two focal geographic areas in HIV’s early history (30) (Fig. 5), with median estimates ranging from 3 (400M run4; P(*≤*5 events) > 0.95) to 8 (1B run3; P(*≥* 5 events) > 0.95). Tree sampling problems thus directly led to convergence failure in the inferred dispersal history.

**Fig. 5.**
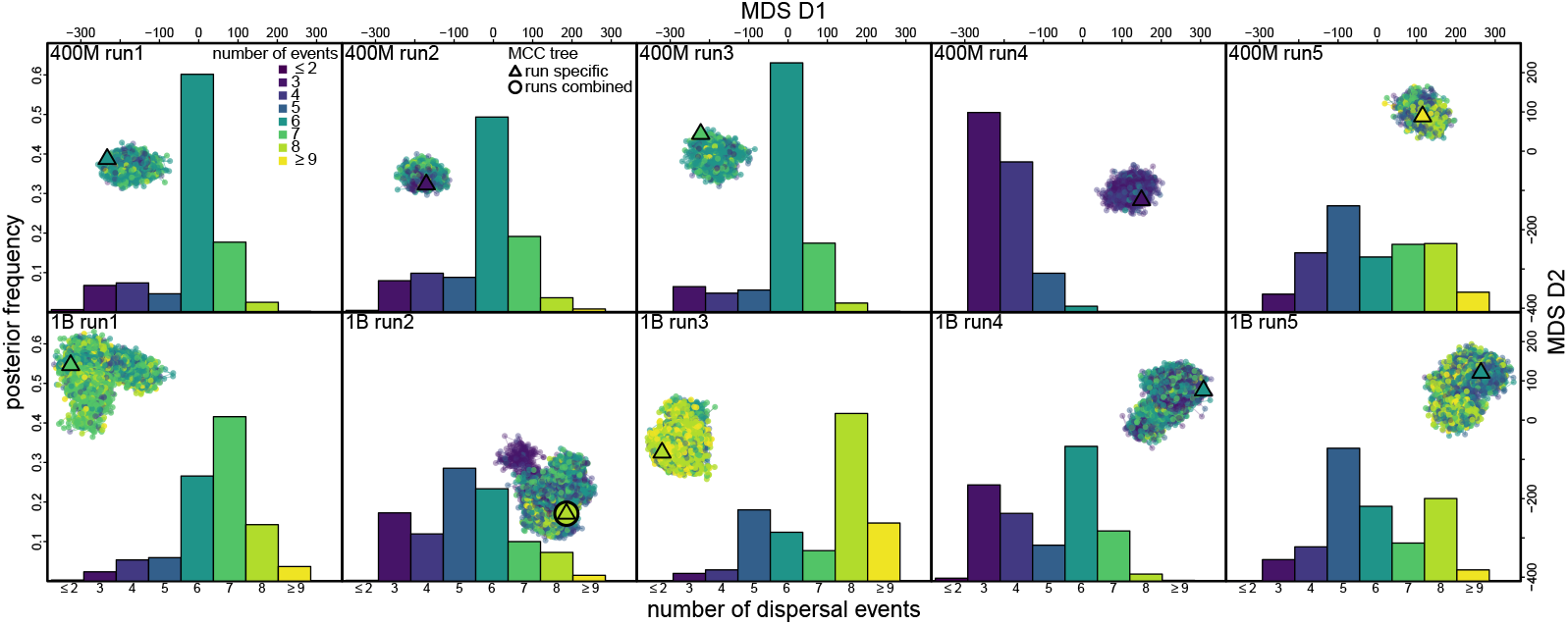
Distinct peaks in tree space can lead to different biological conclusions. Number of dispersal events inferred between two focal geographic areas (Bwamanda and Kinshasa) in HIV’s early history (30). Each panel shows an MCMC chain (top row: 400M runs; bottom row: 1B runs). Each dot represents a posterior tree in MDS tree space, colored by the parsimony number of dispersal events (η) on this route given that tree. Triangles represent the MCC tree of each chain, while the open circle indicates the combined MCC tree. Histograms show the posterior distribution of η for each chain, revealing convergence failure.

For most datasets, root age (Suppl. Fig. S35) and tMRCA of most clades were sampled satisfactorily (Suppl. Fig. S5, Bottom), while mean evolutionary rate estimates differed markedly among chains in five datasets (Suppl. Fig. S37). Both root age and mean evolutionary rate required at least 100 million post-burnin iterations to reach accuracy and precision comparable to the full analysis (Suppl. Figs. S36 and S38), much longer than those required for demographic trajectory. Similar to geographic inference, tMRCA estimates of certain clades can be sensitive to tree sampling. For example, in the HIV dataset, estimates of root age and subtype B origin time differed significantly among chains (Suppl. Fig. S39). In general, sampling problems impact clade-specific estimates (*e.g*., clade tMRCA and geographic source) more strongly than global parameters governed by overall tree shape and lineage fluctuation patterns (*e.g*., demographic trajectory), consistent with the localized origin of tree sampling problems.

## Discussion

Bayesian phylodynamics is often computationally challenging for modern datasets. Recent tools (52, 53) developed for outbreak phylogenetic analysis have focused primarily on accelerating tree evaluation. However, the orthogonal computational challenge of exploring the vast and complex tree space has yet to be assessed. It also has not been clear how well trees must be sampled for reliable inference of focal phylodynamic parameters. Here, to address these key questions for scalable phylodynamics, we comprehensively characterize phylodynamic tree space and tree sampling performance and assess their biological impacts.

We showed that phylodynamic trees are usually markedly imbalanced and inferred with high uncertainty, contrasting with traditional phylogenomic trees. Phylodynamic tree landscapes are diffuse yet often highly rugged, with multiple peaks of comparable height separated by deep valleys, resulting in widespread tree sampling problems. Tree sampling problems were more prevalent than reported for divergent species (18), consistent with their observation that convergence is more difficult with more taxa and shorter branches. We validated the observed valleys through fixed-topology analyses targeting the unsampled trees along the path between the sampled peaks, confirming consistently lower posterior densities at these intermediate trees than at either peak.

Although ruggedness in tree space has been reported (12, 15, 25), our discoveries may seem counterintuitive given the limited genetic diversity in phylodynamic datasets. Limited genetic diversity can indeed drive the ruggedness and sampling problems, as phylogenetic uncertainty greatly inflates tree space (12), inducing numerous terraces (13) and wide flat basins separating local peaks. Moreover, when many closely related sequences differ by at most one nucleotide (*e.g*., SARS-CoV-2 datasets), a single recurrent mutation can create extra peaks that, though small, suffice to cause sampling problems. Our clade-specific diagnostics revealed that tree sampling problems typically arose from a small fraction of clades; sampling of these clades was often disrupted by three or fewer sequences. Removing these problematic sequences retrospectively (by pruning from the sampled trees) or prospectively (via reanalysis after the removal) often dramatically improved sampling performance and eliminated valleys, suggesting the ruggedness may be caused by biological processes (recombination, convergent evolution) or sequencing artifacts. Although previous studies (54, 55) have shown that a few genes can strongly impact tree inference, the undue influence of a few sequences on tree landscape ruggedness and analysis difficulty has not been revealed. We showed that a problematic tip often wobbled between topological positions several nodes apart, while a few such tips can create tens of peaks combinatorially, providing an explanation for the influence.

We also showed that tree sampling problems can conspicuously impact key phylodynamic inferences, with local (clade-specific) parameters more vulnerable than global (tree-wide) parameters. Although demographic trajectory, root age, and most clade tMRCAs were usually robust, estimates of the tMRCA and branch rate associated with certain clades varied significantly between peaks, leading to discrepant mean evolutionary rate estimates in several datasets. Similarly, while the sampling of geographic event counts was generally satisfactory, inferred dispersal history over biologically important routes could be strongly influenced by sampling problems.

Importantly, sampling problems can distort major biological conclusions, especially when focal clades are poorly sampled. For example, for HIV subtype B origin, five chains inferred a time around 1960—supporting the hypothesis that subtype B was seeded by Haitian professionals who worked in the DRC after its independence in 1960 (30)—while the significantly earlier time inferred by the other five chains challenges that hypothesis. Geographic dispersal patterns involving Kinshasa (the putative origin location of HIV group M) also differed markedly among chains, indicating distinct early spread histories of HIV. For Lassa, the non-converging sister relationship between sequences 088 and 091 and the extremely uneven substitution distribution along the genome raise concerns about spuriously identifying human-to-human transmission pairs and thus missing key zoonotic spillover events. More generally, poor tree sampling may mislead transmission tracing and epidemiological hypothesis testing that depend on accurate inference of phylogenetic relationships.

Our results confirmed that traditional and tree-based MCMC diagnostics are useful for quantifying sampling performance in phylodynamic inference but require care with cutoffs. MCMC diagnostics appeared more conservative than the MDS plot in revealing sampling problems, potentially due to overly lenient rule-of-thumb cutoffs (19, 20). Moreover, tree space diffuseness may undermine tree diagnostic power, as tree distances saturate quickly across MCMC iterations, yielding misleadingly small autocorrelation (*i*.*e*., high ESS) and indistinguishable within- and between-chain variations (*i*.*e*., low PSRF). Conversely, ASDCF indicated more pervasive non-convergence than the MDS plot or tree PSRF (consistent with a prior report; 18), though ASDCF’s traditional cutoff (46) may not apply when most clades are weakly supported (see also 19). Thus, we advocate complementing automated diagnostics with visual inspection of the tree space and landscape.

Unlike continuous variable distributions, tree space is metric-dependent (11). As shown here and reported previously (12, 56), although different metrics usually yield qualitatively similar tree space structures for identifying sampling problems, peak number and valley sizes may vary across metrics. These discrepancies can indeed provide information on the cause of tree sampling problems. For example, as shown for Lassa, a valley that is prominent under RF but inconspicuous under rSPR suggests a single clade with multiple topological positions that are similarly supported but far apart. Therefore, we recommend visualizing tree space under multiple metrics. However, exact tree-rearrangement distance computation is often unfeasible for modern phylodynamic datasets; approximate polynomial algorithms (57–59) thus warrant further investigation. Deep-learning methods (*e.g*., PhyloVAE; 60) that directly infer and visualize a representation of latent tree space, bypassing both predefined tree metrics and dimension-reduction steps, also deserve more exploration.

The rugged posterior is also difficult to summarize with a single tree. We showed that two recently developed summary methods, HIPSTR (43) and CCD0-MAP (44), provide qualitatively different summaries than the traditional MCC approach. For example, when chains fail to converge due to isolated peaks, HIPSTR and CCD0-MAP sometimes produced summary trees from between-peak regions that were never sampled. It is unclear whether such intermediate trees represent good summaries across peaks missed by sampling, or if they are actually disfavored by the posterior despite having high conditional clade probabilities. However, both HIPSTR and CCD0-MAP are based on conditional clade distribution (CCD; 61, 62), which has been shown to systematically overestimate the probability of between-peak trees and underestimate those within peaks (12).

Continuous variable diagnostics revealed distinct sampling performance across parameters. While diagnostics for root age, population size, and substitution model parameters almost never exhibited poor sampling, the least satisfactory variable diagnostics (usually for clock model parameters, tree length, or posterior density) often showed worse mixing than the tree, suggesting additional problems sampling rates and times. This confirmed the limited power of existing variable diagnostics for assessing tree sampling (20, 21).

Here we proposed new diagnostics based on the classical parsimony score and demonstrated their effectiveness in identifying tree sampling problems. These diagnostics offer key advantages: they are model-independent, robust to parameter sampling noise, and avoid expensive tree comparisons. We recommend adding them to the standard diagnostic set especially when the topology is a focal parameter.

The distinct sampling efficiencies across parameters and topology suggest room for adapting MCMC settings to specific phylodynamic questions. For example, while billion-iteration runs often inadequately sampled the tree, just 10 million (post-burnin) iterations usually sufficed for reliable demographic trajectory estimates. In theory, many distinct trees may map to nearly identical values for a given parameter, rendering unsatisfactory tree sampling inconsequential for that parameter. Therefore, rather than only checking the ESSs of variables logged by default, or going to the other extreme, exhaustively examining all possible diagnostics, we suggest focusing on diagnosing parameters underlying biological conclusions. When certain topological relationships or clades underlie the focal questions, we recommend applying corresponding clade-specific diagnostics.

When focal estimates are affected by sampling problems, removing the problematic sequences offers a potential solution. Our clade-specific diagnostics proved useful for identifying problematic sequences, but they require costly phylodynamic analyses and work best when few sequences are problematic. Quick prospective detection of problematic sequences remains an open challenge, as data-quality tests show limited success.

The weak association between outliers detected by these tests and MCMC-identified problematic sequences may reflect the heterogeneous origins of these sequences. Some of these sequences may be true recombinants, making recombination tests (*e.g*., `3SEQ`) useful, while others may result from meta-data (sequence and sampling time) mismatches, potentially detectable by RTT regression. Sequencing errors may create other problematic sequences that are hard to detect without raw data. These factors are increasingly concerning as phylodynamic studies now routinely analyze all available pathogen genomes, sporadically collected across outbreaks and sequenced with varying data quality (6, 63).

Rather than simply removing all detected outliers, we recommend re-examining raw data—such as raw reads and sequencing pipelines—for potential errors. If no error is found, or if the outliers are of particular biological interest, we suggest running phylodynamic analyses with and without them to ensure conclusion robustness. If problematic sequences are identified only post-analysis via clade-specific diagnostics, studies can follow our approach to confirm their causal role by pruning them from the sampled trees and re-assessing sampling performance. Once confirmed, rerun phylodynamic analysis with the sequences corrected (or removed), using the last (pruned) tree as the starting tree. When resources allow, we suggest running many (*e.g*., 10) checkpointed (64) chains concurrently and diagnosing them before running too long (*i*.*e*., early post-burnin). If the sequences cause only sampling problems but are not erroneous or recombinant, studies can run additional analyses to re-infer phylodynamic parameters marginalizing over the pruned trees (30, 65, 66).

Some tips that are supported at multiple topological positions may appear problematic only due to the difficulty of moving between the positions. These tips may be viewed as “rogues” (67)—taxa with varying positions across similarly good trees—and many methods (68–70) focus on pruning them from the inferred tree set for better consensus trees. However, their impact on tree search remains unexplored.

Investigating the paths and conditions that enable occasional peak switches could reveal how to increase their frequency, benefiting not only Bayesian but also maximum-parsimony and maximum-likelihood methods that risk getting trapped at local peaks. The peaks are often separated by a single SPR move, but simple proposals chosen uniformly among millions of candidate branch pairs make specific move attempts exceedingly rare. Alternative proposal schemes—*e.g*., extended (46), guided (61, 71), and adaptive (72) tree moves— may increase peak-switch proposal frequency. However, even when proposed, the moves typically have very low acceptance rates despite comparable posterior densities among peaks. Commonly used time tree moves alter topology (and the duration of affected branches as a side effect) but not the branch rates, likely causing rejections due to mismatches between rates and times. Moves that jointly change topology and branch rates to preserve the product of rate and time on affected branches (73–75) could improve acceptance rates and deserve further development.

More generally, understanding the operational nature of tree landscape ruggedness—how it arises from the interplay between biological features of phylodynamic datasets, complex parameter space under phylodynamic models, and routine application of standard time tree moves—is essential for developing solutions navigating the complex tree space in phylodynamic inference. Like the classic DS1–DS11 datasets (46), our compiled datasets and the large number of posterior samples (available in our supplementary repositories) will advance benchmarking and training of new phylodynamic algorithms implementing the solutions.

Our study highlights the need to develop and adopt best practices in phylodynamic analysis, and identifies methodological advances towards scalable phylodynamics. By comprehensively characterizing the operational ruggedness of phylodynamic tree landscapes and dissecting sampling problems, we provide actionable strategies for efficient phylodynamic inference. We are optimistic that Bayesian phylodynamics— with continuing advances in rigorous analysis pipelines and robust computational tools—will be ever more instrumental in pathogen biology research and infectious disease control.

## Materials and Methods

We briefly summarize the methods and analyses here; full details and supplementary results are in SI Appendix.

### Data Curation and Phylodynamic Analyses

To compile the datasets (except for the SARS-CoV-2 ones), we obtained sequence alignments and sampling times from the `BEAST` XML files provided in the original studies. For the SARS-CoV-2 datasets, we acquired sequences from GISAID (76) according to their IDs. We performed Bayesian phylodynamic analyses of each dataset by running 10 independent MCMC replicate chains using `BEAST` v1.10.5 (39) with `BEAGLE` v3.2.0 (77) enabled, including five 400-million-iteration chains to reflect best practices in most empirical studies and five 1-billion-iteration chains aimed at better approximating the posterior distribution. We recorded all tree moves attempted during MCMC, including the branches involved at each move and whether the move was accepted. We specified models, priors, and MCMC proposals and their weights following the original studies; the proposal settings were mostly identical to `BEAUti` v1.10.5 defaults under the specified models. We discarded a conservatively long burnin (the first 200 million iterations) from each chain to focus on the post-burnin behavior, and sampled every 100 thousand iterations.

### Tree Space Characterization and MCMC Diagnosis

We characterized MCMC performance and explored biological impacts by examining three types of variables: (1) continuous model parameters (*e.g*., evolutionary rate), statistics (*e.g*., tree length), and probability densities (*e.g*., joint posterior); (2) clade-specific (*e.g*., node age) and site-specific (*e.g*., site likelihood) variables; and (3) the tree. For each continuous variable, we assessed within-chain mixing by effective sample size (ESS), computed for each chain (using post-burnin samples) with an R (78) implementation of Tracer’s (16) ESS calculation routine via the essTracer function in the convenience package (19). To report mean ESS across all chains for each variable, we truncated the five 1B runs so that all chains had identical length. We assessed between-chain convergence by potential scale reduction factor (PSRF; 45), computed with the Rhat function (79) in the rstan package (80).

We used `TreeSummary` (a BEAST tool) to post-process posterior trees and generate clade-specific variable samples. To compute alignment-wise and site-specific parsimony scores given each sampled tree, we used the `parsimony` function in the `phangorn` package (81). We computed branch-specific parsimony scores by first calling `phangorn`‘s `ancestral.pars` function to reconstruct the parsimonious mutation history for each site given the tree, then counting mutations across all sites for each branch. Similarly, we obtained the total number of parsimony dispersal events over each tree using parsimony. The number of pairwise parsimony dispersal events was computed by first reconstructing the parsimonious geographic history given each tree using `ancestral.pars`, then counting events between each pair of areas.

We characterized the tree space visually and quantitatively with three types of summaries: (1) the MCC summary tree, (2) a two-dimensional MDS plot, and (3) tree-based MCMC diagnostics. For each dataset, we summarized an MCC tree from each chain and the MCC tree from all chains combined using `TreeAnnotator` v1.10.5 (43). We also generated summary trees using two alternative methods—HIPSTR (43) and CCD0-MAP (44)—implemented in `TreeAnnotator` v1.10.5 and v2.7.8 (82), respectively. We calculated pairwise RF distances between rooted trees by counting incompatible clades (instead of splits) obtained via ape’s (83) `prop.part` function, and pairwise rSPR distances using rspr (42, 57). We obtained exact rSPR distances for five smaller datasets after months of computation, while computing approximate rSPR distances for the other 10 datasets. We computed ASDCF by first obtaining the SDCF of each clade, calculated as the standard deviation of estimated clade frequencies among all chains, then taking the mean over all clades (46).

### Visualizing Rugged Tree Landscapes and Assessing Problematic Sequence Impact

To visualize tree landscape ruggedness, we overlaid the posterior density value associated with each sampled tree on the MDS plot. To validate observed valleys, for two datasets (Lassa and Mumps), we performed additional phylodynamic analyses with topology fixed at each position along the valley-crossing path—while other settings follow the original analyses but with fewer and shorter chains as the sampling was much easier with fixed topology—and compared posterior densities across these fixed topologies. To assess the impact of problematic sequences identified by clade-specific diagnostics, we used `TreePruner` (a `BEAST` tool) to prune the problematic tips from each sampled tree and recomputed MCMC diagnostics with the pruned trees. In addition, we reanalyzed two datasets (Lassa and Rabies) after prospectively removing the problematic sequences from the alignments (with other settings following the original 400M runs), and compared the tree space structure, MCMC performance, and phylodynamic parameter estimates between reanalyses and original analyses.

To perform RTT regression (50), we first conducted maximum-likelihood (ML) phylogenetic inference of each dataset using IQ-TREE v2.3.2 (84). We then rooted inferred ML trees by minimizing the root mean squared residuals using `ape`‘s rtt function. Finally, we conducted linear regression between estimated RTT genetic distances and sampling times using `R`‘s `lm` function. We analyzed each dataset with two recombination detection methods, `GARD` (48) and `3SEQ` (49). `GARD` analyses were run with HyPhy v2.5 (85) under default settings. We manually parallelized `3SEQ` analyses so that sequences could be evaluated concurrently, demonstrating the practicality of such tests as quick preliminary checks.

## Data, Materials, and Software Availability

All data and code necessary to reproduce our results are available in the GitHub repository (https://github.com/jsigao/ssstreesupparchive) and the Zenodo repository (https://doi.org/10.5281/zenodo.15574519). The posterior samples are also provided in the repositories; the posterior outputs (*e.g*., posterior trees) excluded from or subsampled in the GitHub repository due to size limit are available in the Zenodo repository.

## ACKNOWLEDGMENTS

We thank Andrew Magee for valuable discussions. This project was partially supported by US National Institutes of Health grants R01 AI162611 and R01 AI153044. Scientific Computing Infrastructure at Fred Hutch was funded by ORIP grant S10OD028685. Dr. Matsen is an Investigator of the Howard Hughes Medical Institute. G.B. acknowledges support from the Research Foundation - Flanders (“Fonds voor Wetenschappelijk Onderzoek - Vlaanderen,” G0E1420N, G098321N), from the European Union Horizon 2023 RIA project LEAPS (grant agreement no. 101094685), and from the DURABLE EU4Health project 02/2023-01/2027, which is co-funded by the European Union (call EU4H-2021-PJ4) under Grant Agreement No. 101102733.

