## Supplementary Material for "Biological causes and impacts of rugged tree landscapes in phylodynamic inference"

### Supplementary Information for: Biological Causes and Impacts of Rugged Tree Landscapes in Phylodynamic Inference

#### Contents

**S1 Supplementary Figures for the Main Text** **S2**
S1.1 Biological Properties of Phylodynamic Datasets Underlie Tree Inference Difficulties . . . . . . S2
S1.2 Widespread Multimodality and Sampling Problems in Phylodynamic Tree Inference . . . . . S4
S1.3 Phylodynamic Posterior Tree Landscapes are Often Operationally Rugged . . . . . . . . . . . S6
S1.4 Identifying Problematic Sequences and Assessing Their Impacts on Phylogenetic Inferences . S12
S1.5 Impact of Tree Sampling on Key Phylodynamic Conclusions . . . . . . . . . . . . . . . . . . . S26

**S2 Analyses of Empirical Datasets** **S36**
S2.1 General Analysis Protocol . . . . . . . . . . . . . . . . . . . . . . . . . . . . . . . . . . . . S36
S2.1.1 Data Curation . . . . . . . . . . . . . . . . . . . . . . . . . . . . . . . . . . . . . . S36
S2.1.2 Bayesian Phylodynamic Analysis Setup . . . . . . . . . . . . . . . . . . . . . . . . . S37
S2.1.3 Analysis Post-processing and MCMC Diagnosis . . . . . . . . . . . . . . . . . . . . . S37
S2.2 Data and Code Availability . . . . . . . . . . . . . . . . . . . . . . . . . . . . . . . . . . . S42
S2.3 Expanded Meta Summaries of Empirical Analyses . . . . . . . . . . . . . . . . . . . . . . . . S43
S2.4 Expanded Dataset-Specific Summaries of Empirical Analyses . . . . . . . . . . . . . . . . . . S52
S2.4.1 Ebola Virus West African Outbreak Dataset . . . . . . . . . . . . . . . . . . . . . . . S52
S2.4.2 Ebola Virus DRC relapse Dataset . . . . . . . . . . . . . . . . . . . . . . . . . . . . . S53
S2.4.3 Human Influenza Virus Datasets . . . . . . . . . . . . . . . . . . . . . . . . . . . . . S54
S2.4.4 Avian Influenza Virus Dataset . . . . . . . . . . . . . . . . . . . . . . . . . . . . . . S56
S2.4.5 HIV Dataset . . . . . . . . . . . . . . . . . . . . . . . . . . . . . . . . . . . . . . . . S57
S2.4.6 Lassa Virus Dataset . . . . . . . . . . . . . . . . . . . . . . . . . . . . . . . . . . . . S58
S2.4.7 Mumps Virus Dataset . . . . . . . . . . . . . . . . . . . . . . . . . . . . . . . . . . . S60
S2.4.8 Rabies Virus Dataset . . . . . . . . . . . . . . . . . . . . . . . . . . . . . . . . . . . S62
S2.4.9 West Nile Virus (WNV) Dataset . . . . . . . . . . . . . . . . . . . . . . . . . . . . . . S64
S2.4.10 Zika Virus Dataset . . . . . . . . . . . . . . . . . . . . . . . . . . . . . . . . . . . . S66
S2.4.11 SARS-CoV-2 Origin Dataset . . . . . . . . . . . . . . . . . . . . . . . . . . . . . . . S67
S2.4.12 SARS-CoV-2 Brazil Dataset . . . . . . . . . . . . . . . . . . . . . . . . . . . . . . . S69
S2.4.13 SARS-CoV-2 Europe Dataset . . . . . . . . . . . . . . . . . . . . . . . . . . . . . . S70

### S1 Supplementary Figures for the Main Text

#### S1.1 Biological Properties of Phylodynamic Datasets Underlie Tree Inference Difficulties

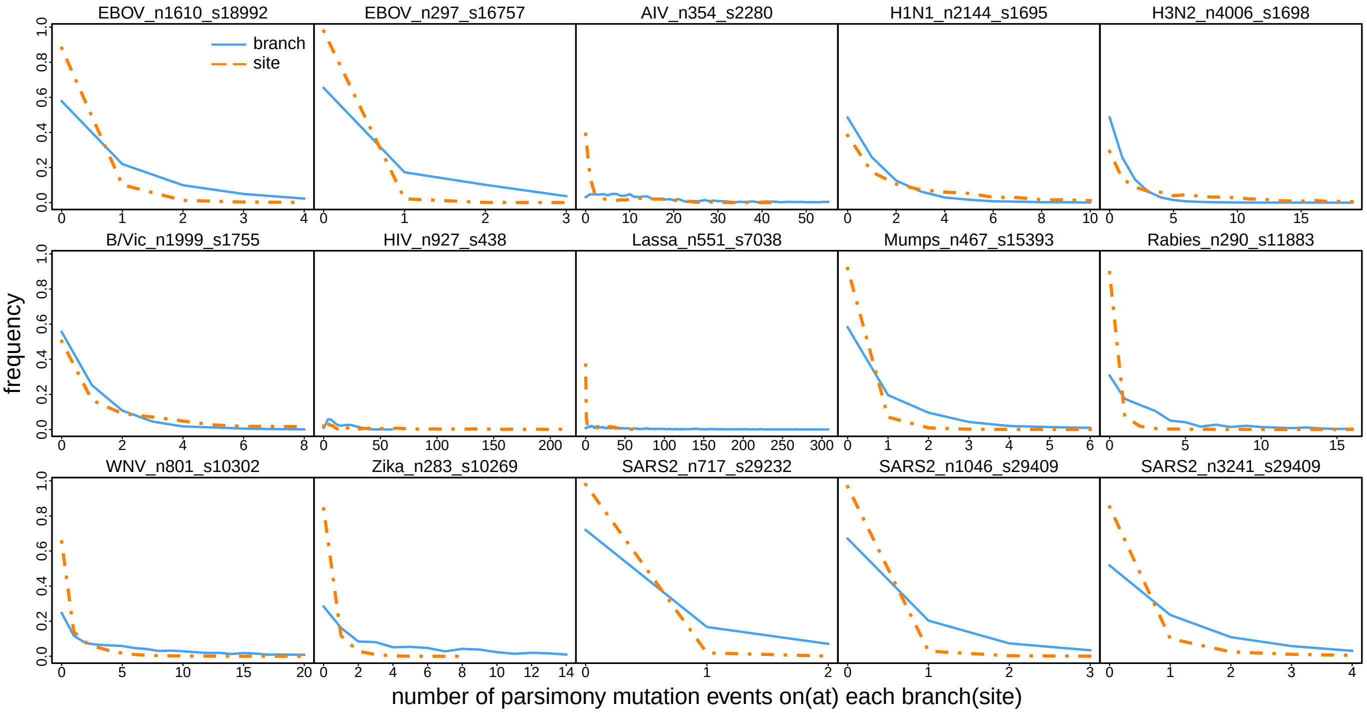


**Figure S1: Limited genetic diversity among densely sampled viral sequences gives rise to limited phylogenetic signal.** Empirical probability mass of the parsimony number of mutations on each branch across the alignment (solid blue) or at each site over the tree (dashed orange). Each panel corresponds to an empirical dataset. For most datasets, the probability mass concentrates on zero or one mutation on each branch and at each site.

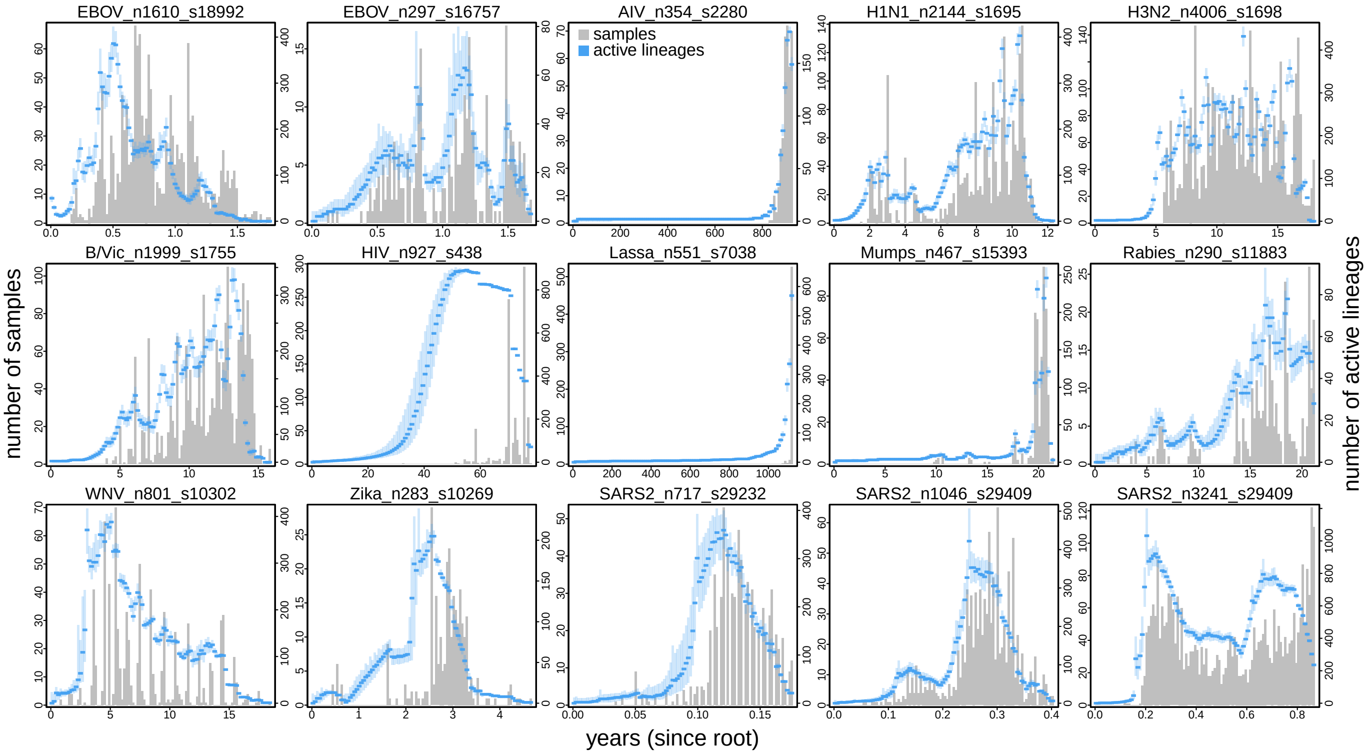


**Figure S2: Viral sequences are sampled unevenly and the number of active phylogenetic lineages fluctuates heavily through time.** The histogram (gray) shows the number of viral samples in each time window from the root (leftmost) to the most recently sampled tip (rightmost), while the trajectory (blue) indicates the inferred number of active lineages (solid line: mean; shaded region: 95% credible interval) in the corresponding time window summarized across the posterior trees.

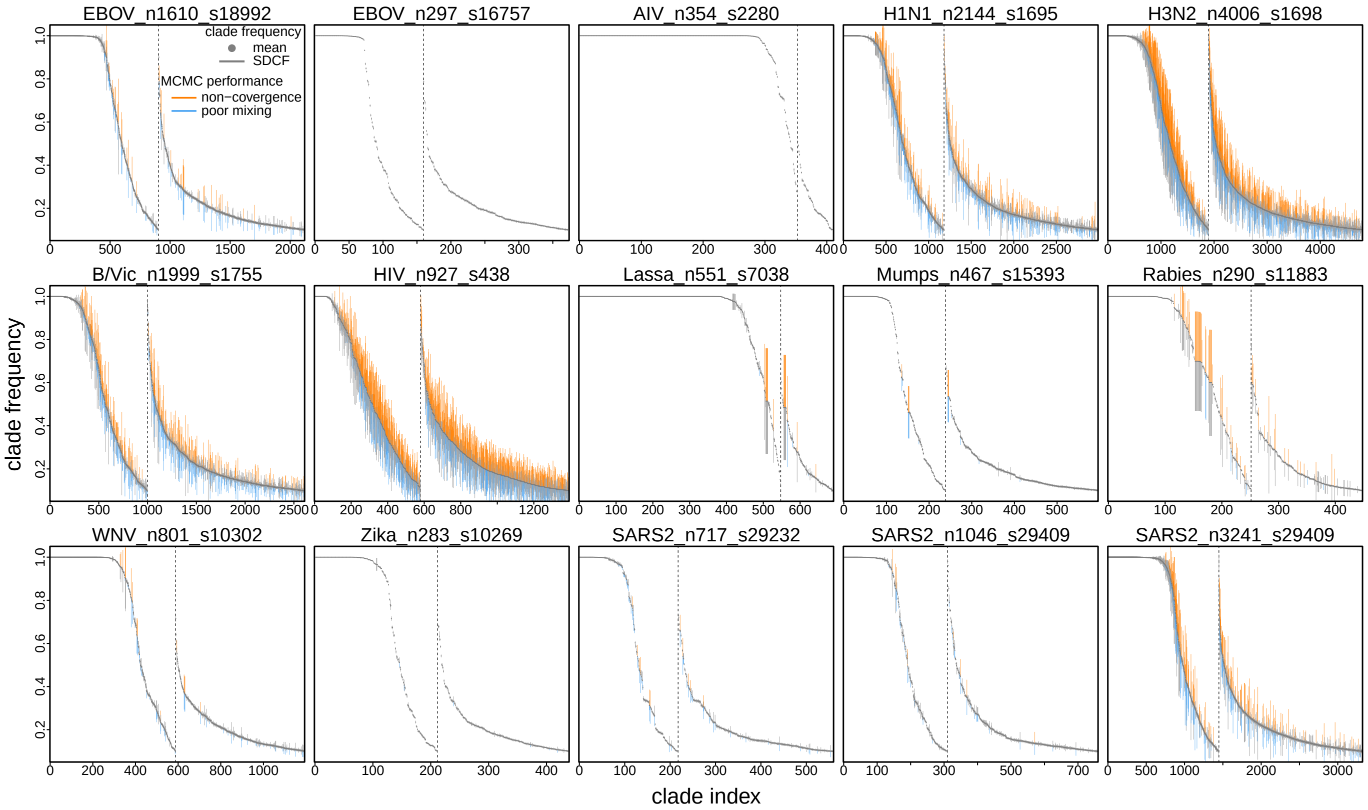


**Figure S3: Most clades in the posterior trees are weakly supported, including those in the MCC tree.** In each panel, a dot represents a clade with the $y$-axis indicating its inferred mean posterior probability across all chains (those clades whose posterior probability is below 0.1 are excluded); the clades are ordered by the posterior probability decreasingly from left to right. The clades that appear in the MCC tree are placed to the left of the vertical dashed line while those absent from the MCC tree are placed to the right of that line. Each dot is plotted with an associated vertical bar centered on it with the bar length indicating the standard deviation of clade frequency (SDCF) of that clade; the upper half of the bar is colored orange to indicate lack of convergence between chains (*i.e.*, SDCF above 0.1), while the lower half is colored blue to indicate poor mixing (*i.e.*, ESS below 200). Across all datasets, most clades are weakly supported, even those in the MCC tree. Except for the human influenza virus (H1N1, H3N2, and B/Vic) and HIV datasets, a relatively small fraction of the clades exhibit sampling problems.

#### S1.2 Widespread Multimodality and Sampling Problems in Phylodynamic Tree Inference

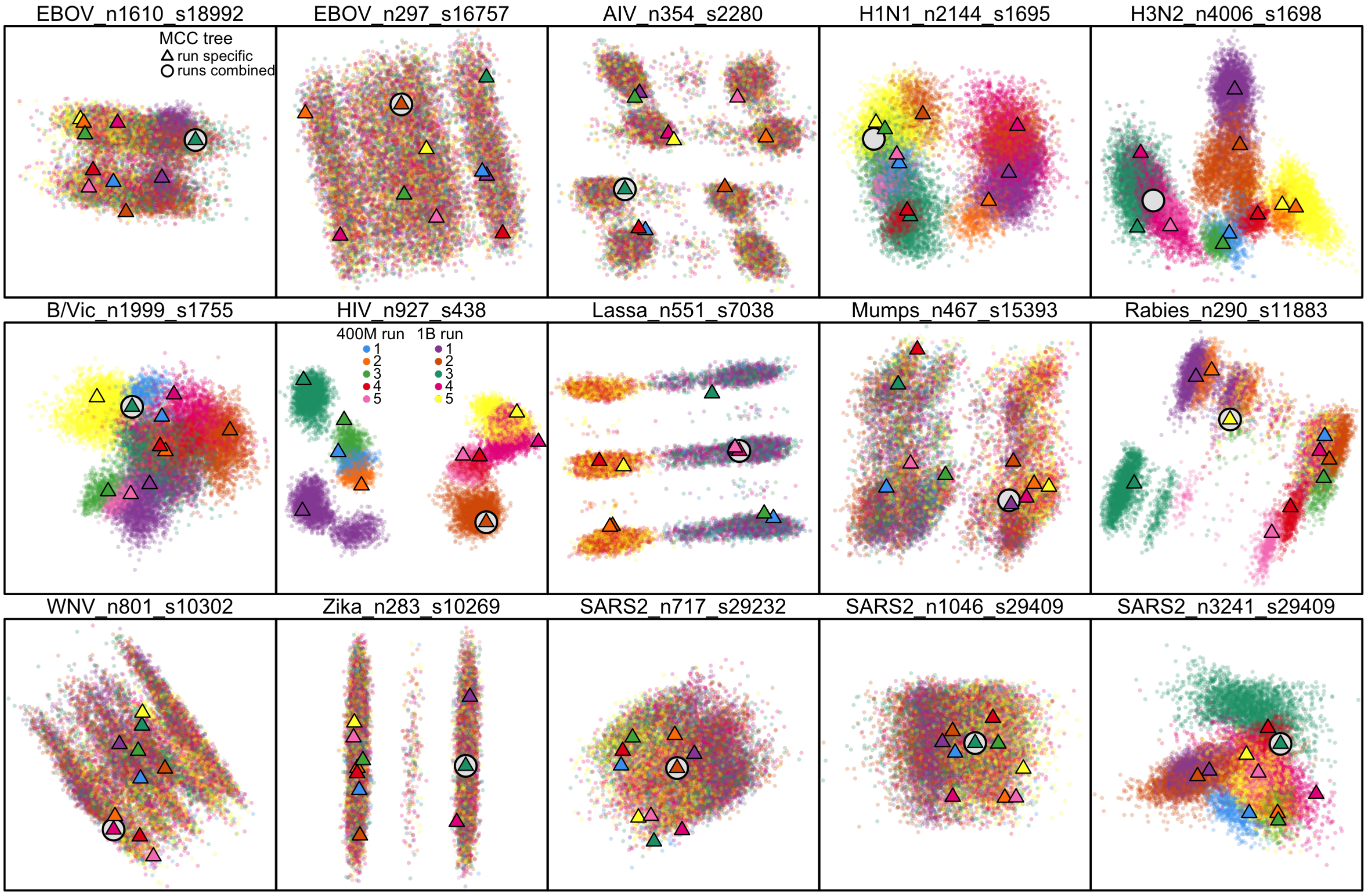


**Figure S4: Phylodynamic tree space is diffuse, and tree sampling problems are widespread.** This figure is generated similarly to Fig. 3 other than that the MDS coordinates are calculated based on the pairwise rooted subtree-prune-and-regraft (rSPR) distances (1, 2) instead of RF distances. For five datasets, including EBOV_n297_s16757, AIV_n354_s2280, Lassa_n551_s7038, Rabies_n290_s11883, and Zika_n283_s10269, we were able to obtain the exact rSPR distances; for the other 10 datasets where the rSPR calculation becomes computationally impractical due to the tree size, we used the approximate rSPR instead. These rSPR tree space plots appear qualitatively similar to their RF counterparts, corroborating Fig. 3 in showing the multiple local peaks. As the impact of using the approximate instead of exact rSPR distances in generating the tree space MDS remains unclear, those approximate-rSPR-based MDS plots should be interpreted with caution and thus we focus on presenting and interpreting the RF-distance based results in our main text. Each panel shows the MDS plot visualizing the posterior tree space of a dataset. Each dot represents a tree sampled from the posterior with its color indicating the corresponding MCMC replicate chain index. A triangle represents the MCC tree summarized from each MCMC chain, while an open circle indicates the MCC tree summarized across all the replicates of that dataset. For most datasets, the posterior tree space appears to be multimodal. Some of these datasets (*e.g.*, Mumps_n467_s15393) exhibit a pattern of slow mixing, as their peaks appear to be commutable albeit with a large number of iterations required for each commute. For some other datasets (*e.g.*, H1N1_n2144_s1695), however, independent chains fail to converge (to the same region in tree space), indicated by the clear separation of colors; *i.e.*, for these datasets, each individual MCMC is stuck at a local peak and is unable to commute between the peaks even after a billion iterations.

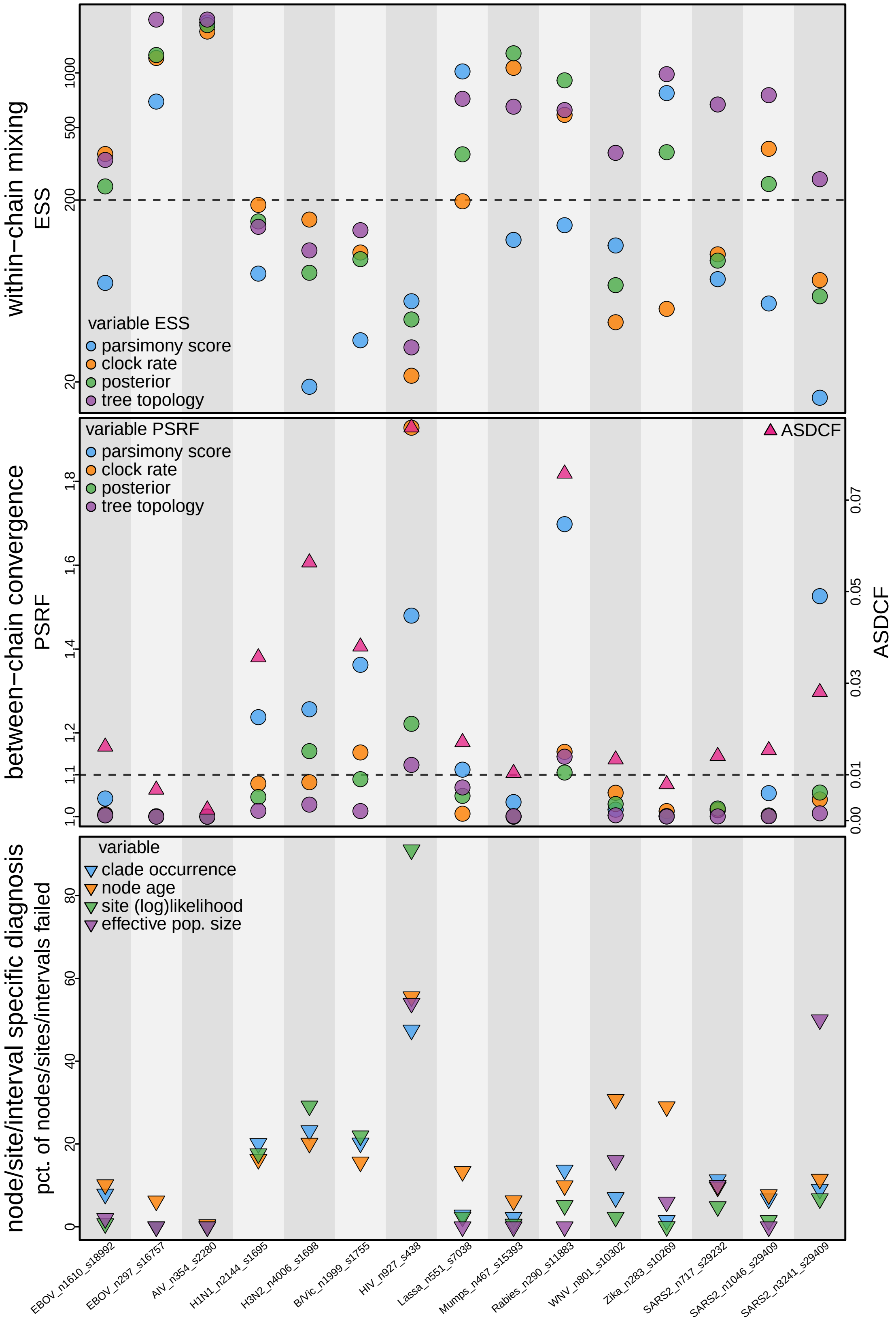


**Figure S5: MCMC diagnostics reveal tree sampling problems, corroborating the visual pattern shown in Fig. 3.** Each column, indicated by the alternating gray/white background, corresponds to a dataset. **(Top)** ESS diagnostics assess the mixing behavior within each MCMC chain. We here present the ESS of four variables, including the parsimony score summed across the sequence alignment, mean molecular evolutionary rate, joint posterior density, and tree topology. The topology ESS is computed following Fréchet correlation (3). Judging by the topology ESS, most datasets exhibit acceptable mixing (*i.e.*, ESS above 200) except for the human influenza virus (H1N1, H3N2, and B/Vic) and HIV datasets. The other ESS diagnostics, especially parsimony-score ESS, appear to be more conservative and indicate inadequate sampling. **(Middle)** PSRF of the above four variables (left axis) and ASDCF (average standard deviation of clade frequency; right axis) evaluate the convergence performance among MCMC replicate chains. Similar to the mixing behavior, most datasets appear to have converged on their tree estimates according to tree PSRF. Parsimony-score PSRF, however, appears to be more conservative, indicating lack of convergence for seven out of the 15 datasets, which corroborates more closely with the visual pattern shown in Fig. 3. The lack of convergence revealed by the parsimony-score PSRF (and to some extent by the posterior PSRF as well) is likely caused by a small fraction of clades (*i.e.*, local topology) that fail to converge between the chains, which is consistent with the pattern of ASDCF and the fraction of failed nodes shown in the bottom panel. **(Bottom)** For most datasets (except for the HIV dataset), only a small fraction of the clades (evaluated by clade occurrence and node age), sites, and interval-specific effective population size estimates failed to reach the acceptable threshold in MCMC diagnosis (*i.e.*, ESS below 200 or PSRF above 1.1 or SDCF above 0.1).

#### S1.3 Phylodynamic Posterior Tree Landscapes are Often Operationally Rugged

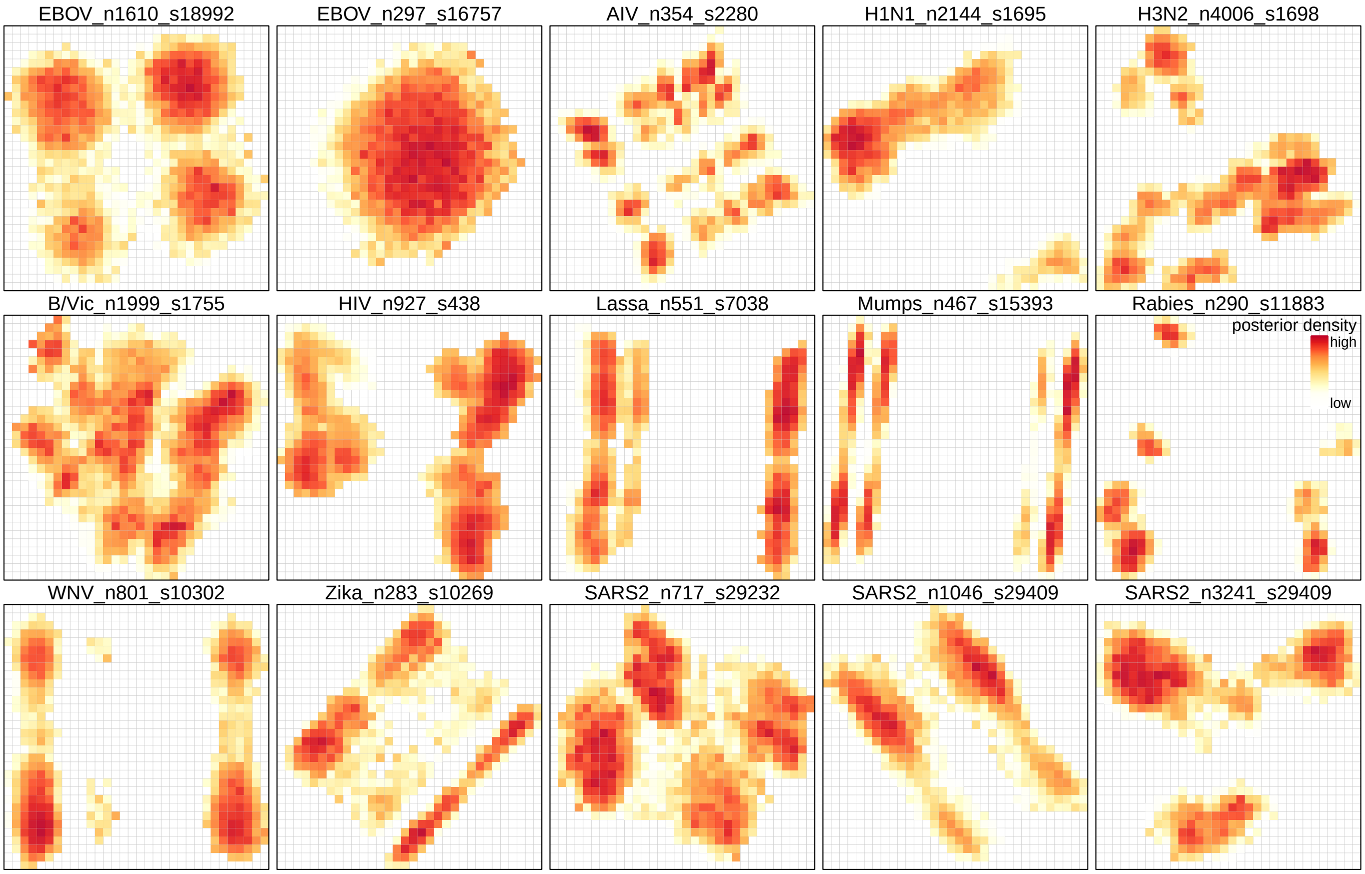


**Figure S6: Posterior density heatmaps across tree space reveal diffuse yet highly rugged phylodynamic tree landscapes.** Heatmap showing joint posterior density across the two-dimensional tree space—which effectively visualizes the posterior landscape of tree space—with warmer colors (red) indicating higher density. Each panel corresponds to a dataset. This figure is an alternative presentation of Fig. S7, highlighting the ruggedness of the posterior tree landscape and the qualitatively equivalent height of the local peaks. We discretize the two-dimensional MDS tree space (based on RF distances) into a $32 \times 32$ grid and then color each grid cell by the mean posterior density value of the tree samples in that cell. To reduce noise, cells with fewer than five samples are not colored; we compute the mean density value of each cell with up to 25 highest-density samples in that cell (approximately half of the average number of samples per cell).

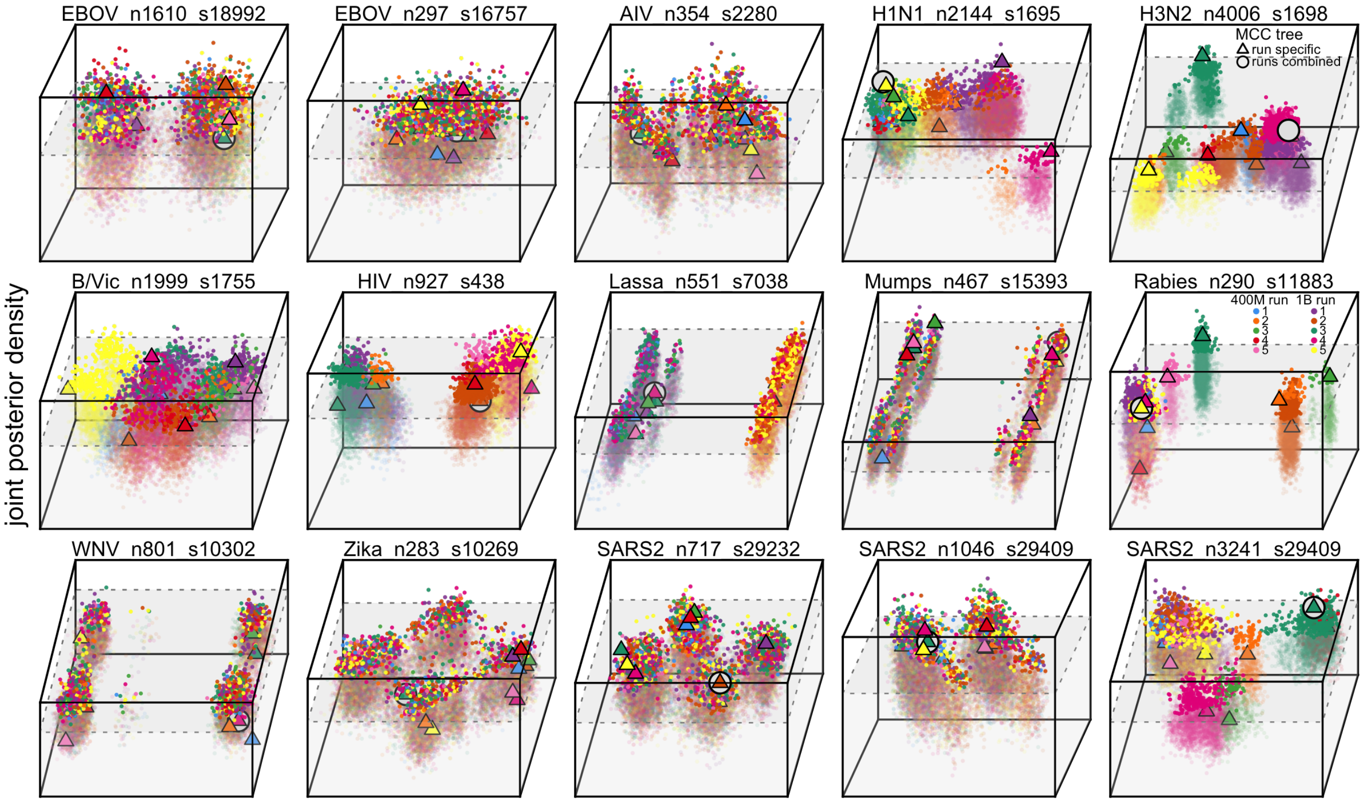


**Figure S7: Three-dimensional MDS plots reveal diffuse yet highly rugged phylodynamic tree landscapes.** Each panel visualizes the posterior landscape of tree space of a dataset. The *x*- and *y*- axes still correspond to the two MDS dimensions (generated based on the pairwise RF distances), respectively, while the *z* dimension is added to indicate the joint posterior density value ("posterior density") of each tree sample. That is, this figure would be identical to Fig. 3 when looking from a top-view angle. To highlight the peaks in the rugged posterior, we put a light-gray semi-transparent cube in this three-dimensional space to cover the part below the 10% highest posterior density region; therefore, the widely separated brighter dots higher up in the space indicate multiple peaks with virtually the same posterior density. A triangle represents the MCC tree summarized from each MCMC chain, while an open circle indicates the MCC tree summarized across all the replicates of that dataset.

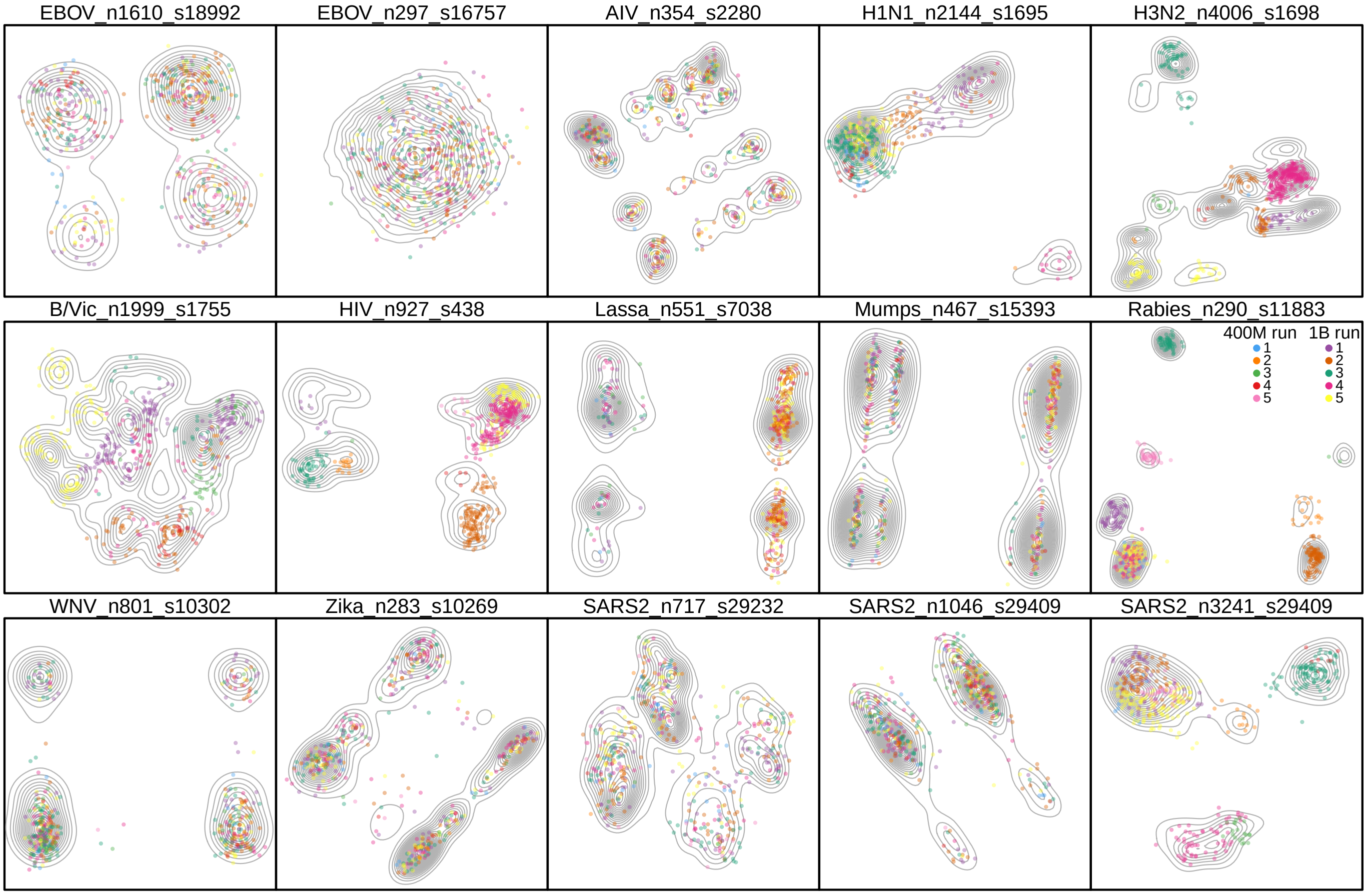


**Figure S8: Tree space MDS contour plots with highest-density tree samples indicate diffuse yet highly rugged phylodynamic tree landscapes.** Each panel shows the MDS plot visualizing the posterior tree space of a dataset. To highlight the peaks in the rugged posterior, we only keep the samples with the highest 1% posterior density in the MDS plot while adding the contours to indicate the distribution of all samples. Similar to Fig. 3, it reveals that, for most datasets, multiple peaks with similar height are present in the posterior tree space and individual chains frequently get stuck on one or a few of the isolated peaks.

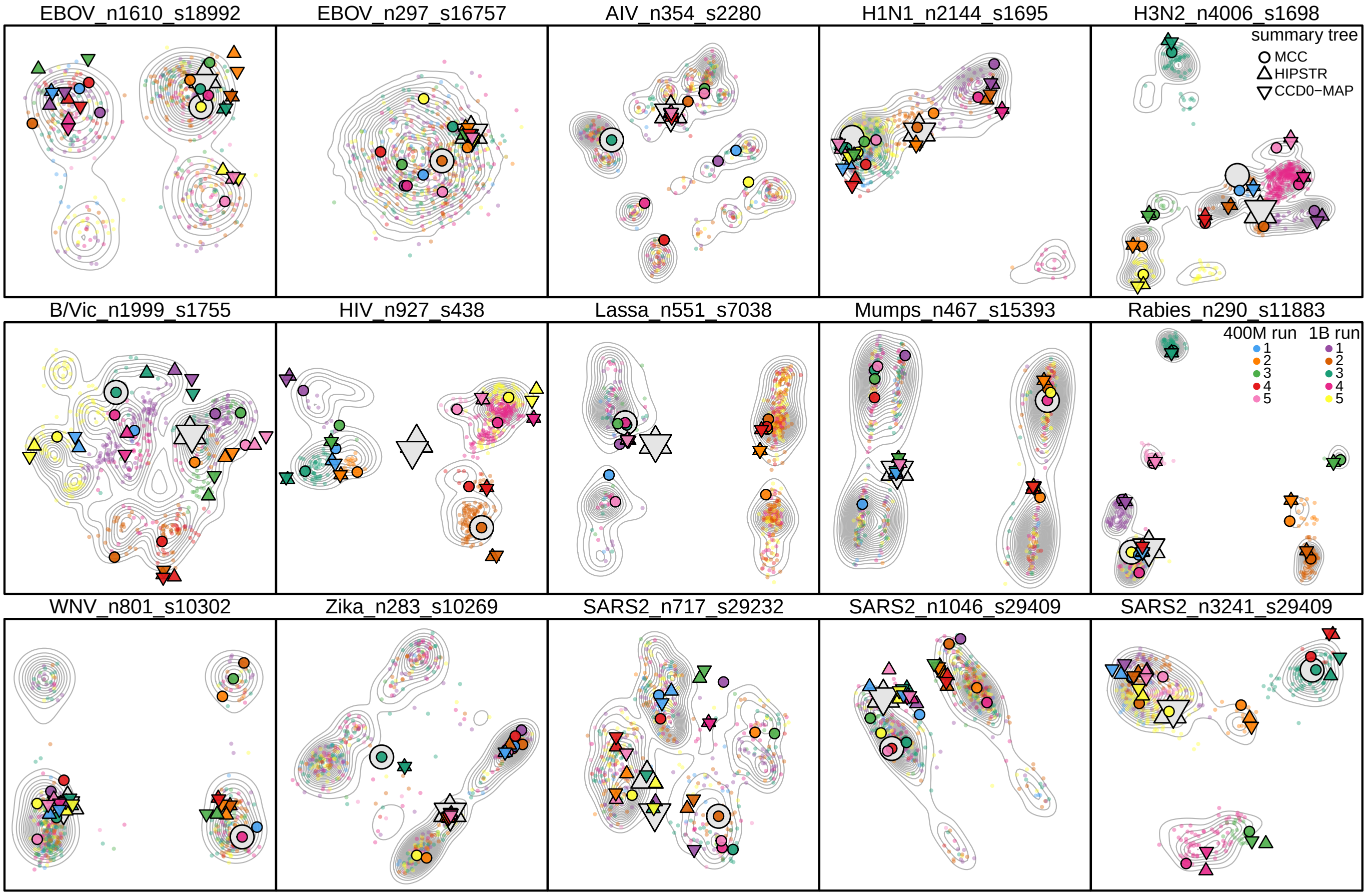


**Figure S9: Summary trees may not adequately represent multimodal phylodynamic tree space, while alternative summary methods can lead to distinct trees.** This figure extends Fig. S8 by adding summary trees. In addition to the commonly used MCC algorithm, we explored two alternative methods developed recently for summarizing posterior trees: HIPSTR (4) and CCD0-MAP (5). These two methods are more computationally and memory intensive than MCC, but have been reported to produce better summary trees. A fundamental difference distinguishing these two methods from MCC is that they can produce summary trees that were never sampled from the posterior. To generate HIPSTR and CCD0-MAP trees within reasonable time and memory constraints (a few days and several hundred gigabytes per dataset), we thinned the posterior to one tree per 100 thousand iterations when collecting clades and computing frequencies. For datasets where CCD0-MAP analyses would require substantially longer, we used its approximation algorithm (ACCD0-MAP). Note that the MCC trees may not be identical to those in Fig. 3, as the MCC trees here were also generated from thinned posterior samples to enable closer comparison among summary methods. A colored outlined symbol represents the summary tree of each chain, while an open symbol indicates the summary tree from all chains combined. Symbol shapes indicate summary methods (circle: MCC; up triangle: HIPSTR; down triangle: CCD0-MAP). In general, summary trees from individual chains often reside on different peaks, while the summary tree from all chains combined typically resides toward MDS plot corners. However, the combined HIPSTR and CCD0-MAP trees appear to be closer to the center than the corresponding MCC tree, especially when the posterior is multimodal. For some non-converging datasets with clearly isolated peaks (*e.g.*, HIV and Lassa), the combined HIPSTR and CCD0-MAP trees may lie between peaks, reflecting their ability to infer summary trees that were never sampled from the posterior. The HIPSTR and CCD0-MAP trees are usually very similar to each other. For datasets without convergence failure, chain-specific summary trees from HIPSTR and CCD0-MAP tend to be closer to their respective combined summary trees than those from MCC. This suggests that HIPSTR and CCD0-MAP may be less sensitive to sampling noise in individual chains and thus more robust to inadequate mixing.

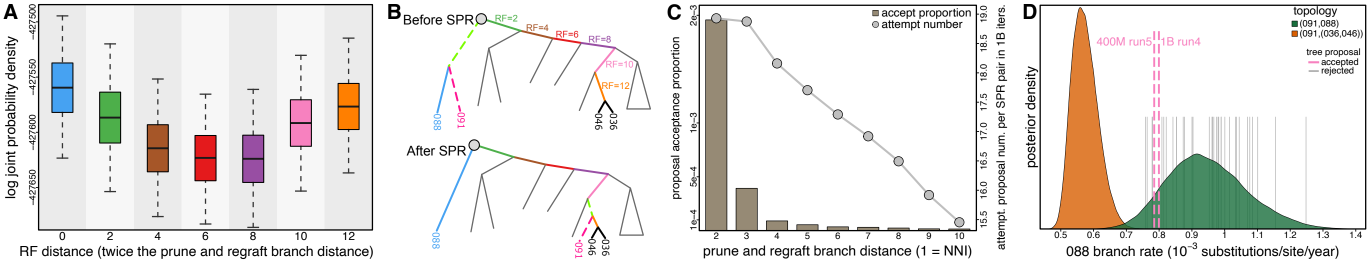


**Figure S10: Causes of the rugged tree landscape and tree sampling problems in the Lassa dataset with respect to alternative positions of the problematic sequence 091.** We additionally reveal the valley in the posterior tree landscape between the identified peaks associated with the two alternative positions of sequence 091 sampled across all the posterior trees of the Lassa dataset. The two topologies can reach each other by one SPR move crossing six nodes in the tree and have an RF distance of 12; the topologies where 091 is regrafted to the intermediate branches between those two alternative positions were never sampled. To demonstrate that those possible but unsampled topologies are associated with lower posterior probabilities, we performed additional phylodynamic analyses where the topology is fixed to be each of the two sampled topologies and the intermediate ones between them. **A)** Each boxplot corresponds to an alternative topology, colored according to the corresponding regraft branch shown in Panel **B**. The leftmost and rightmost boxplots correspond to the two sampled topologies, respectively. $Y$-axis shows the log joint probability density, so the topologies with higher boxplots at the two ends indicate the local peaks and the lower boxplots of intermediate topologies demonstrate the valley. **B)** The two alternative sampled subtree topologies highlighting the positions of focal tips and the intermediate branches in between the alternative positions of 091. **C)** Although the two local peaks can be commuted with one specific SPR move, it is proposed very infrequently and accepted extremely rarely. In general, due to the large number of tips and random selection of the prune and regraft branches, when performing Bayesian phylodynamic analysis of typical viral datasets, each specific SPR move is only proposed a handful to a few dozen times every one billion iterations and a very small fraction of these proposals will be accepted, resulting in widespread tree sampling problems when local peaks exist. Here, we visualize the mean number of proposal attempts for each SPR move in one billion iterations (dotted line, right $y$-axis) and the mean acceptance proportion of each proposal (bars, left $y$-axis) across various distances between the prune and regraft branches ($x$-axis) for the Lassa dataset summarized from the full MCMC history. In particular, when 091 and 088 were sisters, the SPR move required to reach the alternative topology was proposed 36 times in total across all the five billion post-burnin iterations, among which two were accepted, resulting in the the single valley crossing that occurred in two of the 10 MCMC replicate runs; in contrast, when 091 is sister to the cherry (036,046), the SPR move required to regraft 091 back onto the 088 branch is proposed 43 times in total with no acceptance. **D)** The distinct evolutionary rate distribution over the focal branches between the two alternative topologies (Fig. 4C) may contribute to the low acceptance rate of the SPR move required to commute between them. Specifically, as the rate over 088 branch tends to be much higher when 091 is sister to 088 than when they are not, the SPR moving 091 to its alternative position away from 088 may be rejected when the rate over 088 becomes too high for the more than doubled branch length after the move. Indeed, among the values of rate over 088 branch across all the 36 times that the SPR is proposed (thin gray bars), the values at the two rare accepted times (pink dashed lines) are close to the lowest, suggesting that the branch rates can be another contributing factor to the ruggedness of phylodynamic tree landscape.

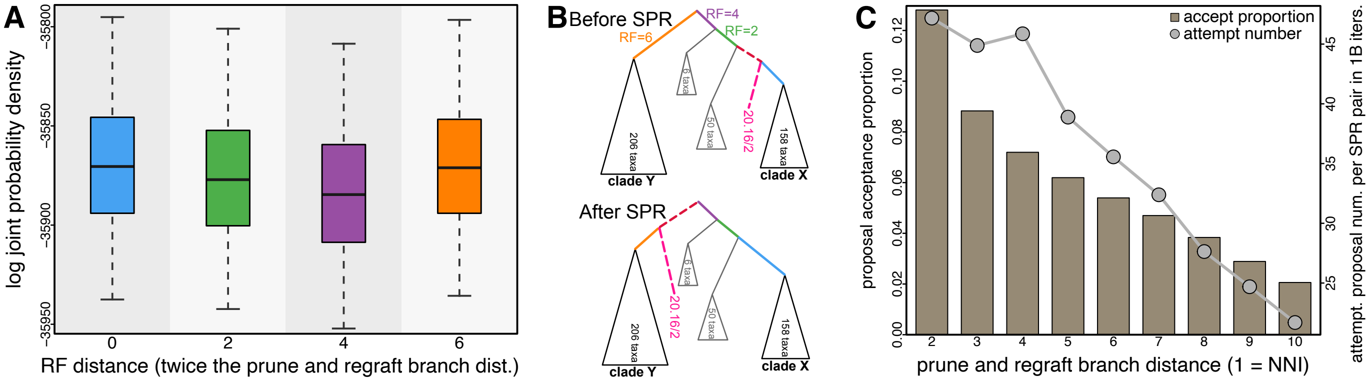


**Figure S11: Causes of the rugged tree landscape and tree sampling problems in the Mumps virus dataset with respect to alternative positions of the problematic sequence 20.16/2.** We additionally reveal the valley in the posterior tree landscape segregating the identified local peaks associated with the two alternative positions of sequence 20.16/2 sampled across all the posterior trees of the Mumps virus dataset. The two topologies can reach each other by one SPR move crossing three nodes in the tree and have an RF distance of six; the topologies where 20.16/2 is regrafted to the intermediate branches between those two alternative positions are almost never sampled. To demonstrate that those possible but rarely sampled topologies are associated with lower posterior probabilities, we set up additional phylodynamic analyses where the topology is fixed to be each of the two sampled topologies and the intermediate ones between them. **A)** Each boxplot corresponds to an alternative topology, colored according to the corresponding regraft branch shown in Panel **B**. The leftmost and rightmost boxplots correspond to the two frequently sampled topologies, respectively. $Y$-axis shows the log joint probability density, so the topologies with higher boxplots at the two ends indicate the local peaks and the lower boxplots of intermediate topologies demonstrate the valley. **B)** The two alternative sampled subtree topologies highlighting the positions of focal tips and the intermediate branches in between the alternative positions of 20.16/2. **C)** Although the two local peaks can be commuted with one SPR move, it is proposed infrequently. We visualize the mean number of proposal attempts for each SPR move in one billion iterations (dotted line, right $y$-axis) and the mean acceptance proportion of each proposal (bars, left $y$-axis) across various distances between the prune and regraft branches ($x$-axis) for the Mumps virus dataset summarized from the full record of the MCMC history. In particular, when 20.16/2 is at one of the two frequently sampled positions, the SPR move required to reach the alternative topology was proposed 354 times in total across all five billion post-burnin iterations, of which 79 were accepted. The occasional acceptance led to infrequent commutes between the local peaks and thus the slow mixing problem.

#### S1.4 Identifying Problematic Sequences and Assessing Their Impacts on Phylogenetic Inferences

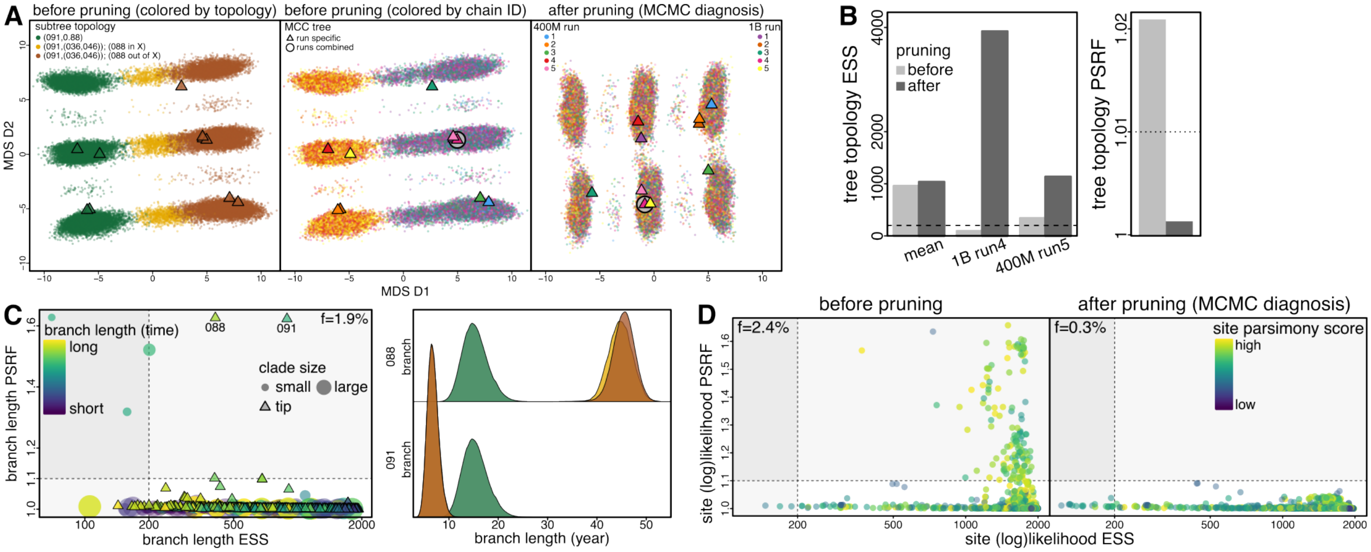


**Figure S12: Removing the two problematic sequences (088 and 091) effectively resolves tree sampling problems for the Lassa dataset. A)** MDS plot visualizing the posterior tree space (based on rSPR distances). Each dot represents a sampled tree; a triangle represents the MCC tree summarized from an MCMC chain, while an open circle indicates the MCC tree summarized across all chains. (Left and Middle) Posterior tree space before pruning, each dot is colored by the topology of its focal subtree (Left) or by the index of the MCMC chain the tree is sampled from (Middle). (Right) Tree space after the pruning where each dot is colored by its MCMC chain index (same as Middle). Panel **A** here is identical to Panels **B** and **E** of Fig. 4 combined, except that the MDS coordinates are computed with rSPR distances instead of RF distances. **B)** Tree topology diagnostics quantify MCMC performance before (light gray) and after (dark gray) the pruning, corroborating the visual pattern in **A**. (Left) The mean tree ESSs across all chains increase slightly after pruning; for the two chains where the sampler crossed the valley once before pruning, all the ESSs increase at least by an order of magnitude. (Right) Tree PSRF decreases dramatically after pruning the problematic sequences, becoming significantly smaller than the acceptable threshold. These tree topology diagnostics are generated similarly and qualitatively identical to the corresponding diagnostics presented in Panel **G** of Fig. 4, except that here the diagnostics are computed based on rSPR distances instead of RF distances. **C)** Length (in unit of time) of the branches subtending the two focal sequences (088 and 091) appears to be determined by local topology of the sampled tree. (Left) MCMC diagnostics computed for the length of each tree branch. PSRF ($y$-axis) quantifies convergence between the chains, and ESS ($x$-axis) quantifies mixing within each chain. Dashed lines indicate the thresholds typically used in phylogenetic studies; *i.e.*, PSRF above 1.1 (horizontal dashed line) indicates lack of convergence and ESS below 200 (vertical dashed line) indicates inadequate mixing. Circles and triangles represent internal and external branches, respectively, whose color indicates the inferred branch length value (yellow: high; purple: low) and size is proportional to the number of descendant tips. A small fraction ($f = 1.9\%$) of the dots reside outside the bottom right quadrant surrounded by the dashed lines, indicating that those branch lengths fail to be estimated reliably. Despite having seemingly sufficient samples indicated by their ESS values, length estimates of the two focal external branches (labeled in the plot) fail to converge. (Right) Posterior distribution of each branch length (top: 088; bottom: 091) conditioned on the sampled subtree topology (colored same as in **A**, Left). **D)** 2.4% of all sites in the sequence alignment, quantified by PSRF ($y$-axis) and ESS ($x$-axis) computed for the log-likelihood of each site, were associated with the convergence problem when the problematic sequences were present (Left). Each dot represents a site, whose color indicates the parsimony number of mutations at that site over the corresponding tree. The fraction of failed sites decreases to 0.3% after pruning, with all the PSRF values that were above the threshold dropping to below it.

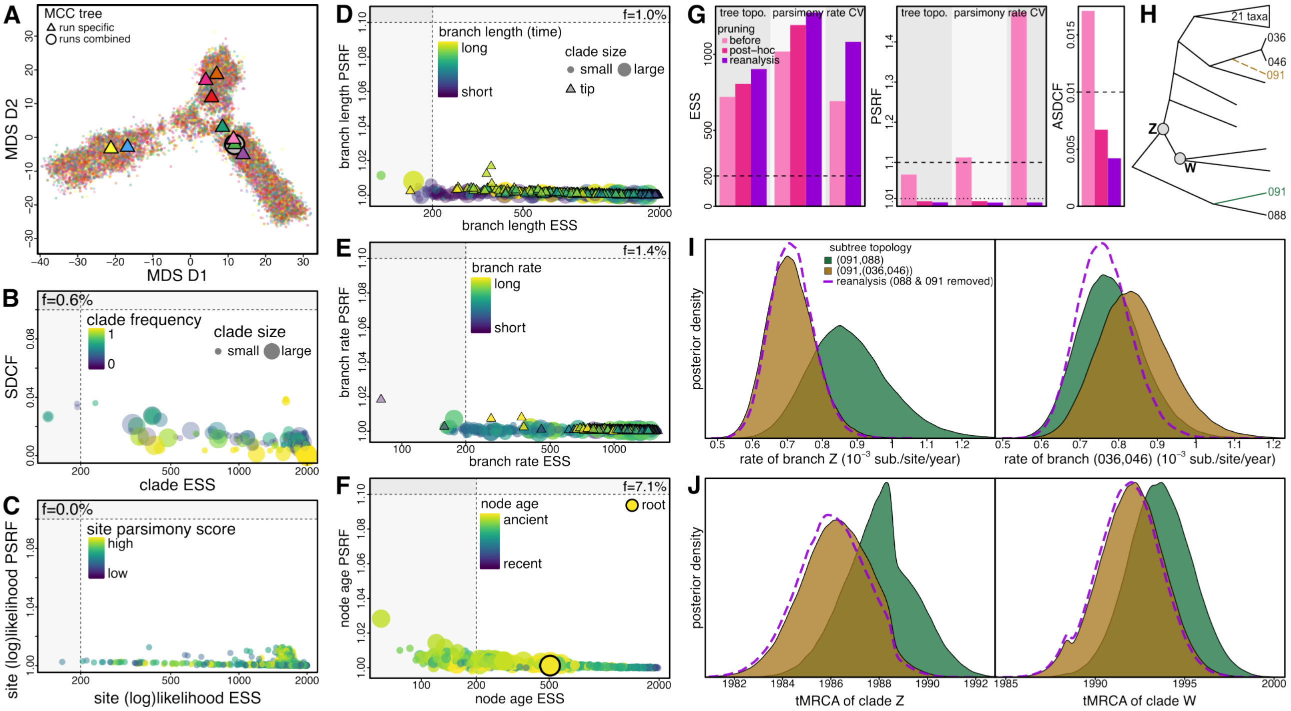

**Figure S13: Reanalyses of the Lassa dataset without the problematic sequences (088 and 091) yield a unimodal tree space with satisfactory sampling performance.** In addition to demonstrating the impact of the identified problematic sequences on tree space and tree sampling problems by pruning them from each sampled tree and then re-assessing MCMC performance with the pruned trees as a post-hoc treatment, we also reran the phylodynamic analyses using a sequence alignment with those sequences removed as a prospective assessment of their impact. The results presented here corroborate Fig. 4, revealing that the post-hoc pruning and prospective removal lead to similar dramatic changes in tree space structures and attenuation of tree sampling problems. Moreover, the reanalyses after removing the problematic sequences additionally demonstrate the significant attenuation of the sampling problems with other continuous parameters (*e.g.*, node ages) for which the post-hoc pruning is inapplicable to assess. **A)** MDS plot visualizing the posterior tree space (based on RF distances) of the problematic-sequence removed reanalyses. Each dot represents a sampled tree; a triangle represents the MCC tree summarized from an MCMC chain, while an open circle indicates the MCC tree summarized across all chains. **B)** Clade occurrence diagnostics of the reanalyses. SDCF ($y$-axis) and clade ESS ($x$-axis) quantify between-chain convergence and within-chain mixing, respectively. Each dot represents a clade, whose color indicates the clade frequency averaged over all chains. The non-converging clades (SDCF above 0.1) no longer exist under the prospective removal reanalysis, and the fraction of clades with poor MCMC performance (0.6%) is similar to that under the post-hoc pruning (0.4%), while both are much smaller than the original analysis fraction (2.9%; Fig. 4F). **C)** The fraction of failed sites in the sequence alignment under the prospective removal reanalysis is 0.0%, which is similar to that under the post-hoc pruning (0.3%), both much smaller than the original analysis fraction (2.4%; Fig. S12D). **D)** MCMC diagnostics computed for the length (in unit of time) of each tree branch in the reanalyses. Circles and triangles represent internal and external branches, respectively, whose color indicates the inferred branch length value (yellow: high; purple: low) and size is proportional to the number of descendant tips. The fraction of branch lengths fail to be estimated reliably decreases to 1.0% under the prospective removal reanalysis from 1.9% (original analysis fraction; Fig. S12C). **E)** The fraction of branch rates fail to be estimated reliably decreases to 1.4% under the prospective removal reanalysis from 1.9% (original analysis fraction; Fig. 4C). **F)** The fraction of node ages fail to be estimated reliably decreases to 7.1% under the prospective removal reanalysis from 13.4% (original analysis fraction; Fig. 4D). Across all the parameter diagnostics, none of them have a PSRF value above or even close to the threshold (1.1), in sharp contrast with those in the original analyses with the problematic sequences. **G)** Tree-based diagnostics quantify MCMC performance for original analysis (light pink), after post-hoc pruning (darker pink), and for prospective reanalysis (purple). Under both pruning treatments, the mean tree and parsimony-score ESSs increase noticeably (Left), while the tree and parsimony-score PSRFs (Middle) as well as the ASDCF (Right) decrease significantly. We additionally compare the ESS and PSRF values of the coefficient of variation of the evolutionary rates among branches (“rate CV”, a continuous parameter of the relaxed-clock model) between original analyses and prospective reanalyses, highlighting the dramatic decrease in its PSRF values after removing the problematic sequences, which corroborates the topology dependent branch rates associated with the problematic sequences as revealed by Fig. 4C. **H)** Subtree focusing on the problematic sequences (088 and 091). Dashed branches indicate the alternative position of 091 when it is not sister to 088. The two focal clades whose associated estimates are visualized in Panels **I** and **J** are marked on the tree. **I)** Evolutionary rate estimates over the branches subtending nodes close to the problematic sequences are strongly affected by the presence or absence of the problematic sequences, showing the inferred posterior distributions of the rate over the subtending branch of clade Z (Right) and of clade (036,046) (Left) under alternative topologies concerning the position of 091 (filled curves) or when 088 and 091 are removed from the alignment (dashed curve). **J)** Age estimates of the nodes close to the problematic sequences are also strongly affected by the problematic sequences, showing the node age distributions of Z (Left) and W (right). For all these clade-specific parameters, we demonstrate the impact of the problematic sequences by revealing that the estimates conditioning on 091 being topologically distant are qualitatively similar to those when 091 is excluded while distinct from the same estimates when 091 is topologically close to that clade.

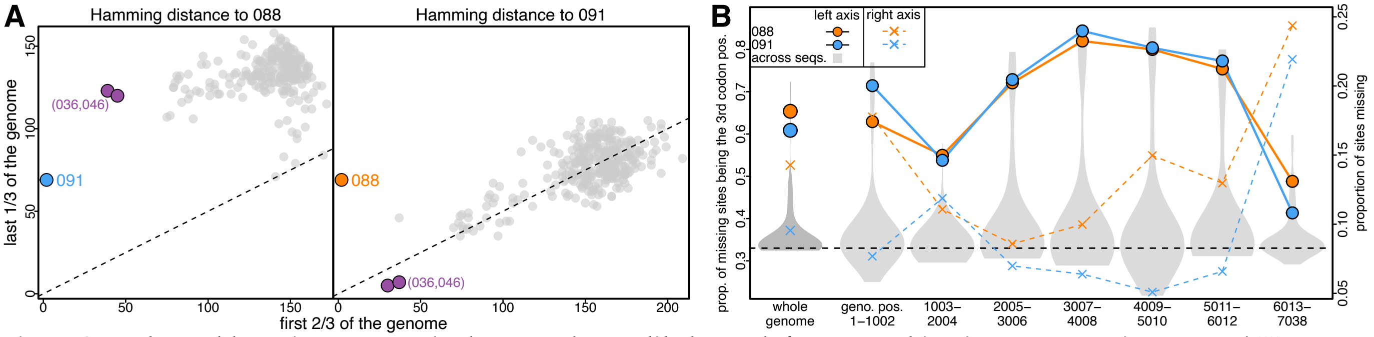


**Figure S14: The problematic sequences in the Lassa dataset likely result from recombination or sequencing error. A)** Hamming distances (*i.e.*, number of incompatible sites) between 088 and its closely related sequences (Left) and between 091 and its closely related sequences (Right). The distances from 088 to the other sequences in the last $\frac{1}{3}$ of the genome is significantly larger than those in the first $\frac{2}{3}$ of the genome (*i.e.*, the dots are mostly far above the dashed line, which corresponds to the case where it is equally likely for a site to be incompatible whether it is in the first $\frac{2}{3}$ or the last $\frac{1}{3}$ of the genome). Specifically, the distance between 088 and 091 is 69 in the last $\frac{1}{3}$ of the genome, but merely 2 in the first $\frac{2}{3}$ of the genome. By contrast, the distances from 091 to the other sequences, except for that to 088, are virtually evenly distributed along the genome. **B)** Noticeable proportions of sites in both 088's and 091's genomes are missing (crosses, colored dashed lines, right axis), especially at the two ends of the genomes. Surprisingly, most of the missing sites are on the third codon position, which is much higher than expected under the null expectation ($\frac{1}{3}$, horizontal dashed line). Together, these lines of evidence suggest that these two problematic sequences might each have been produced by taking a consensus sequence from mixed viral samples due to superinfection or contamination.

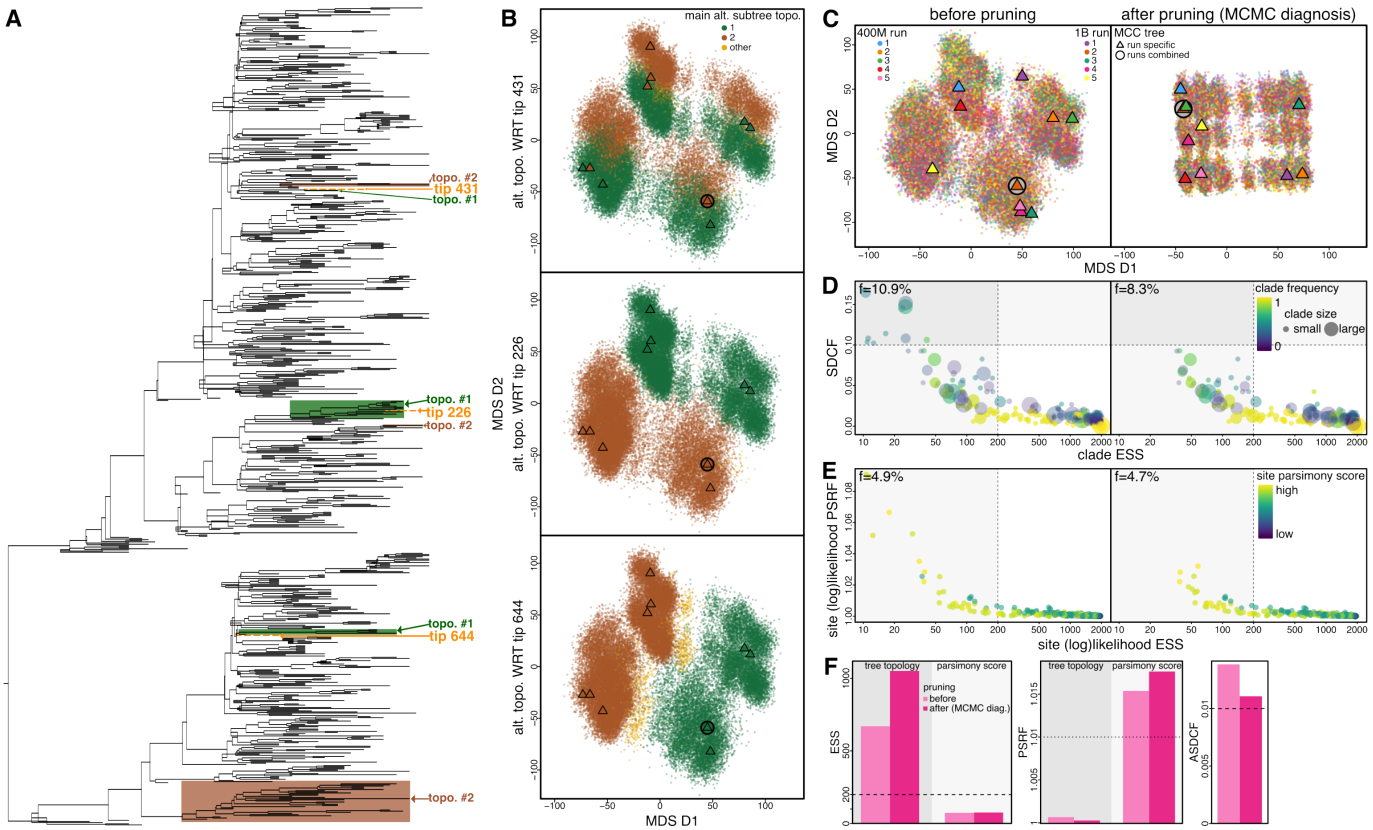


**Figure S15: Tree sampling problems with the SARS-CoV-2 Origin dataset appear to be mainly caused by a few sequences.** **A)** MCC tree of the SARS-CoV-2 Origin dataset (6) focusing on the problematic sequences [tip number (GISAID ID): tip 226 (EPI_ISL_414569), tip 431 (EPI_ISL_962567), tip 644 (EPI_ISL_406223)]. Dashed branches indicate the three problematic tips identified by MCMC diagnosis, with the two major alternative topological positions of each identified tip marked by color filled rectangles (one green and one brown for each tip). **B)** MDS plot visualizing the posterior tree space (based on RF distance) with each sampled tree colored by the topology of its focal subtree. Each subpanel corresponds to a problematic tip shown in **A** and the colors follow those in **A**; *i.e.*, these subpanels are identical other than the dot color. Each dot represents a sampled tree; a triangle represents the MCC tree summarized from an MCMC chain, while an open circle indicates the MCC tree summarized across all chains. **C)** Posterior tree space before (Left) and after (Right) pruning the problematic sequences from each sampled tree, with the color of each dot showing which MCMC chain the sampled tree is from (*i.e.*, Left of **C** is identical to the subpanels in **B** other than the dot color). (Left) Before pruning, at least eight peaks (potentially caused by the three problematic sequences) appear in the posterior tree space and the sampler commuted between them somewhat infrequently; (Right) after the pruning, mixing is significantly improved as the peaks vanish and sampler moves around the space much more efficiently. **D)** Clade occurrence MCMC diagnostics before (Left) and after (Right) the pruning. SDCF ($y$-axis) quantifies the between-chain convergence and clade ESS ($x$-axis) quantifies the within-chain mixing performance, respectively. Each dot represents a clade, whose color indicates the clade frequency averaged across all replicates. All the clades whose SDCF were above 0.1 "disappear" after pruning and the fraction of clades with unsatisfactory MCMC performance (*i.e.*, SDCF greater than 0.1 or ESS smaller than 200) decreases from 10.9% to 8.3%, indicating the attenuation of the convergence issue. **E)** 4.9% of all sites in the sequence alignment, quantified by PSRF ($y$-axis) and ESS ($x$-axis) computed for the log-likelihood of each site, failed to converge with the presence of the problematic sequences (Left). Each dot represents a site, whose color indicates the parsimony number of mutations at that site over the corresponding tree. The fraction of failed sites decreases only slightly to 4.7% after the pruning, but the five sites with the largest PSRFs and smallest ESSs "disappear". **F)** Tree-based diagnostics quantify MCMC performance before (light pink) and after (pink) the pruning, corroborating the visual pattern in **C**. The mean tree and parsimony-score ESSs increase, while the tree and parsimony-score PSRFs as well as the ASDCF decrease, after pruning off the problematic sequences identified by MCMC diagnosis.

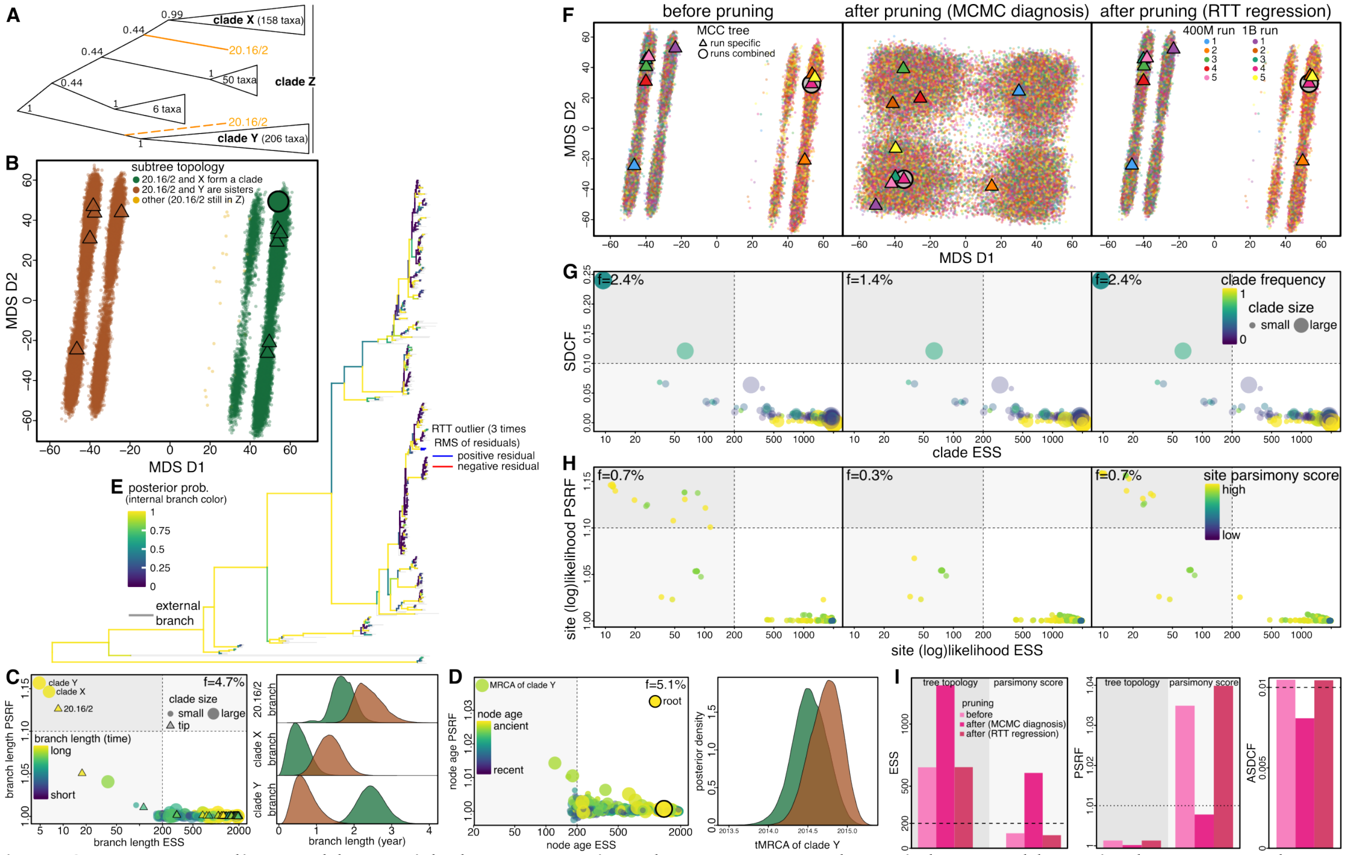

**Figure S16: Tree sampling problems with the Mumps virus dataset appear to be mainly caused by a single sequence that can be identified by MCMC diagnosis but not by RTT regression test.** **A)** Subtree of the Mumps virus dataset (7) focusing on the problematic sequence 20.16/2 (full name: Massachusetts.USA/20.16/2/G|2016-05-20|north_america|usa|massachusetts). Dashed branches indicate the alternative topological position of sequence 20.16/2. **B)** MDS plot visualizing the posterior tree space (based on RF distances) with each sampled tree colored by the topology of its focal subtree. Each dot represents a sampled tree; a triangle represents the MCC tree summarized from an MCMC chain, while an open circle indicates the MCC tree summarized across all chains. **C)** Length (in unit of time) of the branches subtending the focal clades appears to be determined by local topology of the sampled tree, to the extent that a certain branch varies by several times given one topology versus the other. (Left) MCMC diagnostics computed for the length of each branch. Dashed lines indicate the thresholds typically used in phylogenetic studies; *i.e.*, PSRF above 1.1 (horizontal dashed line) indicates lack of convergence and ESS below 200 (vertical dashed line) indicates inadequate mixing. Circles and triangles represent internal and external branches, respectively, whose color indicates the inferred branch length value (yellow: high; purple: low) and size is proportional to the number of descendant tips. A small fraction ($f = 4.7\%$) of the dots reside outside the bottom right quadrant surrounded by the dashed lines, indicating that those branch lengths fail to be estimated reliably. Length estimates of the focal branches (labeled in the plot) fail to converge. (Right) Posterior distribution of each branch length conditioned on the sampled subtree topology (colored same as in **B**). **D)** Age estimates of the nodes (*i.e.*, tMRCA) that are close to 20.16/2 can also be strongly affected by tree sampling problems. (Left) Node-age PSRF of clade Y reveals the corresponding estimates failed to converge among replicates. (Right) tMRCA of clade Y is inferred to be significantly younger when it is sister to 20.16/2. **E)** MCC tree with the two tips identified as outliers in the root-to-tip (RTT) regression test (marked in the tree). **F)** Posterior tree space before and after pruning problematic sequences from each sampled tree, with the color of each dot showing which MCMC chain the sampled tree is from (*i.e.*, the dots in Left of **F** are identical to those in **B** other than their colors). (Left) Before pruning, the sampler crossed the valley infrequently, with the MCC trees summarized from different chains separated by the valley. (Middle) The valley largely vanishes after pruning off sequence 20.16/2 (identified by MCMC diagnosis), indicating that mixing is greatly improved. (Right) The tree space is effectively unchanged after pruning off the RTT test outliers. **G)** Clade occurrence MCMC diagnostics computed with the original posterior trees (Left), after pruning the tips identified by MCMC diagnosis (Middle) or by the RTT regression test (Right). SDCF ($y$-axis) quantifies the between-chain convergence and clade ESS ($x$-axis) quantifies the within-chain mixing performance, respectively. Each dot represents a clade, whose color indicates the clade frequency averaged across all replicates. The clades with the highest SDCFs and lowest ESSs disappear after pruning off 20.16/2 and the fraction of clades with unsatisfactory MCMC performance decreases from 2.3% to 1.3%, indicating the attenuation of the convergence issue. By contrast, pruning off the RTT outliers does not alleviate the sampling problems. **H)** 0.7% of all sites in the sequence alignment, quantified by PSRF ($y$-axis) and ESS ($x$-axis) computed for the log-likelihood of each site, failed to converge in the original posterior (Left). Each dot represents a site, whose color indicates the parsimony number of mutations at that site over the corresponding tree. The fraction of failed sites decreases to 0.3% after pruning off 20.16/2, with all the PSRF values that were above the threshold (1.1) dropping to below it. Conversely, the site-specific MCMC diagnostics are effectively unchanged after pruning off the RTT outliers. **I)** Tree-based diagnostics quantify MCMC performance before (light pink) and after (darker pinks) the pruning, corroborating the visual pattern in **F**. The mean tree and parsimony-score ESSs increase dramatically, while the tree and parsimony-score PSRFs as well as the ASDCF decrease noticeably, after pruning off 20.16/2 but not after pruning off the RTT outliers.

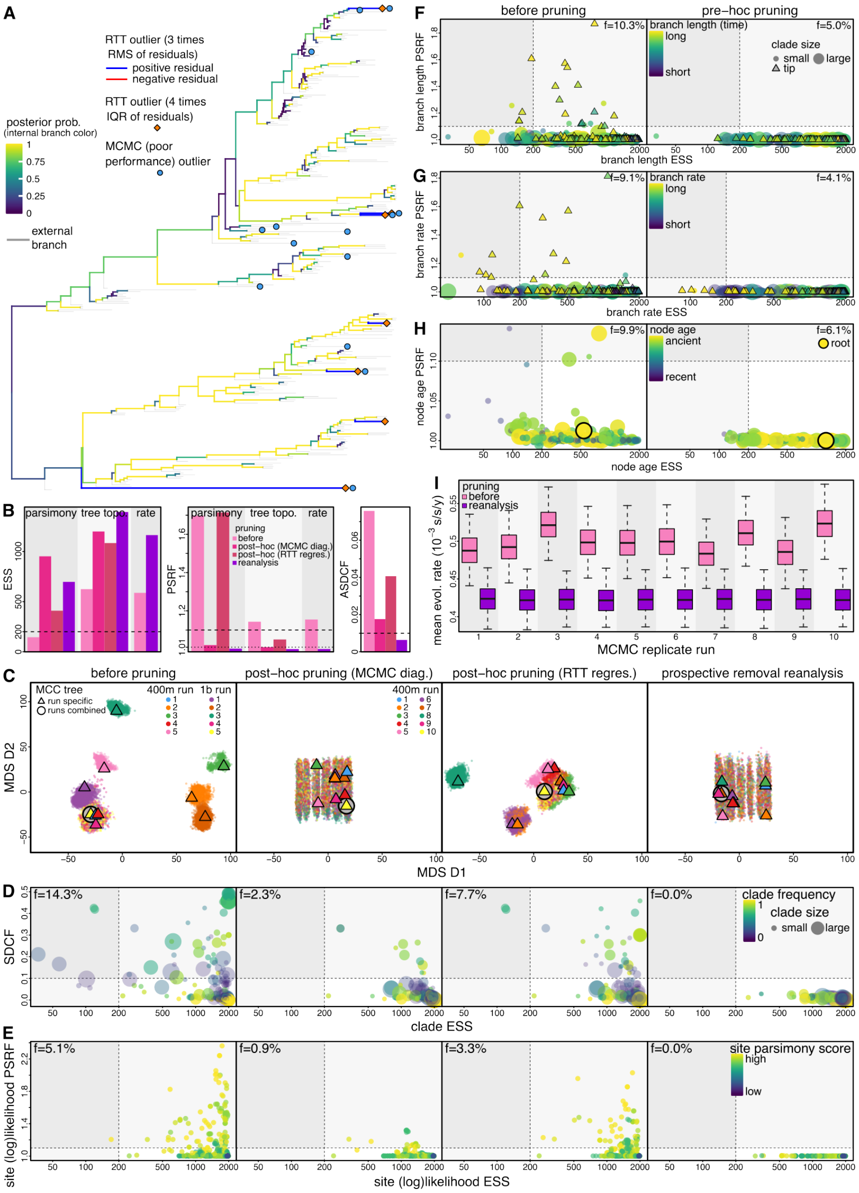


**Figure S17: Tree sampling problems with the rabies virus dataset appears to be mainly caused by 14 sequences identified by MCMC diagnosis.** **A)** MCC tree of the rabies virus dataset (8) focusing on the problematic sequences. The problematic sequences are marked to indicate the ones identified by MCMC diagnosis (blue circles) or RTT regression test (orange diamond and blue subtending branches). Five of those sequences were identified by both approaches. **B)** Tree-based diagnostics quantify MCMC performance for original analyses (light pink), after post-hoc pruning treatments (darker pinks), and for prospective removal reanalyses (purple), corroborating the visual pattern in Panel **C**. The mean tree and parsimony-score ESSs increase, while the tree and parsimony-score PSRFs as well as the ASDCF decrease, after pruning off the problematic sequences identified by MCMC diagnosis. Pruning off the RTT outliers induces a similar pattern but to a much lesser extent. In addition to demonstrating the impact of the identified problematic sequences by pruning them from each sampled tree as a post-hoc treatment, we also reran the phylodynamic analyses using a sequence alignment with the sequences identified by MCMC diagnosis removed as a prospective assessment of their impact. Similar to the corresponding post-hoc pruning treatment, this prospective removal reanalysis led to a dramatic improvement on tree sampling performance. Moreover, we compare the ESS and PSRF values of the mean evolutionary rate among branches ("rate") between the original analyses and prospective removal reanalyses, highlighting the significant improvement on sampling this biologically important parameter after removing the problematic sequences; the inferred posterior distributions of the mean rate across replicates under these two settings are shown in Panel **I**. **C)** Posterior tree space before and after pruning the problematic sequences. (Leftmost) Before pruning, conspicuous valleys segregating the peaks and the sampler was unable to cross the valleys. (Left-center) The valley largely vanished after pruning off the tips identified by MCMC diagnosis, indicating that tree sampling is improved. (Right-center) Some but not all of the valleys vanished after pruning off the RTT test outliers. (Rightmost) Similar to the corresponding post-hoc pruning, the prospective removal led to significant improvement in tree sampling, consistent with the tree-based diagnostics shown in Panel **B**. **D)** Clade occurrence MCMC diagnostics computed with the original posterior trees (Leftmost), after pruning the tips identified by MCMC diagnosis (Left-center) or by RTT regression test (Right-center), and with the posterior trees from the prospective removal reanalyses (Rightmost). The clades with the highest SDCF and lowest ESS (*i.e.*, dots in the top left quadrant) disappear after pruning off the tips identified by MCMC diagnosis and the fraction of clades with unsatisfactory MCMC performance decreases from 14.3% to 2.3%, indicating the attenuation of the sampling problems. Pruning off the RTT outliers also alleviates the sampling problems but to a lesser degree (7.7%). Prospective removal further reduces the fraction of failed clades to 0.0%. **E)** 5.1% of all sites in the sequence alignment failed to converge in the original posterior (Leftmost). The fraction of failed sites decreases dramatically to 0.9% after pruning off the tips identified by MCMC diagnosis, with most of the PSRF values that were above the threshold (1.1) dropping to below it (Left-center). The fraction of failed sites decreases to a lesser extent (3.3%) after pruning off the RTT outliers (Right-center). No site failed under the prospective removal reanalyses (Rightmost). **F)** MCMC diagnostics computed for the duration of each tree branch in the original and pruned reanalyses. The fraction of branch lengths fail to be estimated reliably decreases from 10.3% to 5.0% after the pruning. **G)** The fraction of branch rates fail to be estimated reliably decreases from 9.1% to 4.1% after the pruning. **H)** The fraction of node ages fail to be estimated reliably decreases from 9.9% to 6.1% after the pruning. Across all the parameter diagnostics, none of them have a PSRF value above or even close to the threshold (1.1), in sharp contrast with those in the original analyses. **I)** Inferred posterior distributions of the mean evolutionary rate across replicates in the original (pink) and pruned reanalyses (purple). The estimates from the original analyses exhibit apparent lack of convergence among replicate runs, consistent with the corresponding PSRF presented in Panel **B**. By contrast, the estimates from the pruned reanalyses converge perfectly among replicates. We note that the data are different between these two sets of analyses, resulting in distinct parameter estimates—where the quantitative differences can be biologically significant—in addition to MCMC performance improvement.

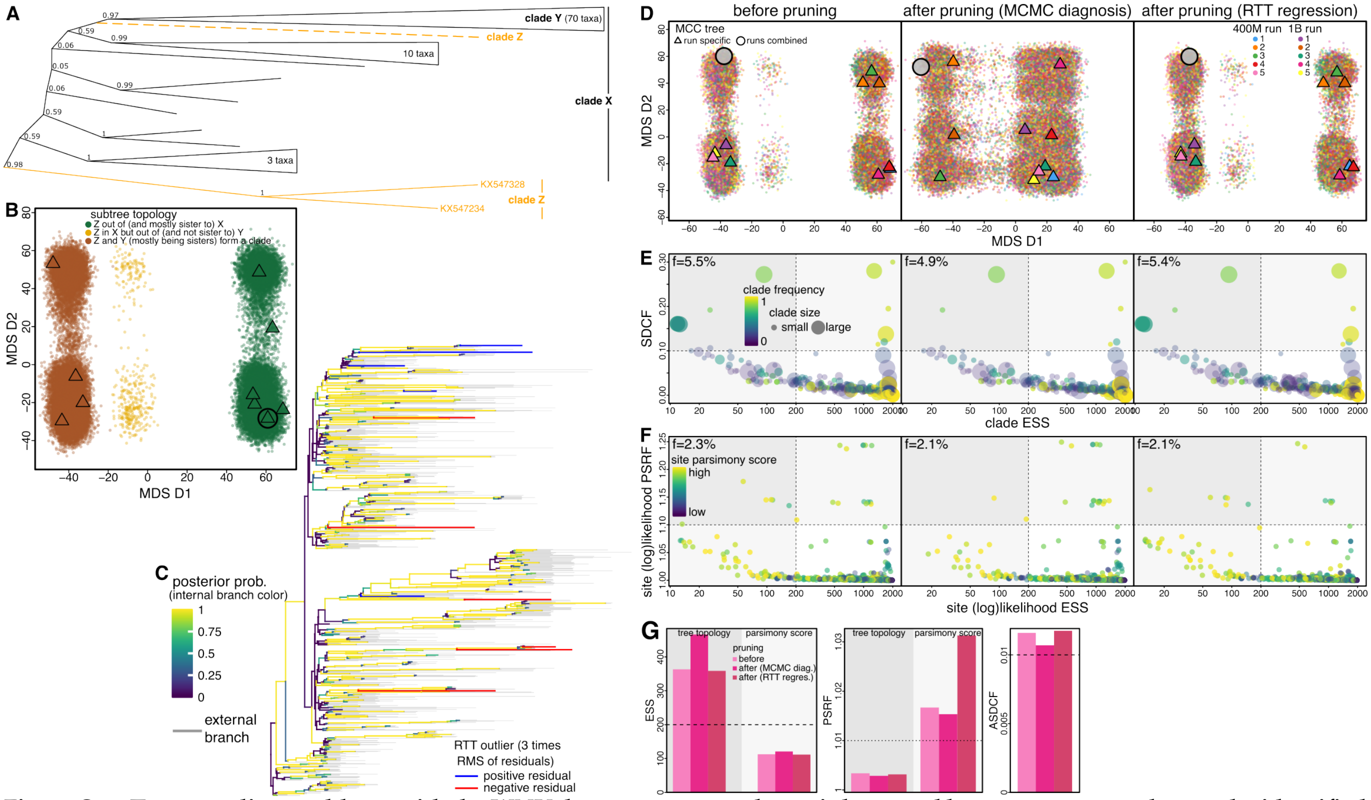


**Figure S18: Tree sampling problems with the WNV dataset appears to be mainly caused by two sequences that can be identified by MCMC diagnosis but not by RTT regression test.** **A)** Subtree of the West Nile virus (WNV) dataset (9) focusing on the problematic sequences. The dashed branch indicates the alternative topological position of clade Z that comprises those two problematic sequences. **B)** MDS plot visualizing the posterior tree space (based on RF distances) with each sampled tree colored by the topology of its focal subtree. Each dot represents a sampled tree; a triangle represents the MCC tree summarized from an MCMC chain, while an open circle indicates the MCC tree summarized across all chains. **C)** MCC tree with the tips identified as outliers in the root-to-tip (RTT) regression test (marked in the tree). **D)** Posterior tree space before and after pruning problematic sequences from each sampled tree, with the color of each dot showing which MCMC chain the sampled tree is from (*i.e.*, the dots in the left panel here are identical to those in **B** other than their colors). (Left) Before pruning, a conspicuous valley separated the peaks and the tree sampler crossed the valley infrequently. (Middle) The valley largely vanishes after pruning off clade Z (identified by MCMC diagnosis), indicating that mixing is improved. (Right) The tree space is effectively unaltered after pruning off the RTT outliers. **E)** Clade occurrence MCMC diagnostics computed with the original posterior trees (Left), after pruning the tips identified by MCMC diagnosis (Middle) or by the RTT regression test (Right). SDCF ($y$-axis) quantifies the between-chain convergence and clade ESS ($x$-axis) quantifies the within-chain mixing performance, respectively. Each dot represents a clade, whose color indicates the clade frequency averaged across all replicates. The clades with the lowest ESSs disappear after pruning off clade Z and the fraction of clades with unsatisfactory MCMC performance decreases from 7.1% to 6.4%, indicating the attenuation of the sampling problems. By contrast, pruning off the RTT outliers does not alleviate the sampling problems. **F)** 2.3% of all sites in the sequence alignment, quantified by PSRF ($y$-axis) and ESS ($x$-axis) computed for the log-likelihood of each site, failed to converge in the original posterior (Left). Each dot represents a site, whose color indicates the parsimony number of mutations at that site over the corresponding tree. The fraction of failed sites decreases only slightly after the pruning. **G)** Tree-based diagnostics quantify MCMC performance before (light pink) and after (darker pinks) the pruning, corroborating the visual pattern in **D**. The mean tree and parsimony-score ESSs increase, while the tree and parsimony-score PSRFs as well as the ASDCF decrease, after pruning off clade Z but not after pruning off the RTT outliers.

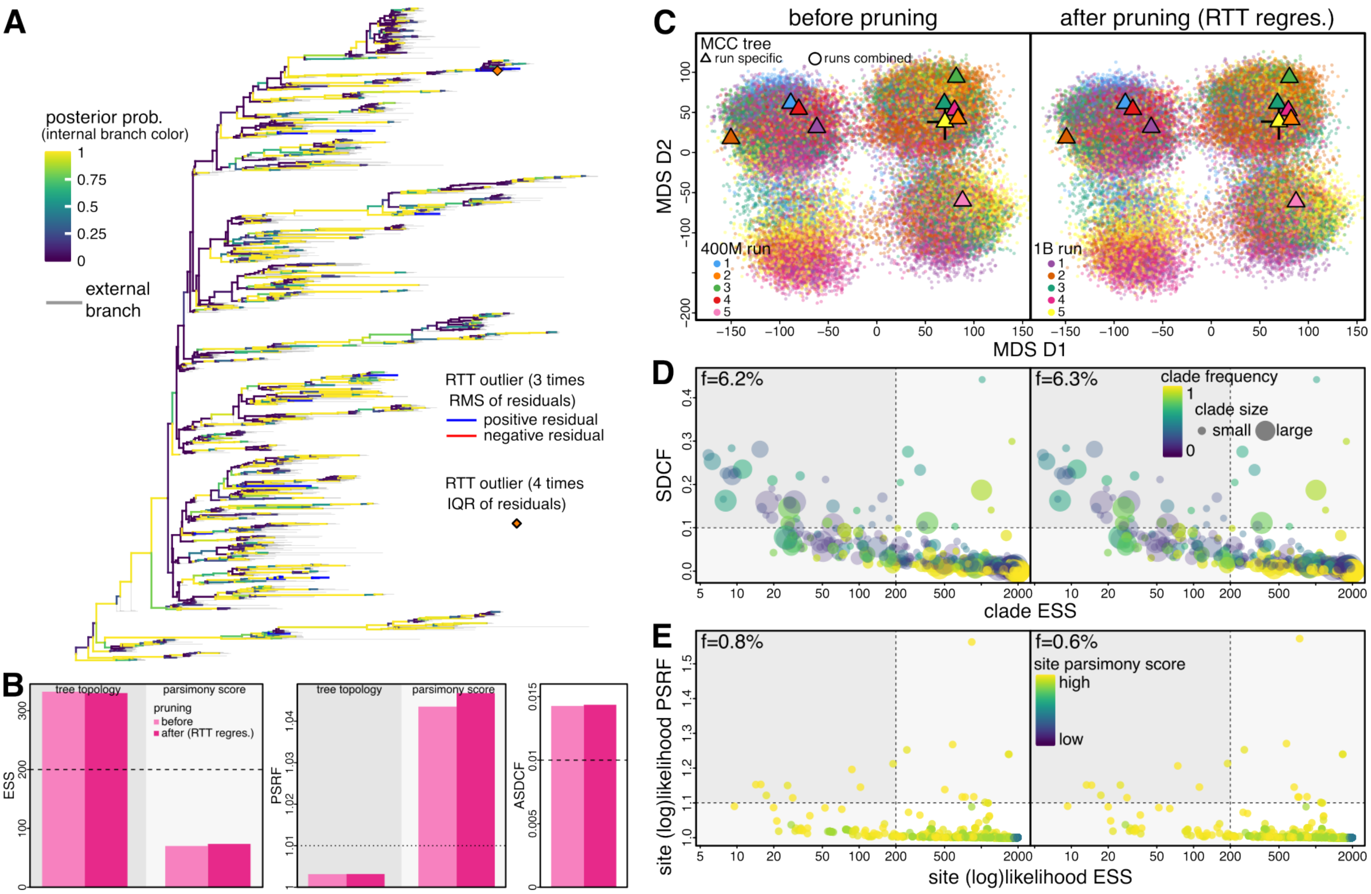


**Figure S19: Pruning off the RTT outlier sequences has little impact on MCMC performance for the Ebola virus dataset.** Note that for this dataset the number of sequences causing the poorly sampled clades revealed by the MCMC diagnosis are too many ($\gg$10) to examine closely and a crude removal of all such sequences might result in a significantly reduced tree that loses part of its topology backbone, so we did not assess the impact of pruning off sequences based on the MCMC diagnostics here. **A)** MCC tree of the Ebola virus West African Outbreak dataset (10) focusing on the problematic sequences identified as outliers in the RTT regression test (marked as orange diamond and blue subtending branches). **B)** Tree-based diagnostics that quantify MCMC performance before (light pink) and after (pink) the pruning corroborate the visual pattern shown in **C**. **C)** Posterior tree space before (Left) and after (Right) pruning the RTT outlier sequences from each sampled tree, with the color of each dot showing which MCMC chain the sampled tree is from. Each dot represents a sampled tree; a triangle represents the MCC tree summarized from an MCMC chain, while an open circle indicates the MCC tree summarized across all chains. **D)** Clade occurrence MCMC diagnostics before (Left) and after (Right) the pruning. SDCF ($y$-axis) quantifies the between-chain convergence and clade ESS ($x$-axis) quantifies the within-chain mixing performance, respectively. Each dot represents a clade, whose color indicates the clade frequency averaged across all replicates. **E)** Site-specific MCMC diagnostics before (Left) and after (Right) the pruning. The site-specific PSRF ($y$-axis) and ESS ($x$-axis) are computed using the log-likelihood of each site. Each dot represents a site, whose color indicates the parsimony number of mutations at that site over the corresponding tree.

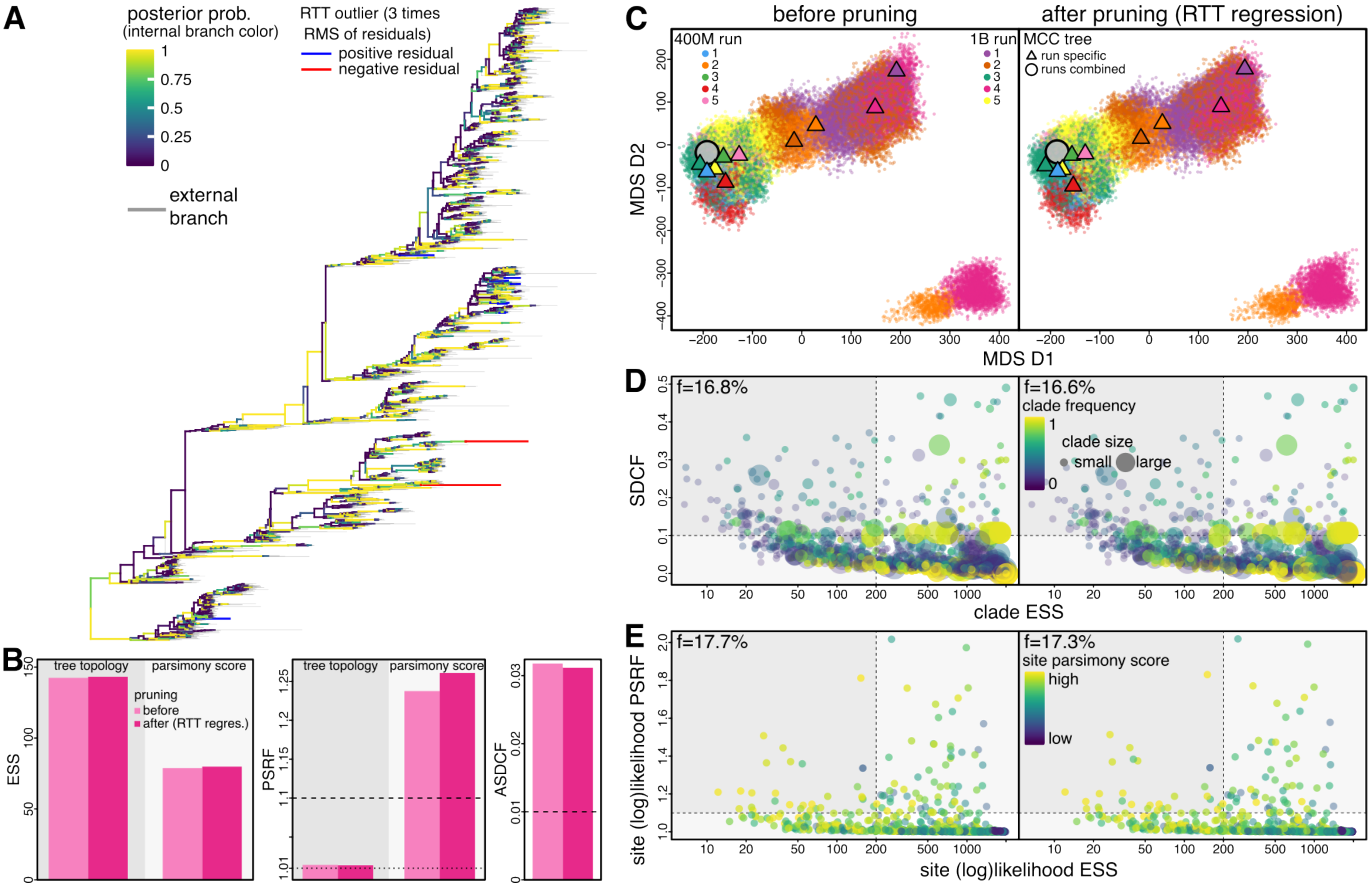


**Figure S20: Pruning off the RTT outlier sequences has little impact on MCMC performance for the influenza H1N1 virus dataset.** Note that for this dataset the number of sequences causing the poorly sampled clades revealed by the MCMC diagnosis are too many (≫10) to examine closely and a crude removal of all such sequences might result in a significantly reduced tree that loses part of its topology backbone, so we did not assess the impact of pruning off sequences based on the MCMC diagnostics here. **A)** MCC tree of the influenza H1N1 dataset (11) focusing on the problematic sequences identified as outliers in the RTT regression test (marked as orange diamond and blue subtending branches). **B)** Tree-based diagnostics that quantify MCMC performance before (light pink) and after (pink) the pruning corroborate the visual pattern shown in **C**. **C)** Posterior tree space before (Left) and after (Right) pruning the RTT outlier sequences from each sampled tree, with the color of each dot showing which MCMC chain the sampled tree is from. Each dot represents a sampled tree; a triangle represents the MCC tree summarized from an MCMC chain, while an open circle indicates the MCC tree summarized across all chains. **D)** Clade occurrence MCMC diagnostics before (Left) and after (Right) the pruning. SDCF ($y$-axis) quantifies the between-chain convergence and clade ESS ($x$-axis) quantifies the within-chain mixing performance, respectively. Each dot represents a clade, whose color indicates the clade frequency averaged across all replicates. **E)** Site-specific MCMC diagnostics before (Left) and after (Right) the pruning. The site-specific PSRF ($y$-axis) and ESS ($x$-axis) are computed using the log-likelihood of each site. Each dot represents a site, whose color indicates the parsimony number of mutations at that site over the corresponding tree.

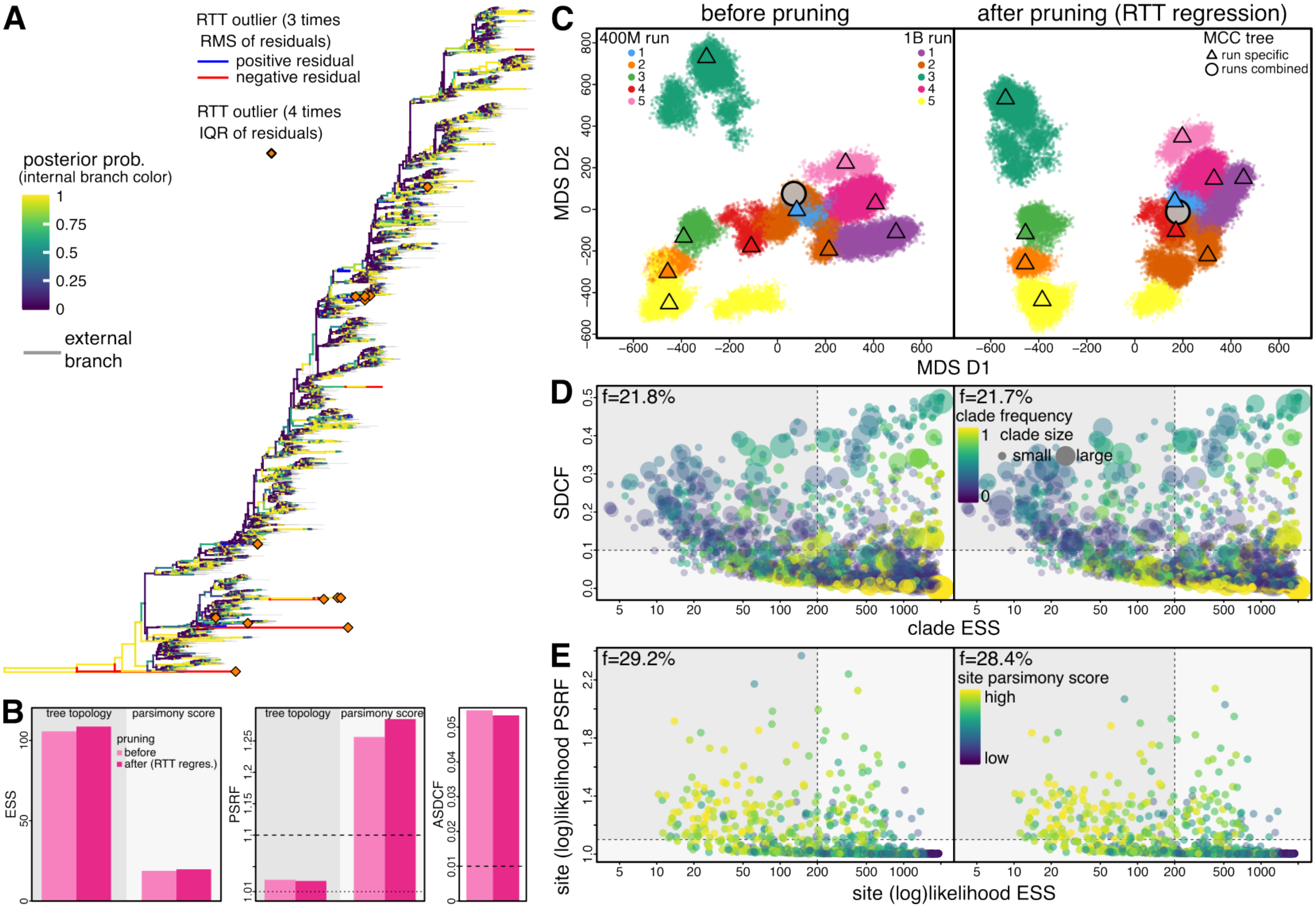


**Figure S21: Pruning off the RTT outlier sequences has little impact on MCMC performance for the influenza H3N2 virus dataset.** Note that for this dataset the number of sequences causing the poorly sampled clades revealed by the MCMC diagnosis are too many ($\gg$10) to examine closely and a crude removal of all such sequences might result in a significantly reduced tree that loses part of its topology backbone, so we did not assess the impact of pruning off sequences based on the MCMC diagnostics here. **A)** MCC tree of the influenza H3N2 dataset (11) focusing on the problematic sequences identified as outliers in the RTT regression test (marked as orange diamond and blue subtending branches). **B)** Tree-based diagnostics that quantify MCMC performance before (light pink) and after (pink) the pruning corroborate the visual pattern shown in **C**. **C)** Posterior tree space before (Left) and after (Right) pruning the RTT outlier sequences from each sampled tree, with the color of each dot showing which MCMC chain the sampled tree is from. Each dot represents a sampled tree; a triangle represents the MCC tree summarized from an MCMC chain, while an open circle indicates the MCC tree summarized across all chains. **D)** Clade occurrence MCMC diagnostics before (Left) and after (Right) the pruning. SDCF ($y$-axis) quantifies the between-chain convergence and clade ESS ($x$-axis) quantifies the within-chain mixing performance, respectively. Each dot represents a clade, whose color indicates the clade frequency averaged across all replicates. **E)** Site-specific MCMC diagnostics before (Left) and after (Right) the pruning. The site-specific PSRF ($y$-axis) and ESS ($x$-axis) are computed using the log-likelihood of each site. Each dot represents a site, whose color indicates the parsimony number of mutations at that site over the corresponding tree.

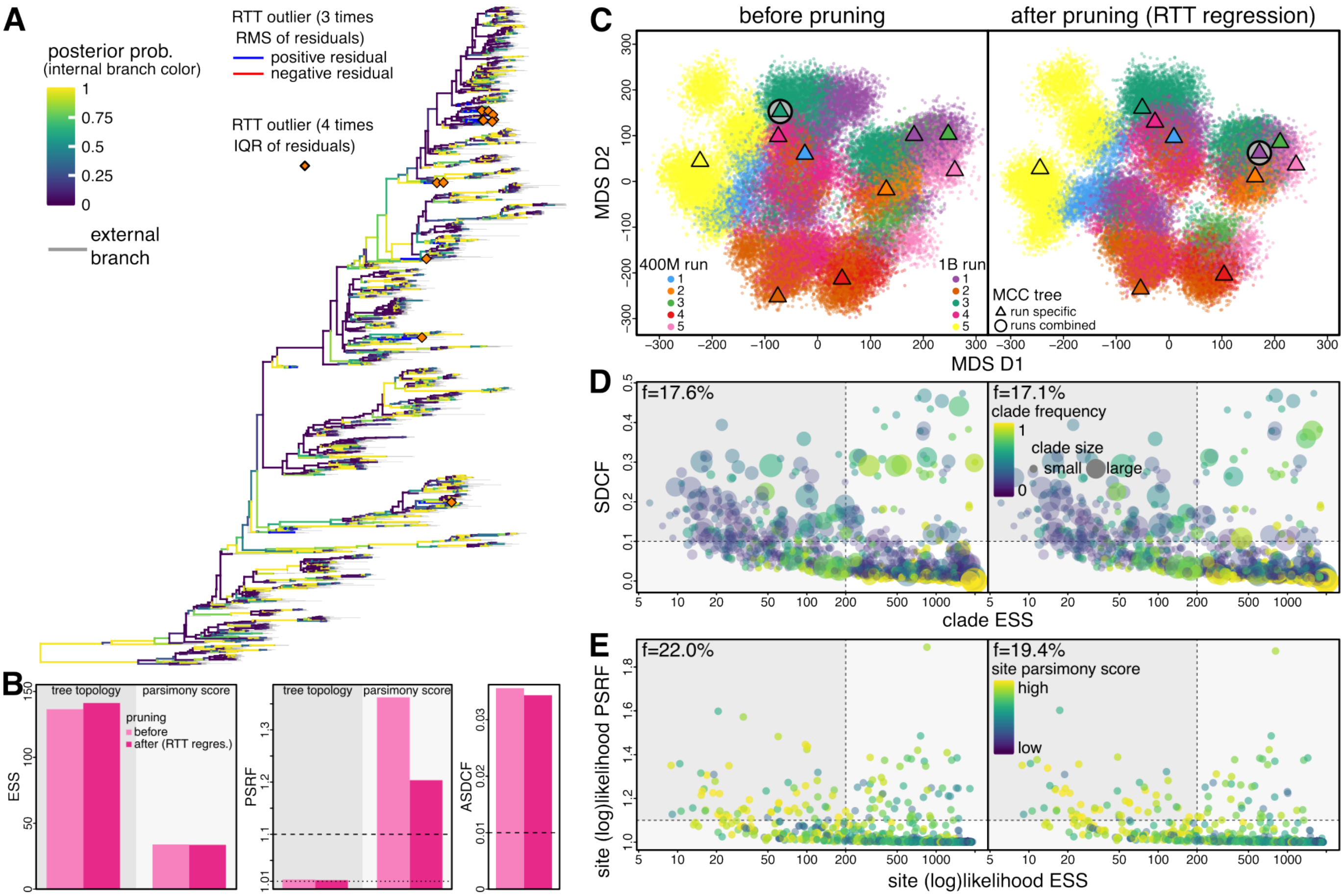


**Figure S22: Pruning off the RTT outlier sequences has little impact on MCMC performance for the influenza B/Vic virus dataset.** Note that for this dataset the number of sequences causing the poorly sampled clades revealed by the MCMC diagnosis are too many (≫10) to examine closely and a crude removal of all such sequences might result in a significantly reduced tree that loses part of its topology backbone, so we did not assess the impact of pruning off sequences based on the MCMC diagnostics here. **A)** MCC tree of the influenza B Victoria dataset (11) focusing on the problematic sequences identified as outliers in the RTT regression test (marked as orange diamond and blue subtending branches). **B)** Tree-based diagnostics that quantify MCMC performance before (light pink) and after (pink) the pruning corroborate the visual pattern shown in **C**. **C)** Posterior tree space before (Left) and after (Right) pruning the RTT outlier sequences from each sampled tree, with the color of each dot showing which MCMC chain the sampled tree is from. Each dot represents a sampled tree; a triangle represents the MCC tree summarized from an MCMC chain, while an open circle indicates the MCC tree summarized across all chains. **D)** Clade occurrence MCMC diagnostics before (Left) and after (Right) the pruning. SDCF ($y$-axis) quantifies the between-chain convergence and clade ESS ($x$-axis) quantifies the within-chain mixing performance, respectively. Each dot represents a clade, whose color indicates the clade frequency averaged across all replicates. **E)** Site-specific MCMC diagnostics before (Left) and after (Right) the pruning. The site-specific PSRF ($y$-axis) and ESS ($x$-axis) are computed using the log-likelihood of each site. Each dot represents a site, whose color indicates the parsimony number of mutations at that site over the corresponding tree.

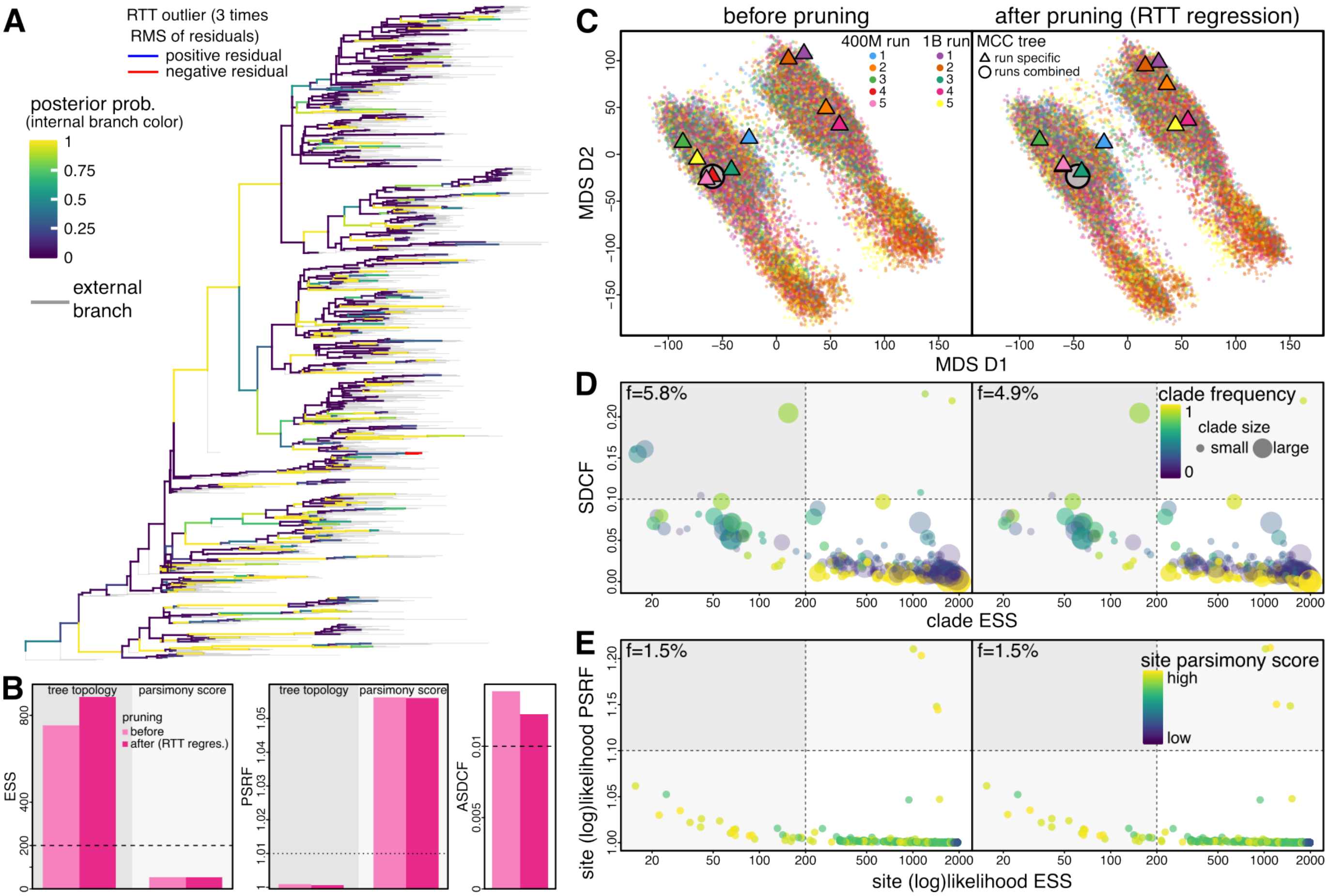


**Figure S23: Pruning off the RTT outlier sequences has little impact on MCMC performance for the SARS-CoV-2 Brazil dataset.** Note that for this dataset the number of sequences causing the poorly sampled clades revealed by the MCMC diagnosis are too many (≫10) to examine closely and a crude removal of all such sequences might result in a significantly reduced tree that loses part of its topology backbone, so we did not assess the impact of pruning off sequences based on the MCMC diagnostics here. **A)** MCC tree of the SARS-CoV-2 Brazil dataset (12) focusing on the problematic sequences identified as outliers in the RTT regression test (marked as orange diamond and blue subtending branches). **B)** Tree-based diagnostics that quantify MCMC performance before (light pink) and after (pink) the pruning corroborate the visual pattern shown in **C**. **C)** Posterior tree space before (Left) and after (Right) pruning the RTT outlier sequences from each sampled tree, with the color of each dot showing which MCMC chain the sampled tree is from. Each dot represents a sampled tree; a triangle represents the MCC tree summarized from an MCMC chain, while an open circle indicates the MCC tree summarized across all chains. **D)** Clade occurrence MCMC diagnostics before (Left) and after (Right) the pruning. SDCF ($y$-axis) quantifies the between-chain convergence and clade ESS ($x$-axis) quantifies the within-chain mixing performance, respectively. Each dot represents a clade, whose color indicates the clade frequency averaged across all replicates. **E)** Site-specific MCMC diagnostics before (Left) and after (Right) the pruning. The site-specific PSRF ($y$-axis) and ESS ($x$-axis) are computed using the log-likelihood of each site. Each dot represents a site, whose color indicates the parsimony number of mutations at that site over the corresponding tree.

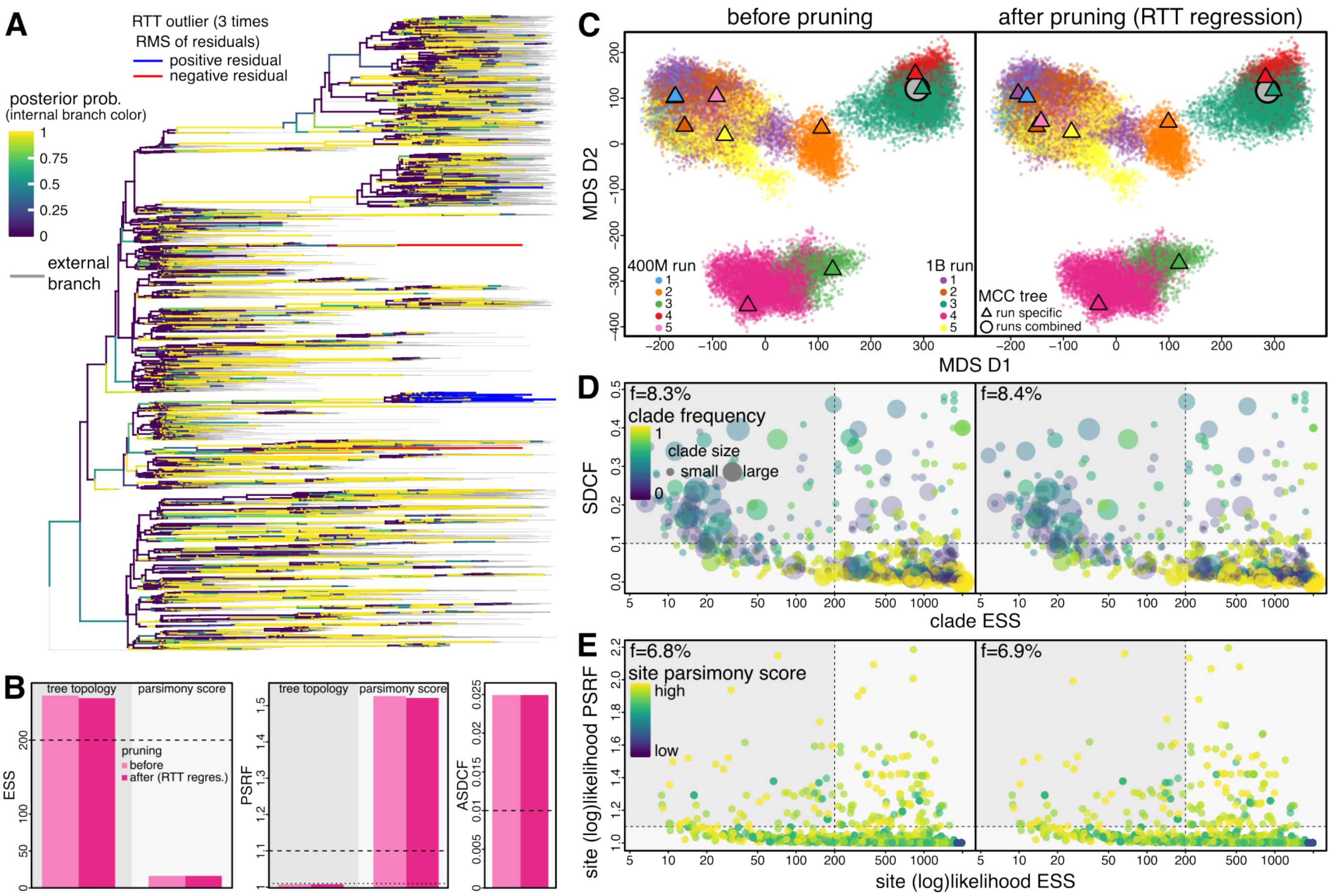


**Figure S24: Pruning off the RTT outlier sequences has little impact on MCMC performance for the SARS-CoV-2 Europe dataset.** Note that for this dataset the number of sequences causing the poorly sampled clades revealed by the MCMC diagnosis is too large (≫10) to examine closely and a crude removal of all such sequences might result in a significantly reduced tree that loses part of its topology backbone, so we did not assess the impact of pruning off sequences based on the MCMC diagnostics here. **A)** MCC tree of the SARS-CoV-2 Europe dataset (13) focusing on the problematic sequences identified as outliers in the RTT regression test (marked as orange diamond and blue subtending branches). **B)** Tree-based diagnostics that quantify MCMC performance before (light pink) and after (pink) the pruning corroborate the visual pattern shown in **C**. **C)** Posterior tree space before (Left) and after (Right) pruning the RTT outlier sequences from each sampled tree, with the color of each dot showing which MCMC chain the sampled tree is from. Each dot represents a sampled tree; a triangle represents the MCC tree summarized from an MCMC chain, while an open circle indicates the MCC tree summarized across all chains. **D)** Clade occurrence MCMC diagnostics before (Left) and after (Right) the pruning. SDCF ($y$-axis) quantifies the between-chain convergence and clade ESS ($x$-axis) quantifies the within-chain mixing performance, respectively. Each dot represents a clade, whose color indicates the clade frequency averaged across all replicates. **E)** Site-specific MCMC diagnostics before (Left) and after (Right) the pruning. The site-specific PSRF ($y$-axis) and ESS ($x$-axis) are computed using the log-likelihood of each site. Each dot represents a site, whose color indicates the parsimony number of mutations at that site over the corresponding tree.

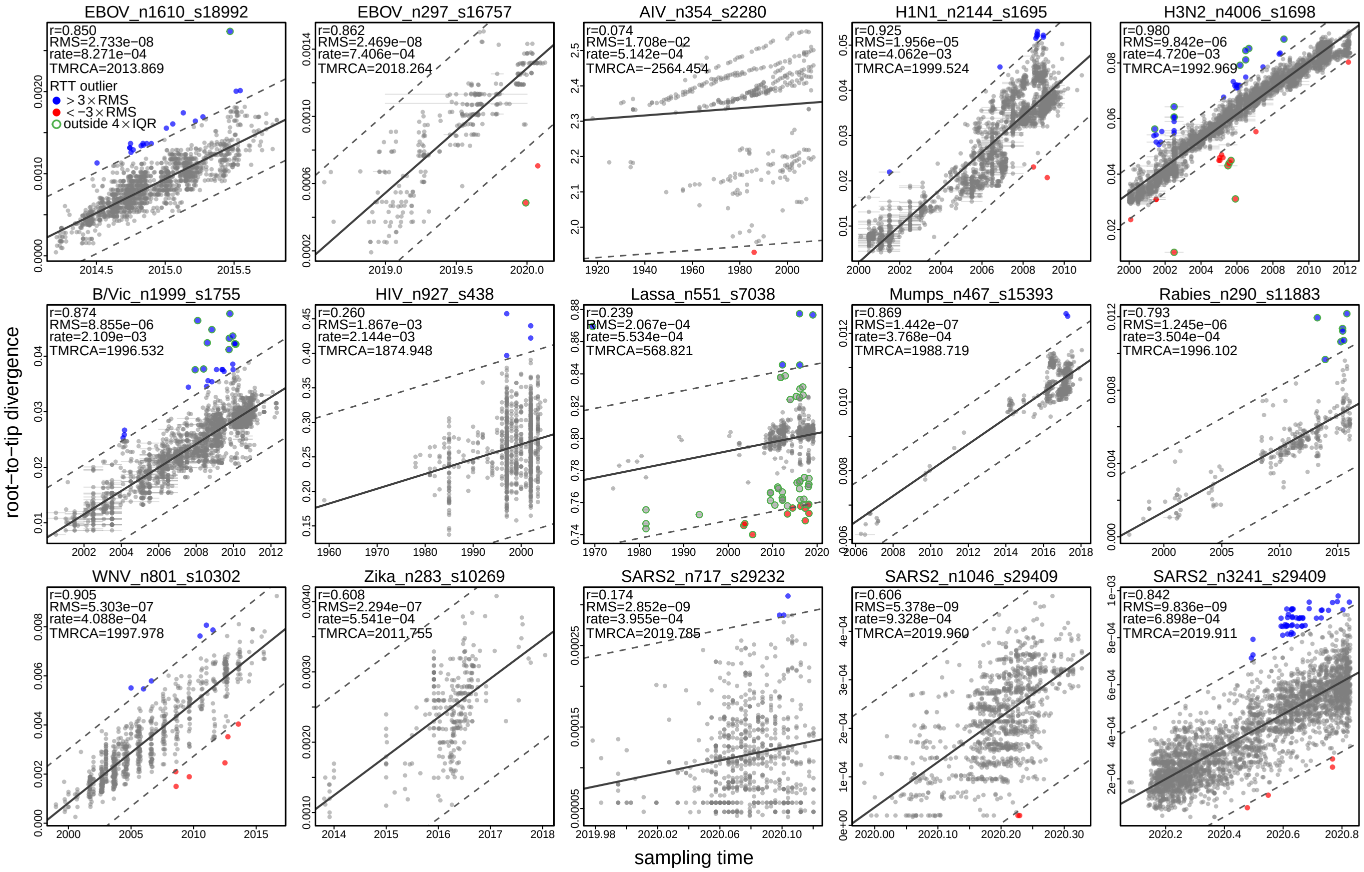


**Figure S25: Root-to-tip (RTT) regression test reveals strong temporal signal and outlier sequences in most datasets.** We perform linear regression between the sampling time of each tip ($x$-axis) and its genetic distance to the root ($y$-axis). Each panel corresponds to a dataset and each dot represents a tip; uncertainty in the sampling time of a tip sequence is illustrated by the associated horizontal bar. Results from the linear regression test are presented in the top left corner of each panel, including Pearson's correlation (r), root mean square of the residuals (RMS), inferred mutation rate (rate, which corresponds to the slope of the regression), and the inferred root time (tMRCA). For most datasets (with AIV_n354_s2280 as a clear exception), most dots fall along the regression line (solid), although for a handful of the datasets the correlation is rather weak. The tips whose residual fall outside three times of RMS (dashed lines) are colored (blue: positive; red: negative). Green circled dots indicate the tips whose absolute residual is greater than four times the interquartile range (IQR) of the residuals.

#### S1.5 Impact of Tree Sampling on Key Phylodynamic Conclusions

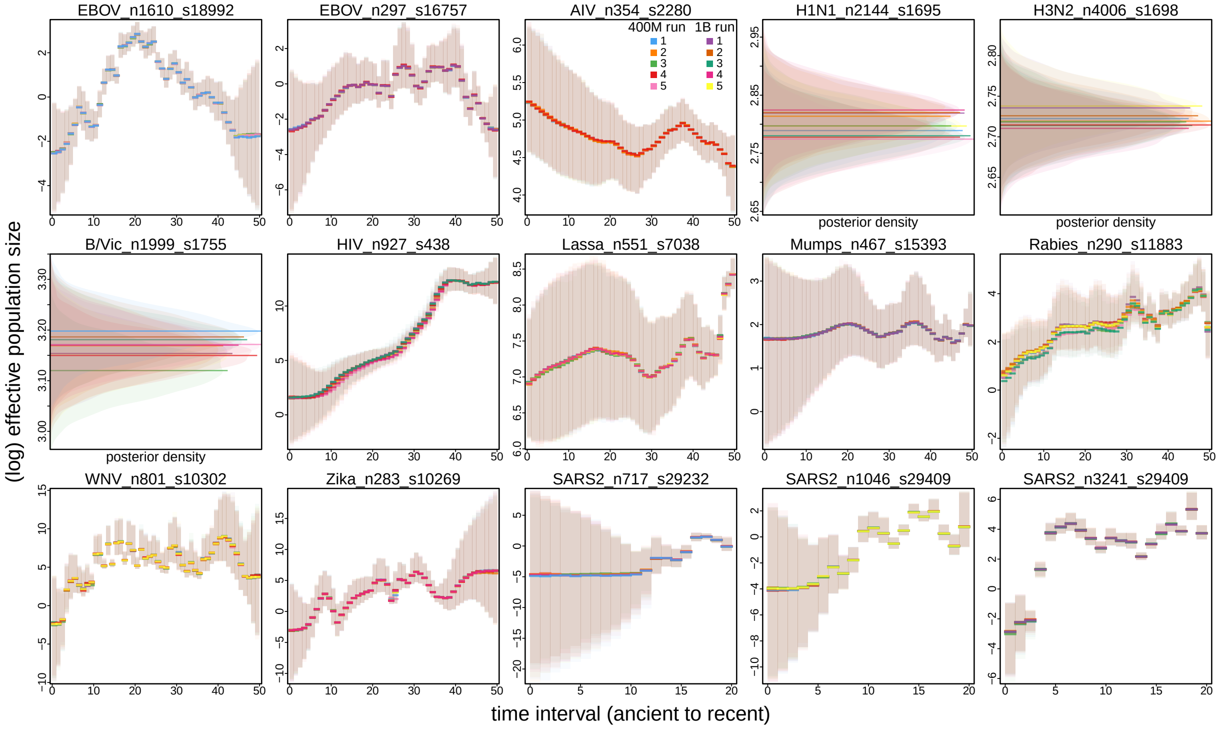


**Figure S26: Impact of tree sampling problems tends to be negligible on effective population size estimates.** Each panel depicts the (log) effective population size estimates through time (solid line: median; shaded region: 95% credible interval) of a dataset and the color indicates the corresponding MCMC chain index. For the three human influenza virus datasets that assumed a constant coalescent model in their analyses, we use the shaded region to represent its inferred posterior density. The demographic trajectories inferred under different replicates are barely distinguishable, except for a few datasets where it appears that the inferred curve is shifted to be slightly older or younger.

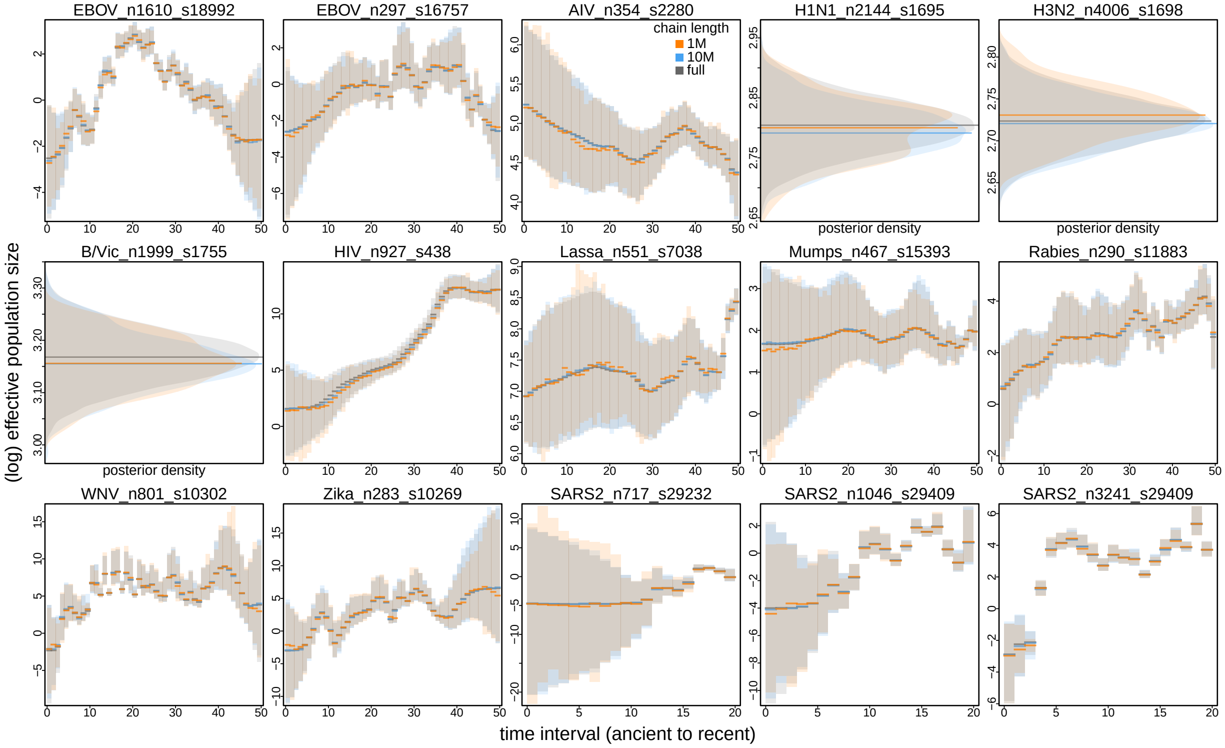


**Figure S27: Virtually identical accuracy and precision of effective population size estimates can be obtained with orders of magnitude shorter MCMC chains.** This figure is similar to Fig. S26 but focuses on the discrepancy between combined MCMC chains with different lengths instead of among replicate chains. The estimates (blue) summarized from combining the first ten million iterations of each of the ten replicate chains are virtually identical to those (gray) summarized from the full analyses, while the estimates (orange) summarized from combining the first one million iterations of each chain appear to be somewhat less accurate and precise. That is, effectively 2% of the full analyses would be sufficient to obtain the effective population size estimates.

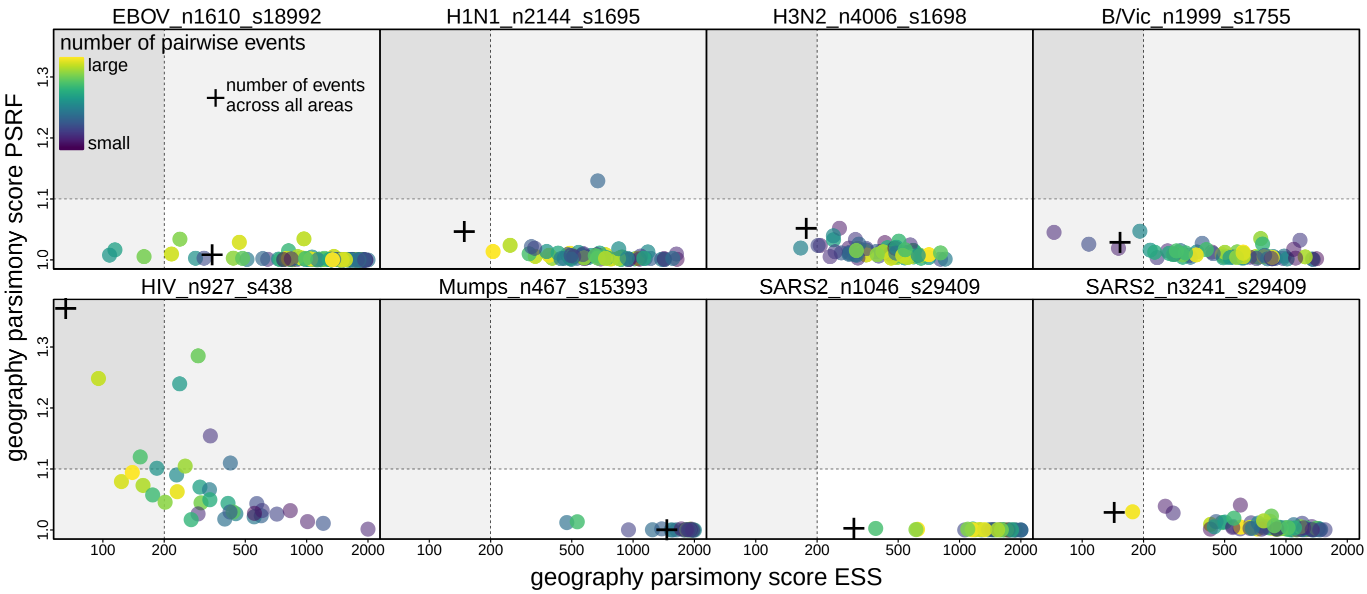


**Figure S28: The numbers of geographic events over most dispersal routes are sampled satisfactorily.** Each panel shows the MCMC diagnostics computed for the parsimony number of geographic events conditioned on each sampled tree in a dataset. (Here only the eight datasets for which the original study involved a discrete phylogeographic analysis and the geographic data are available are included.) Each dot represents a geographic dispersal route between a pair of geographic areas, whose color indicates the parsimony number of dispersal events on that particular route averaged over the posterior trees (yellow: large; purple: small); the cross indicates the total number of dispersal events among all geographic areas. (Here only the routes with at least one dispersal event occurring in at least 95% of the posterior samples are included.) PSRF ($y$-axis) quantifies convergence between the chains, and ESS ($x$-axis) quantifies mixing within each chain. Dashed lines indicate the thresholds typically used in phylogenetic studies; *i.e.*, PSRF above 1.1 (horizontal dashed line) indicates lack of convergence and ESS below 200 (vertical dashed line) indicates inadequate mixing. For most datasets, a small fraction of the dots reside outside the bottom right quadrant surrounded by the dashed lines, indicating that the numbers of dispersal events over those routes failed to be estimated reliably.

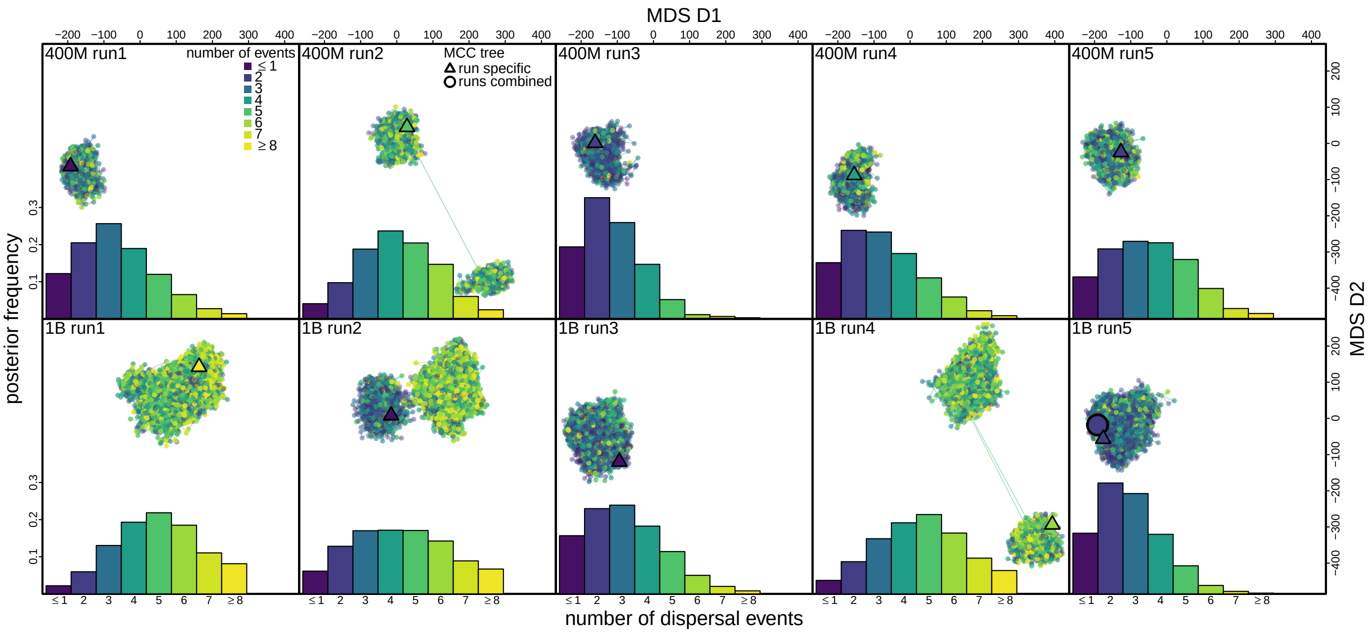


**Figure S29: Pronounced impact of tree sampling problems on influenza H1N1 virus dispersal patterns.** Here we focus on the number of dispersal events of influenza H1N1 virus inferred between two focal geographic areas (South America to the United States and Canada) in the study investigating the global dispersal dynamics of human influenza viruses (11). Each panel corresponds to an MCMC replicate, with the top row showing the posterior of each 400M run and the bottom row showing that of each 1B run. The dots in each panel visualize the MDS tree space; each dot represents a sampled tree, colored according to the number of parsimony geographic events over this particular dispersal route across that tree. A triangle represents the MCC tree summarized from an MCMC chain, while an open circle indicates the MCC tree summarized across all the replicates of that dataset. The histogram in each panel summarizes the distribution of the parsimony number of dispersal events across all the trees sampled in that MCMC chain. Large discrepancies—which appear to be caused by tree sampling problems—in the posterior estimates among these replicate chains exist: for some chains (*e.g.*, 400M run3) the median estimate is two while the 95% posterior interval includes no more than five events, whereas for some others (*e.g.*, 1B run1 or 1B run4) the median estimate is five.

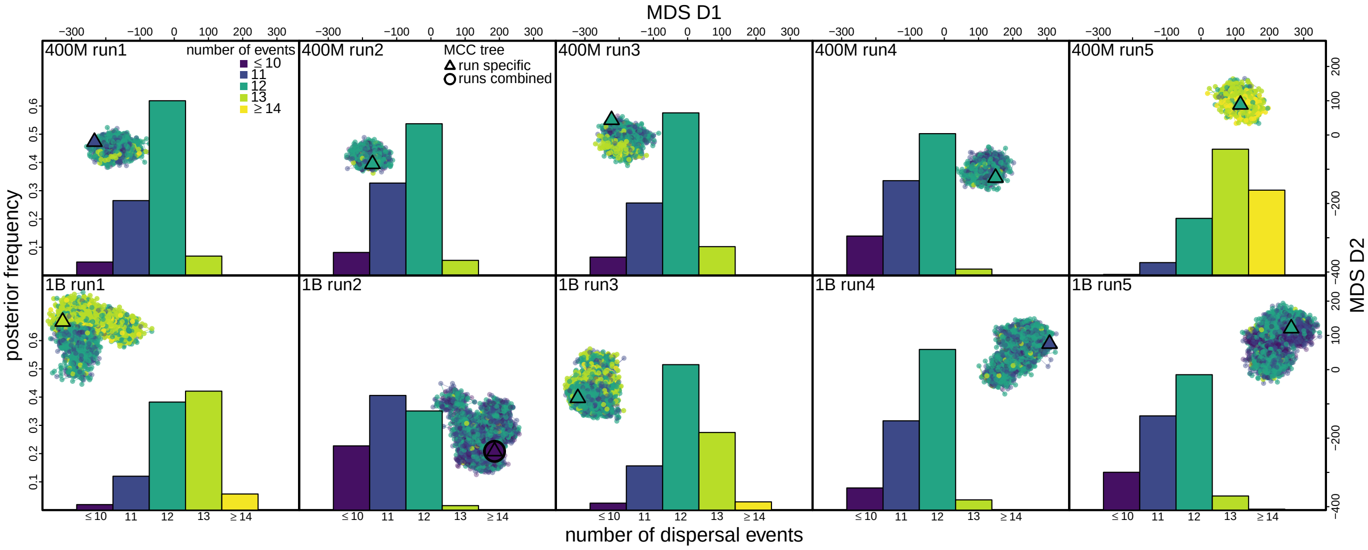


**Figure S30: Pronounced impact of tree sampling problems on geographic dispersal inferences for the HIV dataset.** Here we focus on the number of dispersal events inferred between two focal geographic areas (Kinshasa and Likasi) in the study investigating the early spread history of HIV (14). Each panel corresponds to an MCMC replicate, with the top row showing the posterior of each 400M run and the bottom row showing that of each 1B run. The dots in each panel visualize the MDS tree space; each dot represents a sampled tree, colored according to the number of parsimony geographic events over this particular dispersal route across that tree. A triangle represents the MCC tree summarized from an MCMC chain, while an open circle indicates the MCC tree summarized across all the replicates of that dataset. The histogram in each panel summarizes the distribution of the parsimony number of dispersal events across all the trees sampled in that MCMC chain. Large discrepancies—which appear to be caused by tree sampling problems—in the posterior estimates among these replicate chains exist: for some chains (*e.g.*, 1B run2) the median estimate is 11 and the 95% posterior interval is $[9, 12]$, whereas for some others (*e.g.*, 400M run5) the median estimate is 13 and the 95% posterior interval is $[11, 14]$.

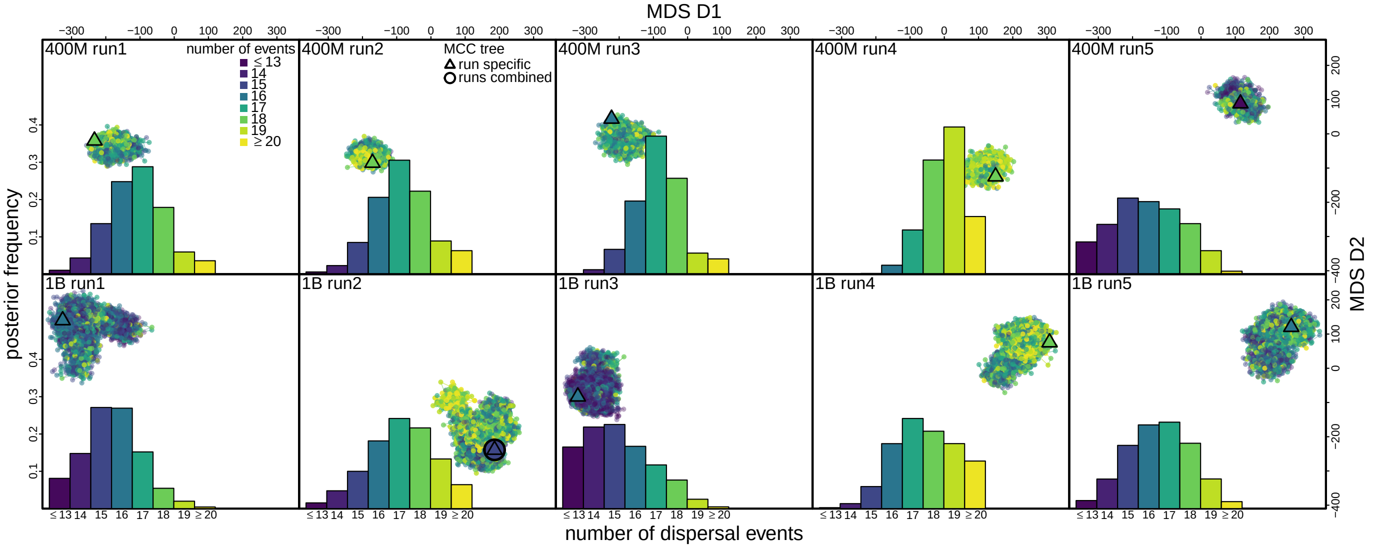

**Figure S31: Pronounced impact of tree sampling problems on geographic dispersal inferences for the HIV dataset (continued).** Here we focus on the number of dispersal events inferred between two focal geographic areas (Kinshasa and Bwamanda) in the study investigating the early spread history of HIV (14). Each panel corresponds to an MCMC replicate, with the top row showing the posterior of each 400M run and the bottom row showing that of each 1B run. The dots in each panel visualize the MDS tree space; each dot represents a sampled tree, colored according to the number of parsimony geographic events over this particular dispersal route across that tree. A triangle represents the MCC tree summarized from an MCMC chain, while an open circle indicates the MCC tree summarized across all the replicates of that dataset. The histogram in each panel summarizes the distribution of the parsimony number of dispersal events across all the trees sampled in that MCMC chain. Large discrepancies—which appear to be caused by tree sampling problems—in the posterior estimates among these replicate chains exist: for some chains (*e.g.*, 1B run3) the median estimate is 15 and the 95% posterior interval is [12, 18], whereas for some others (*e.g.*, 400M run4) the median estimate is 19 and the probability for there being fewer than 18 events is only 0.145. Note that this dispersal route is the reverse of that shown in Fig. 5; it appears that there are interactions between these two estimates (*e.g.*, 400M run4 infers the most dispersal events on this route from Kinshasa to Bwamanda while inferring the least on the opposite direction); *i.e.*, distinct tree topologies inferred across the MCMC replicates appear to affect the inferred direction of certain events over this dispersal route.

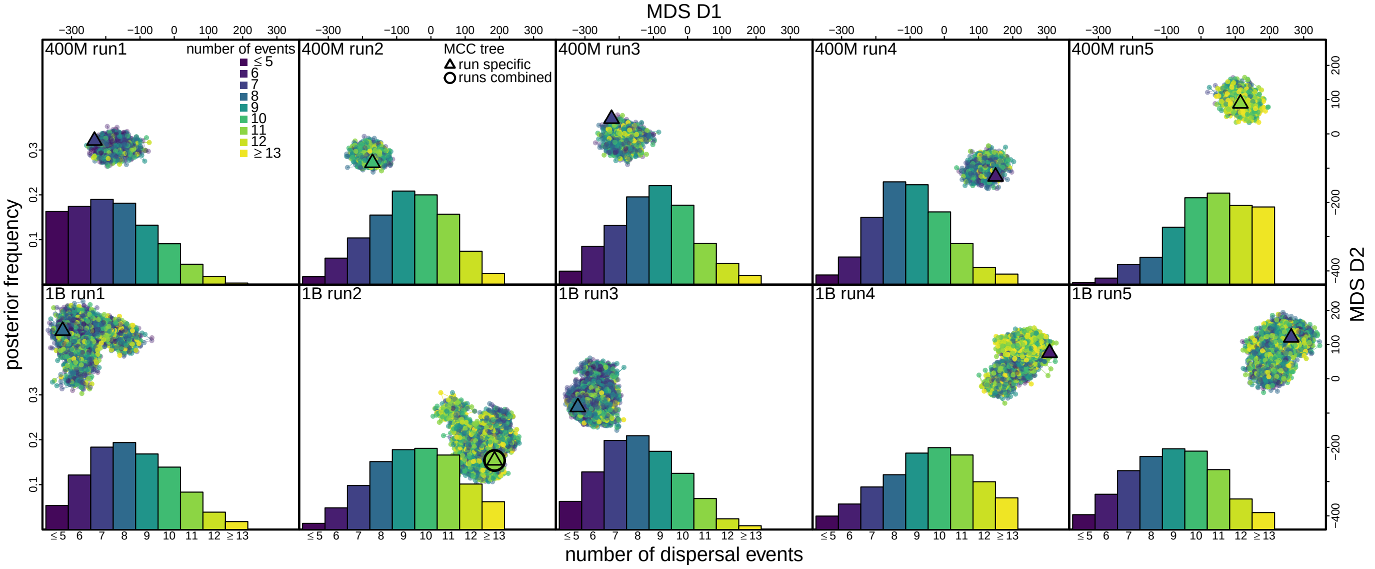

**Figure S32: Pronounced impact of tree sampling problems on geographic dispersal inferences for the HIV dataset (continued).** Here we focus on the number of dispersal events inferred between two focal geographic areas (Other C and Kinshasa) in the study investigating the early spread history of HIV (14). Each panel corresponds to an MCMC replicate, with the top row showing the posterior of each 400M run and the bottom row showing that of each 1B run. The dots in each panel visualize the MDS tree space; each dot represents a sampled tree, colored according to the number of parsimony geographic events over this particular dispersal route across that tree. A triangle represents the MCC tree summarized from an MCMC chain, while an open circle indicates the MCC tree summarized across all the replicates of that dataset. The histogram in each panel summarizes the distribution of the parsimony number of dispersal events across all the trees sampled in that MCMC chain. Large discrepancies—which appear to be caused by tree sampling problems—in the posterior estimates among these replicate chains exist: for some chains (*e.g.*, 400M run1) the median estimate is 7 and the 95% posterior interval is [4, 11], whereas for some others (*e.g.*, 400M run5) the median estimate is 11 and the 95% posterior interval is [7, 14].

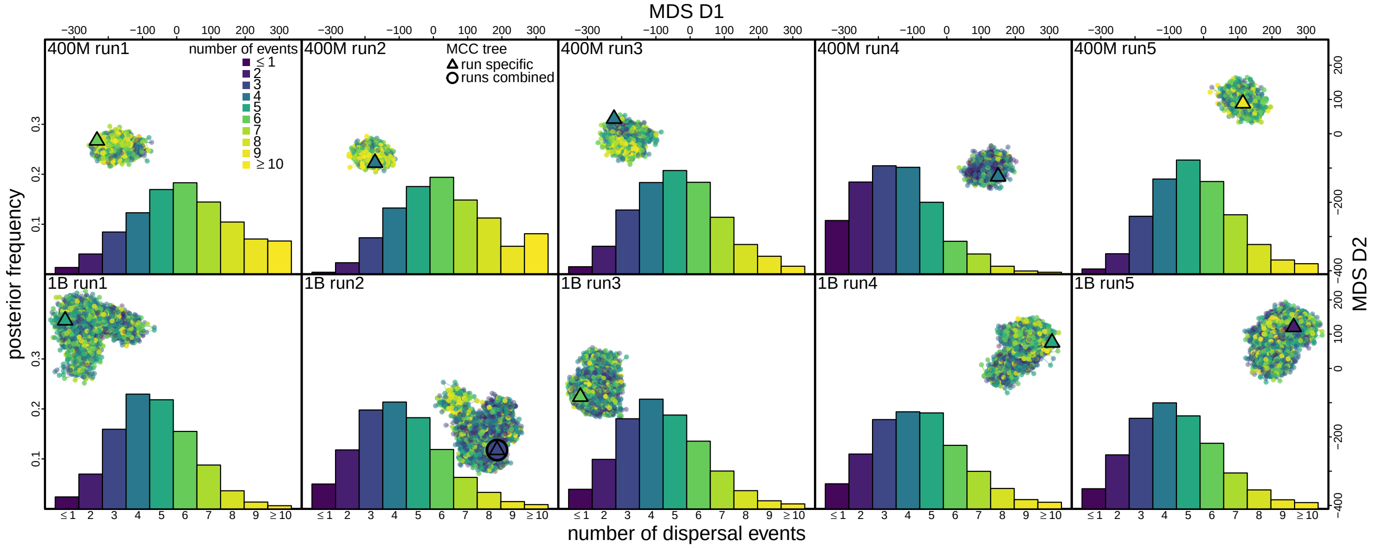


**Figure S33: Pronounced impact of tree sampling problems on geographic dispersal inferences for the HIV dataset (continued).** Here we focus on the number of dispersal events inferred between two focal geographic areas (Lubumbashi and Kinshasa) in the study investigating the early spread history of HIV (14). Each panel corresponds to an MCMC replicate, with the top row showing the posterior of each 400M run and the bottom row showing that of each 1B run. The dots in each panel visualize the MDS tree space; each dot represents a sampled tree, colored according to the number of parsimony geographic events over this particular dispersal route across that tree. A triangle represents the MCC tree summarized from an MCMC chain, while an open circle indicates the MCC tree summarized across all the replicates of that dataset. The histogram in each panel summarizes the distribution of the parsimony number of dispersal events across all the trees sampled in that MCMC chain. Large discrepancies—which appear to be caused by tree sampling problems—in the posterior estimates among these replicate chains exist: for some chains (*e.g.*, 400M run4) the median estimate is 3 and the 95% posterior interval is $[1, 7]$, whereas for some others (*e.g.*, 400M run2) the median estimate is 6 and the 95% posterior interval is $[3, 11]$.

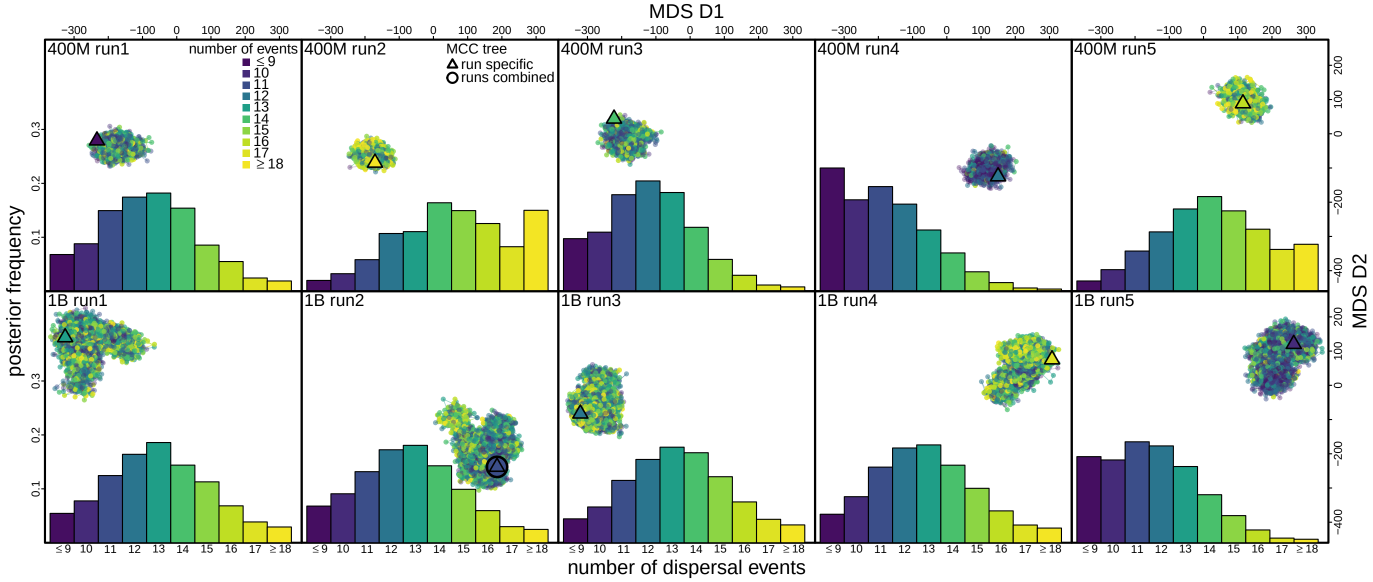


**Figure S34: Pronounced impact of tree sampling problems on geographic dispersal inferences for the HIV dataset (final).** Here we focus on the number of dispersal events inferred between two focal geographic areas (Mbuji-Mayi and Kinshasa) in the study investigating the early spread history of HIV (14). Each panel corresponds to an MCMC replicate, with the top row showing the posterior of each 400M run and the bottom row showing that of each 1B run. The dots in each panel visualize the MDS tree space; each dot represents a sampled tree, colored according to the number of parsimony geographic events over this particular dispersal route across that tree. A triangle represents the MCC tree summarized from an MCMC chain, while an open circle indicates the MCC tree summarized across all the replicates of that dataset. The histogram in each panel summarizes the distribution of the parsimony number of dispersal events across all the trees sampled in that MCMC chain. Large discrepancies—which appear to be caused by tree sampling problems—in the posterior estimates among these replicate chains exist: for some chains (*e.g.*, 400M run4) the median estimate is 11 and the 95% posterior interval is $[7, 15]$, whereas for some others (*e.g.*, 400M run2) the median estimate is 15 and the 95% posterior interval is $[10, 20]$.

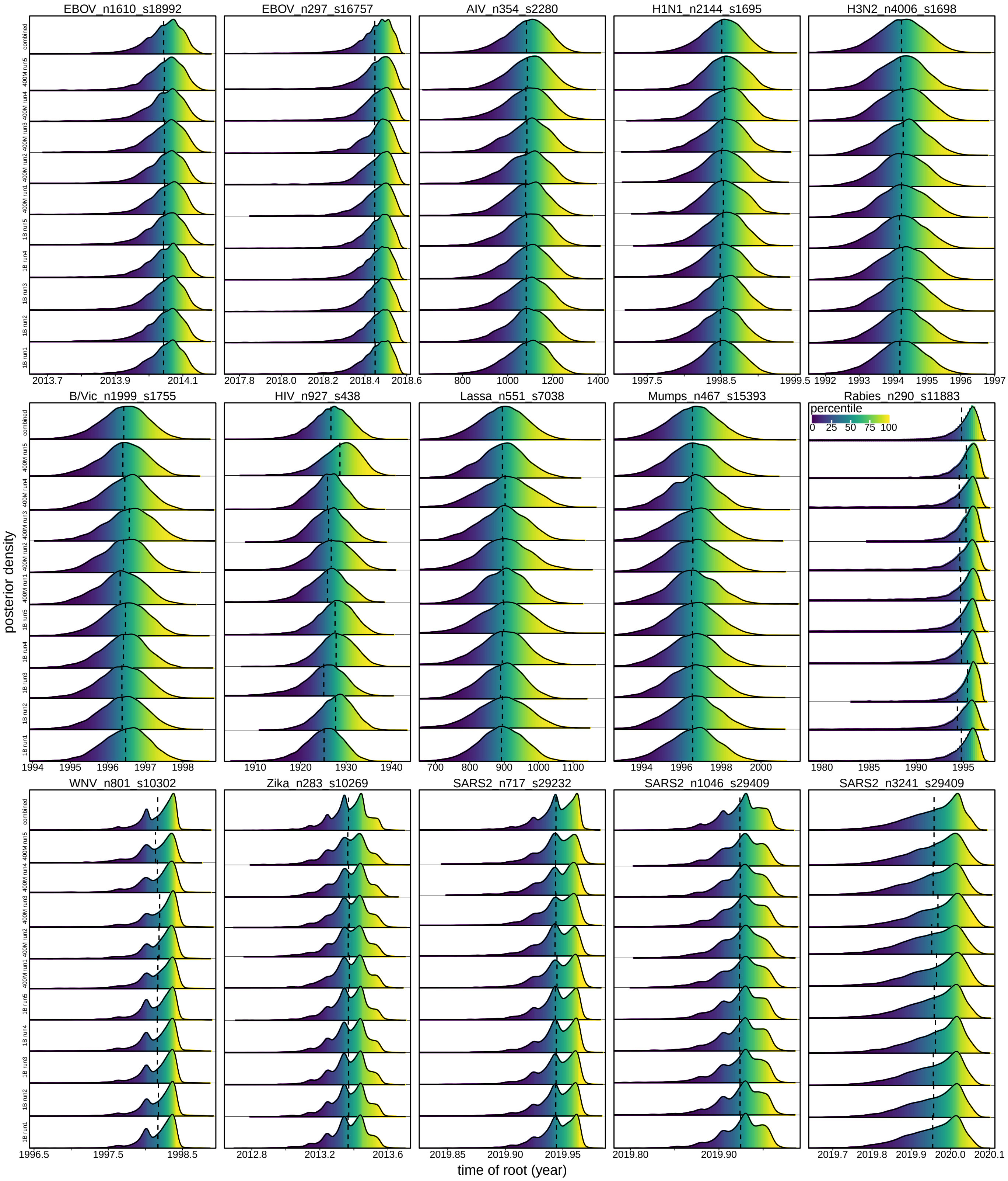


**Figure S35: Minor impact of tree sampling problems on root age estimates.** Posterior distribution of the root age inferred from each MCMC chain and all chains combined. Each panel corresponds to a dataset, and the 11 rows in each panel correspond to the combined chain (top) and the 10 replicate chains. Each distribution is divided into vertical bins where each bin is colored according to the percentile (see inset legend bar) of the posterior distribution summarized across all chains in which that bin's $x$-axis value falls, aiding the comparison between the inferred distributions across the chains. The vertical dashed line on each distribution indicates the mean estimate.

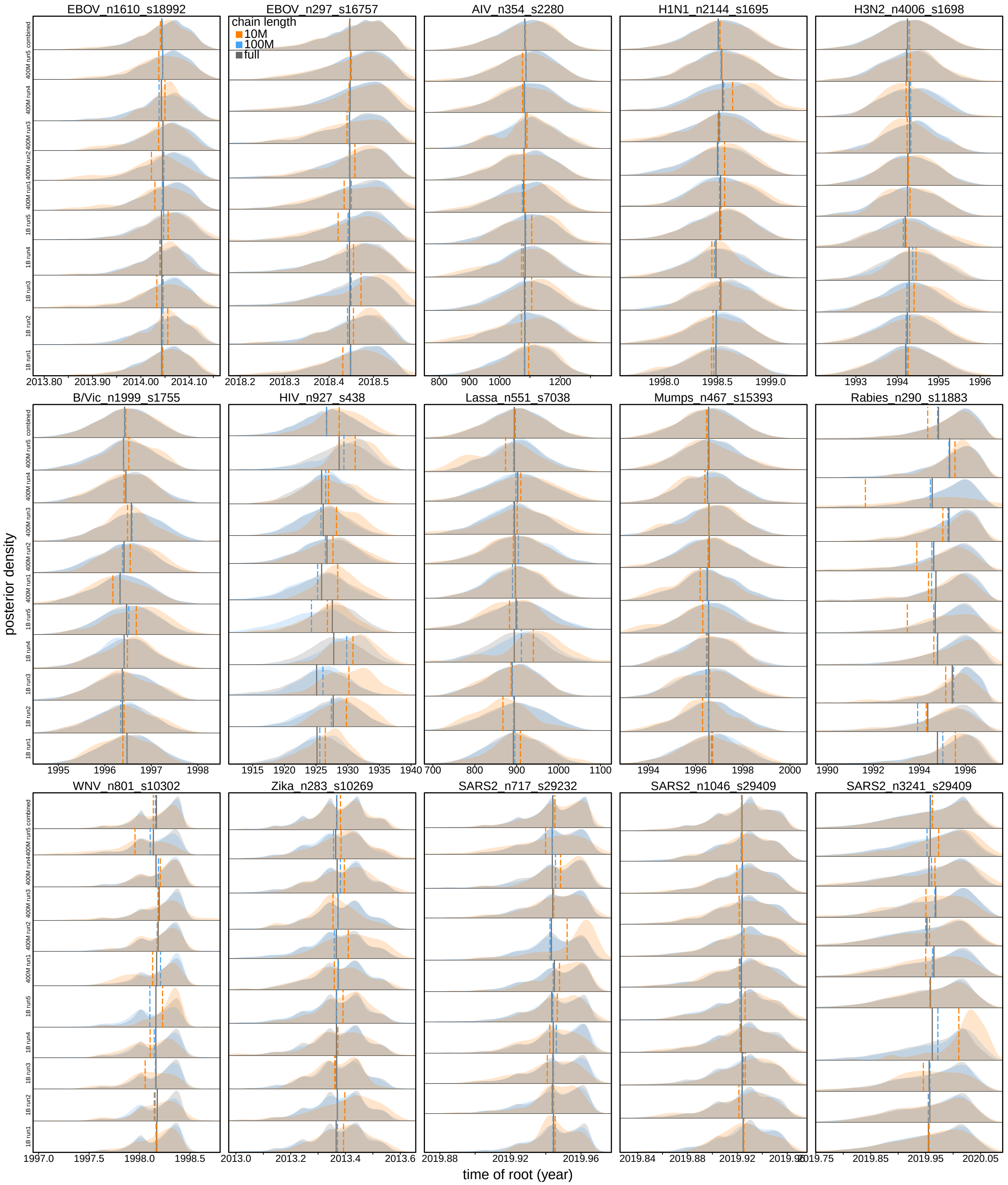


**Figure S36: Similar accuracy and precision of root age estimates can be obtained with shorter (but not too short) MCMC chains.** This figure is similar to Fig. S35 but focuses on the discrepancy between combined MCMC chains with different lengths instead of among replicate chains. The inferred distributions (blue) summarized from the first 100 million iterations of each replicate chain are barely distinguishable from those (gray) summarized from the full analyses, whereas the distributions (orange) summarized from the first 10 million iterations are noticeably distinguishable.

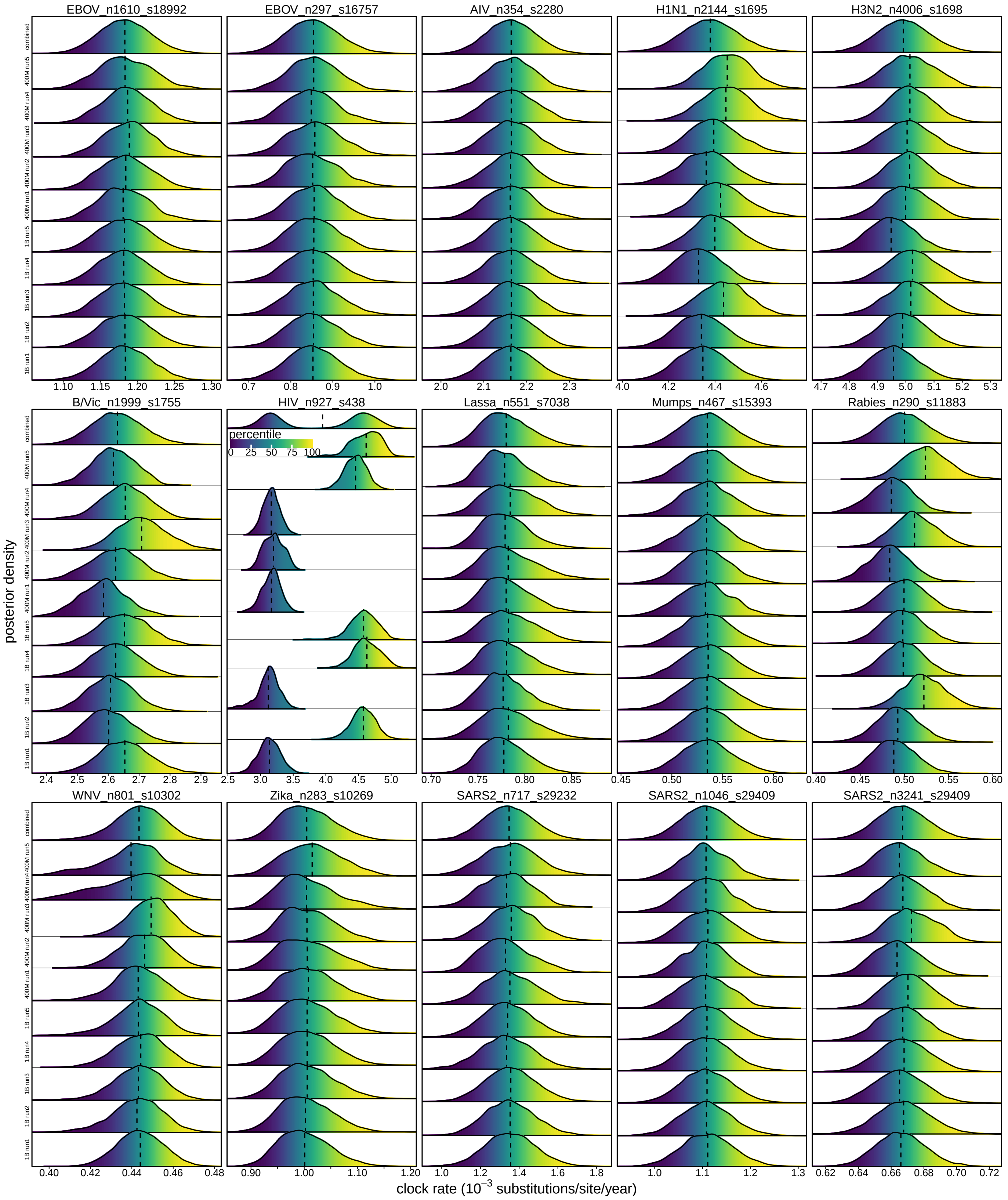


**Figure S37: Noticeable impact of tree sampling problems on mean evolutionary rate estimates.** Posterior distribution of the evolutionary rate inferred from each MCMC chain and all chains combined. Each panel corresponds to a dataset, and the 11 rows in each panel correspond to the combined chain (top) and the 10 replicate chains. Each distribution is divided into vertical bins where each bin is colored according to the percentile (see inset legend bar) of the posterior distribution summarized across all chains in which that bin's $x$-axis value falls, aiding the comparison between the inferred distributions across the chains. The vertical dashed line on each distribution indicates the mean estimate.

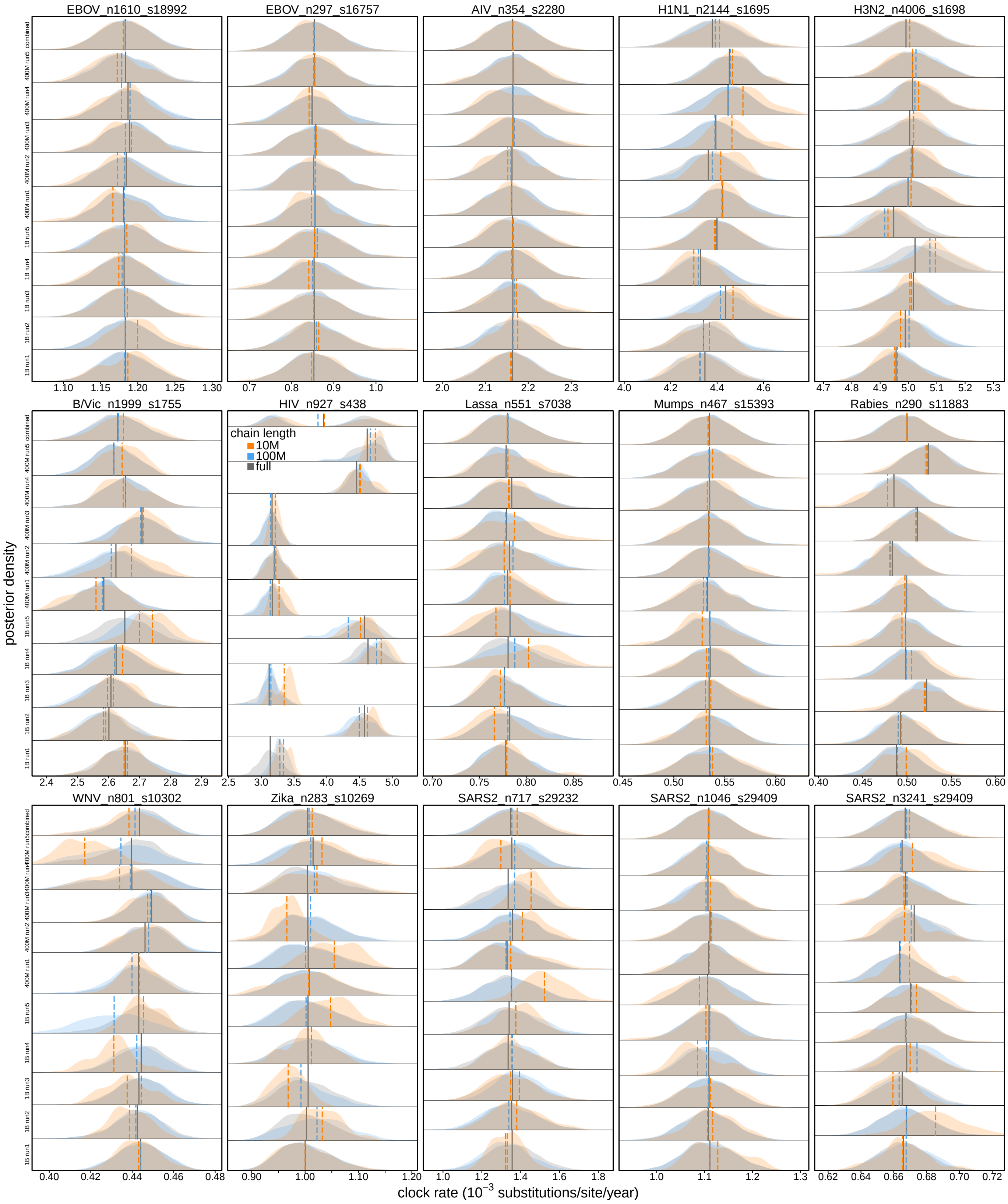


**Figure S38: Similar accuracy and precision of mean evolutionary rate estimates can be obtained with shorter (but not too short) MCMC chains.** This figure is similar to Fig. S37 but focuses on the discrepancy between combined MCMC chains with different lengths instead of among the replicate chains. The inferred distributions (blue) summarized from the first 100 million iterations of each replicate chain are barely distinguishable from those (gray) summarized from the full analyses, whereas the distributions (orange) summarized from the first 10 million iterations are significantly different.

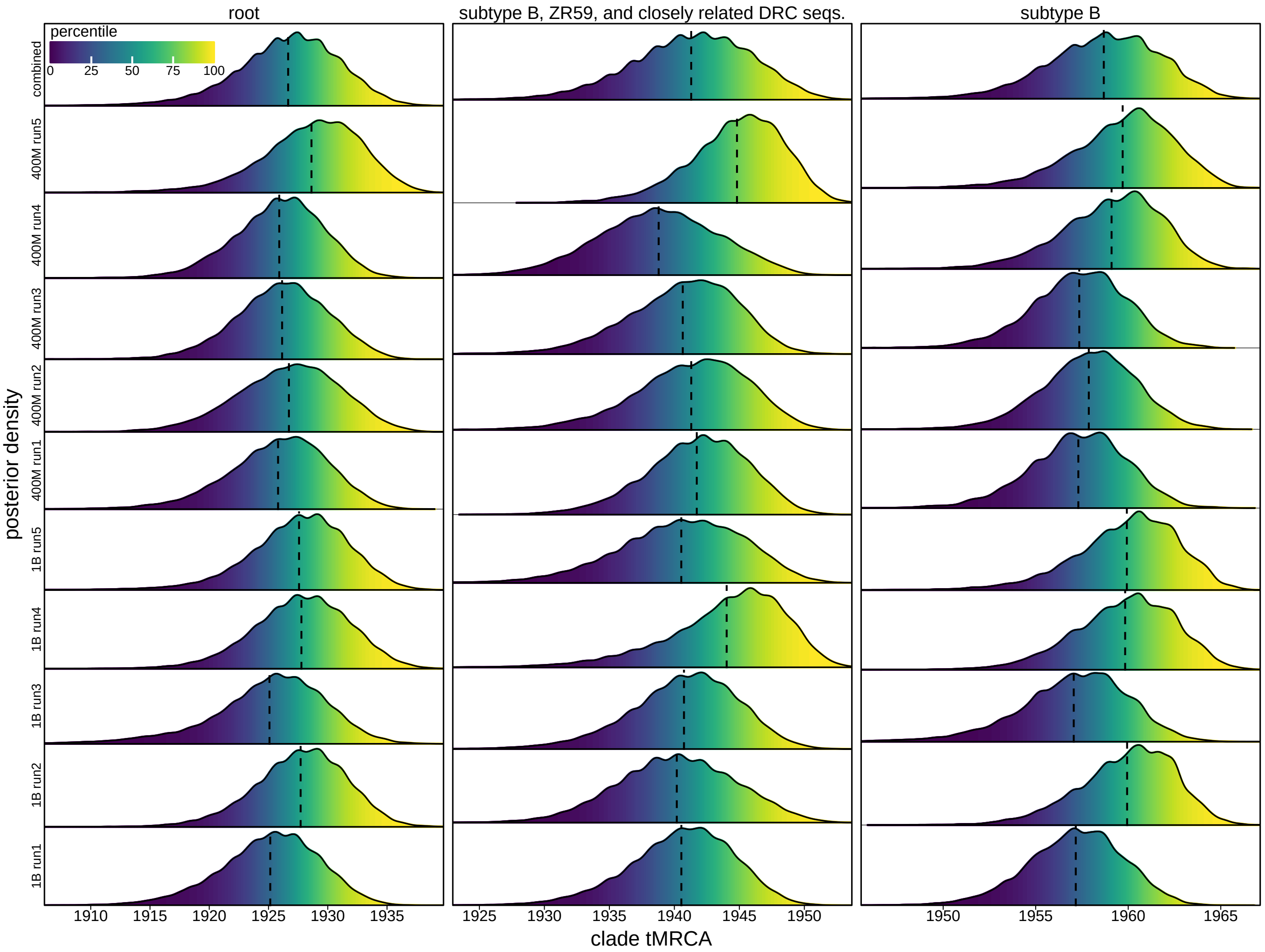


**Figure S39: Impact of tree sampling problems can be pronounced for clade tMRCA and origin time estimates.** Posterior distribution of the tMRCA of several focal clades in the HIV dataset (14). Each column corresponds to a focal clade and each row shows the posterior distribution inferred from the corresponding MCMC chain. Each distribution is divided into vertical bins where each bin is colored according to the percentile (see inset legend bar) of the posterior distribution of that parameter summarized across all replicates in which that bin's *x*-axis value falls, highlighting the discrepancy in the time estimates across the replicate chains. The vertical dashed line on each distribution indicates the mean estimate, further revealing the discrepancy across chains.

### S2 Analyses of Empirical Datasets

#### S2.1 General Analysis Protocol

##### S2.1.1 Data Curation

We collected 15 classic viral datasets from published empirical phylodynamic studies (see Table S1), performed extensive Bayesian phylodynamic analyses of each dataset with `BEAST` v1.10.5 (15), and then analyzed the posterior outputs. For all but the SARS-CoV-2 datasets, we directly obtained the sequence data as well as the associated sampling times from the `BEAST` XML scripts of each dataset provided in its published repository. For the studies that performed geographic phylodynamic inference to investigate viral dispersal dynamics, we extracted the sampling location of each viral sample from their `BEAST` XML scripts. For the SARS-CoV-2 datasets, as the sequence data were not available in the corresponding XML scripts, we acquired these sequences from GISAID (16) using their accession IDs provided by the original study. This sequence downloading process was automated by using the `GISAIDR` (17) package in `R`. We aligned the downloaded SARS-CoV-2 sequences using `MUSCLE` v5 (18) and removed the 5′ and 3′ UTRs at the two ends of the alignment, resulting in 29409 nucleotide sites for each SARS-CoV-2 alignment. To mitigate the impact of missing data in our phylodynamic inference given the limited information in the sequence data, we filtered each SARS-CoV-2 alignment to exclude the viral samples with too many missing or gap sites, resulting in an alignment with 1046 sequences for the SARS-CoV-2 Brazil dataset (12) and an alignment with 3241 sequences for the SARS-CoV-2 Europe dataset (13). Given the extremely limited genetic diversity in the SARS-CoV-2 Origin dataset (6), we further truncated its alignment by only keeping the coding region, and applied additional filtration steps. In particular, we translated the nucleotide alignment of this dataset into an amino-acid alignment using the `seqinr` package (19) in `R` to identify sequences with premature stop codons and then excluded these sequences. We also discarded sequences with more than 30 ambiguous sites in the nucleotide alignment or more than 10 ambiguous sites in the translated amino-acid sequence. These steps led to a final alignment with 717 sequences and 29232 nucleotide sites for the SARS-CoV-2 Origin dataset.

**Table S1: Empirical dataset information.**

| Study | Virus | Dataset | ID | $n$ | $s$ | $s'$ |
|---|---|---|---|---|---|---|
| Dudas et al. (10) | Ebola | West Africa | EBOV_n1610_s18992 | 1610 | 18992 | 10210 |
| Mbala-Kingebeni et al. (20) | Ebola | DRC relapse | EBOV_n297_s16757 | 297 | 16757 | 1906 |
| Worobey et al. (21) | Avian influenza | PB2 | AIV_n354_s2280 | 354 | 2280 | 1261 |
| Bedford et al. (11) | Human Influenza | H1 Large | H1N1_n2144_s1695 | 2144 | 1695 | 1286 |
| | | H3 Large | H3N2_n4006_s1698 | 4006 | 1698 | 1353 |
| | | Vic Large | B/Vic_n1999_s1755 | 1999 | 1755 | 1092 |
| Faria et al. (14) | HIV | B | HIV_n927_s438 | 927 | 438 | 435 |
| Klitting et al. (22) | Lassa | L | Lassa_n551_s7038 | 551 | 7038 | 6119 |
| Moncla et al. (7) | Mumps | — | Mumps_n467_s15393 | 467 | 15393 | 4116 |
| Viana et al. (8) | Rabies | — | Rabies_n290_s11883 | 290 | 11883 | 2275 |
| Dellicour et al. (9) | West Nile | — | WNV_n801_s10302 | 801 | 10302 | 3654 |
| Grubaugh et al. (23) | Zika | — | Zika_n283_s10269 | 283 | 10269 | 4136 |
| Pekar et al. (6) | SARS-CoV-2 | Origin | SARS2_n717_s29232 | 717 | 29232 | 737 |
| Candido et al. (12) | SARS-CoV-2 | Brazil | SARS2_n1046_s29409 | 1046 | 29409 | 1975 |
| Lemey et al. (13) | SARS-CoV-2 | Europe | SARS2_n3241_s29409 | 3241 | 29409 | 8565 |

$n$ denotes the number of sequences, $s$ denotes the number of sites in the sequence alignment, and $s'$ denotes the number of site patterns.

##### S2.1.2 Bayesian Phylodynamic Analysis Setup

In our phylodynamic analyses of these empirical datasets, we specified models and priors, as well as the MCMC proposals and their weights, following the source XML files; the proposals and weights are mostly identical to the defaults in `BEAUti` v1.10.5 under the specified models, reflecting standard empirical practices. For each dataset, we ran 10 independent MCMC replicate chains using `BEAST` v1.10.5 with `BEAGLE` v3.2.0 (24) enabled, including five chains with 400 million iterations each and the other five with one billion iterations each, and then discarded the first 200 million iterations from each chain as the burnin. We extended `BEAST` to record all tree moves attempted during MCMC, including the branches involved at each move, their distance in the tree, and whether the move was accepted. Each chain was sampled every 100 thousand iterations to log the continuous model variables as well as the tree to files. We set up the 400-million-iteration chains ("400M runs") to reflect best practices in most empirical studies, while the excessively long 1-billion-iteration chains ("1B runs") aim to establish the "ground-truth" posterior tree distribution. We specified a conservatively long burnin (200 million iterations) to focus on the post-burnin MCMC behavior.

##### S2.1.3 Analysis Post-processing and MCMC Diagnosis

We characterize the MCMC performance and explore the biological consequences of such performance by examining three types of variables sampled in or summarized from the posterior outputs of each MCMC chain, including: (1) continuous model parameters (*e.g.*, evolutionary rate) and statistics (*e.g.*, joint posterior density); (2) clade-specific (*e.g.*, node age), branch-specific (*e.g.*, branch rate) and site-specific (*e.g.*, site likelihood) variables; and (3) the tree.

*Compute MCMC Diagnostics for Continuous Variables*

For each continuous variable, we assess its mixing performance within each MCMC chain by the effective sample size (ESS), which was computed from the post-burnin samples with an `R` implementation of `Tracer`'s (25) ESS calculation routine provided by the `essTracer` function in the `convenience` package (26); we then took the mean ESS values across all chains to represent the mixing performance of each dataset and report it throughout this study. To assess the convergence behavior across the MCMC replicate chains, we computed the potential scale reduction factor (PSRF; 27) with the `Rhat` function (28) available in the `rstan` (29) `R` package. We also computed ESS using the `effectiveSize` function provided by `R` package `CODA` (30) and the `ess_bulk` function available in `rstan`, as well as PSRF using `CODA`'s `gelman.diag` function, to ensure that the calculated values with various methods are consistent. We used the thresholds that are commonly adopted in Bayesian phylogenetic studies to determine the sampling adequacy within chain (ESS $> 200$) and convergence between chains (PSRF $< 1.1$).

As the MCMC replicate chains we ran vary in length (five chains with 400 million iterations and the other five with a billion iterations), we pursued two subsampling schemes to allow single ESS and PSRF values to be computed across all the chains for each dataset. The first is the truncation scheme, where we took the first 400 million iterations from each of the 1B runs so that all the 10 chains end up with identical length. The MCMC diagnostics computed under this truncation scheme are reported throughout this study. For the second scheme, we computed another set of the MCMC diagnostics with just the five 1B runs, demonstrating how ESS increases when the chains run longer. The results under these two schemes are qualitatively congruent, and thus those diagnostics computed under the second scheme are not shown here.

*Summarize Clade- and Site-specific Variables and Compute MCMC Diagnostics for These Variables*

The clade- and site-specific variables are not directly available in the posterior outputs `BEAST` generates by default, so we post-process `BEAST` outputs to obtain them.

*Clade-specific MCMC diagnostics.* We used the `TreeSummary` tool provided by `BEAST` to generate the clade-specific variable samples in a post-hoc manner. Specifically, we first processed the (post-burnin) trees sampled by each MCMC chain with `TreeSummary` to generate a list of clades whose frequency is at least 10% in that chain, and then combined such lists across all replicates. We then provided that combined clade list as an input to `TreeSummary` and ran it again to extract the value of the desired variables associated with each clade obtained from the sampled trees, including the node age, subtending branch length, and branch-specific evolutionary rate (for the datasets that assume a relaxed-clock model).

The variables associated with the clades whose posterior probability equals one are treated identically with the other continuous model variables with respect to calculating the MCMC diagnostics. For the clades that do not appear in all the sampled trees, we conducted additional truncation when computing PSRF to ensure that the number of samples included for each chain is identical (*i.e.*, for each clade, each chain is truncated according to the chain with the fewest meaningful samples of that clade); when computing ESS of a variable associated with such a clade, we took the average ESS across the chains weighted by the number of meaningful samples (*i.e.*, the number of sampled trees containing that clade) from each chain. We further excluded the clades that are inferred to have a posterior probability below 0.1 in more than half of the chains to avoid extreme ESS or PSRF values caused by such insufficient samples. For the clade-specific variables whose MCMC samples fail to mix or converge according to the corresponding MCMC diagnostics, and whose clade is decisively supported (*i.e.*, posterior probability above 0.9), we also plotted the inferred posterior distributions to visually examine the impact of such MCMC failure.

*Clade ESS and SDCF.* Apart from the clade-specific variables, we also compute MCMC diagnostics for the clade occurrence itself by treating it as a binary variable whose value is one when the clade is present in a sampled tree and zero otherwise (26). The clade occurrence samples were similarly fetched from the posterior trees using `TreeSummary`. For each clade that is inferred with a posterior probability above 0.01 ("detected") in at least half of the replicates, we calculated its ESS in each of the replicates in which that clade is detected and then took the average across those replicates to be the mean ESS of that clade. For a replicate chain where the clade is inferred with posterior probability 1, we assigned its chain-specific ESS to be the largest finite ESS among all the clades sampled in that chain.

We then evaluate the convergence of clade occurrence across chains using the standard deviation of clade frequency (SDCF). Average standard deviation of split frequencies (ASDSF; 31) is a commonly used diagnostic in Bayesian phylogenetics that assesses the topological convergence among replicate chains, obtained by first calculating the standard deviation of the estimated frequency (*i.e.*, posterior probability) of a given split (SDSF) among the chains and then taking the mean of the SDSF across splits whose mean split frequency is above 0.1. An ASDSF below 0.01 is generally taken as very good indication of convergence, whereas a value above 0.05 is deemed to indicate lack of convergence (32). The SDCF diagnostic we use in this study is identical to the SDSF in terms of the way it is computed and assessed, except that here we deal with clades in rooted trees instead of splits in unrooted trees.

*Site-specific MCMC diagnostics.* In addition to the tree and continuous variables that `BEAST` logs by default, we edited the XML scripts manually to record additional variables throughout the MCMC, such as the log-likelihood of each site (*i.e.*, site likelihood). We also used the `pml` function available in `R` (33) package `phangorn` (34) to compute the likelihood of each site given each MCMC sample; these two ways of computing the site likelihood produce effectively identical values. To compute the site-specific parsimony score, we used the `parsimony` function available in `phangorn`. The branch-specific parsimony score was instead computed by first calling the `ancestral.pars` function in `phangorn` to reconstruct the parsimonious mutation history for each site given the sampled tree, and then for each branch summarizing the parsimony number of mutations across all sites in the alignment. When multiple mutation histories that are equally maximally parsimonious exist at a site on a sampled tree, we randomly sampled one such history and summarize the branch-specific parsimony scores based on it. We excluded the trivial branches and sites whose parsimony score is identical across all MCMC samples from our MCMC diagnosis.

*Geographic dispersal inference and the corresponding MCMC diagnostics.* For the datasets where the geographic location of each viral sample is available, we assessed the impact of tree convergence on phylogeographic inference by counting the parsimony numbers of geographic dispersal events across all areas as well as between each pair of areas. Similar to the site-specific parsimony score, we obtained the total number of parsimony dispersal events on each sampled tree via `phangorn`'s `parsimony` function. The number of pairwise parsimony events was computed similarly by first reconstructing the parsimonious geographic history given each sampled tree in the posterior via `phangorn`'s `ancestral.pars` function, and then counting the number of events between each pair of areas in the reconstructed history. We only include the pairwise dispersal routes whose mean number of events is above 0.5 in all the replicate chains in our MCMC diagnosis to avoid extreme values due to insufficient samples. We treated these dispersal parsimony scores as continuous variables for computing ESS and PSRF.

*Characterize Posterior Tree Space and Compute Tree-Based MCMC Diagnostics*

We characterize the posterior tree space visually and quantitatively by three types of summaries, including: (1) a consensus summary tree, (2) a two-dimension representation of the tree space produced by multidimensional scaling (MDS), and (3) numeric MCMC diagnostics focusing on tree (*e.g.*, tree ESS and tree PSRF). To produce a summary tree of each dataset, we generated a maximum clade credibility (MCC) tree from the sampled posterior trees combined across all replicate chains using `TreeAnnotator` v1.10.5 (4); we also generated an MCC tree for each individual chain. In addition, we explored two alternative methods developed recently for summarizing posterior trees: HIPSTR (4) and CCD0-MAP (5). We used `TreeAnnotator` v1.10.5 for inferring the HIPSTR trees and `TreeAnnotator` v2.7.8 (35) for inferring the CCD0-MAP trees. To generate HIPSTR and CCD0-MAP trees within reasonable time and memory constraints (a few days and several hundred gigabytes per dataset), we thinned the posterior to one tree per 100 thousand iterations when collecting clades and computing frequencies. For datasets where CCD0-MAP analyses would require substantially longer, we used its approximation algorithm (ACCD0-MAP).

*Compute pairwise tree distances.* Both the tree-space visualization and tree MCMC diagnostics require a matrix of distances between each pair of trees in the posterior as the input; we computed both the pairwise Robinson-Foulds (RF) (36) and rooted subtree prune and regraft (rSPR) distances. Specifically, we calculated the pairwise RF distance between rooted trees by counting the number of incompatible clades (instead of splits); the list of clades in each tree was obtained by running the `prop.part` function provided by `R` package `ape` (37). The pairwise rSPR distances were computed using `rspr` (1, 2), a stand-alone C++ program; for all but five smaller datasets (including EBOV_n297_s16757, AIV_n354_s2280, Lassa_n551_s7038, Rabies_n290_s11883, and Zika_n283_s10269), we resorted to the `-approx` option to compute the approximate instead of exact rSPR distances due to the prohibitive computation time.

*Tree space visualization with MDS.* To visualize the posterior distribution of trees for a more intuitive presentation of the topological exploration, we conducted dimensionality reduction on each of the computed RF and rSPR pairwise distance matrices using multidimensional scaling (MDS) (38). We employed various MDS approaches to ensure that the generated two-dimension tree space is visually consistent among those approaches and thus insensitive to the selected MDS approach; these approaches include classical MDS [*i.e.*, principal coordinates analysis (PCoA)] via `R`'s `cmdscale` function, metric MDS via the `smacofSym` function from the `smacof` `R` package (39), and non-metric MDS via the `isoMDS` function from the `MASS` `R` package (40). We report the visualization results under the classical MDS approach in this study as these approaches lead to qualitatively similar results.

*Tree ESS and PSRF.* We quantitatively evaluated the single-chain mixing performance with tree ESS statistics (3) and multi-chain convergence behavior with tree PSRF (41) based on the computed pairwise distance matrices. Tree PSRF is a topological Gelman–Rubin-like convergence diagnostic developed by Whidden and Matsen (2015) (41), effectively comparing the within- and between-chain mean square distances. We implemented a function to calculate the tree PSRF following the description provided by Whidden and Matsen (2015).

We considered six different tree ESS measures in this study, including: (1) the Fréchet Correlation ESS (`frechetCorrelationESS`), (2) the median pseudo-ESS (`medianPseudoESS`), (3) the minimum pseudo-ESS (minPseudoESS), (4) the approximate ESS (approximateESS), (5) the classical MDS ESS (CMDSESS), and (6) a multivariate MDS ESS (MultiMDSESS). The frechetCorrelationESS, medianPseudoESS, minPseudoESS, approximateESS were computed via the corresponding functions provided by the treess R package (3); CMDSESS was computed by treating the first dimension of the two-dimension MDS matrix as a univariate continuous variable. Detailed descriptions of these ESS measures (other than that of the approximateESS) are available in Magee et al. (2024) (3), while the description of the approximateESS can be found in Lanfear et al. (2016) (42). Finally, we calculated MultiMDSESS by performing 10-dimension MDS (instead of the regular two-dimension MDS) on the distance matrix first, treating each row of the resulting 10-d MDS matrix as a multivariate variable, and computing an ESS of this multivariate variable using the multiESS function of the mcmcse R package (43). We confirmed that these tree ESS measures (other than the approximateESS; see results) show consistent pattern across the datasets, and thus, as recommended by Magee et al. (2024), we used the frechetCorrelationESS to represent tree ESS in our main results.

We also compared the visualization outputs and the tree MCMC diagnostics obtained from the two distance metrics (RF and rSPR) to ensure that our results are robust to the distance metric selection. Indeed, the results under the two distance metrics are qualitatively similar, and thus we focused on the results under the more commonly used RF distance throughout this manuscript.

*Visualize and Validate Tree Landscape Ruggedness*

To visualize tree landscape ruggedness, we overlaid the posterior density value associated with each sampled tree on the MDS plot. Specifically, we discretized the two-dimensional RF-based MDS tree space into a grid and then color each grid cell by the mean posterior density value of the tree samples in that cell. To reduce noise, the mean density value of each cell was computed with up to 25 highest-density samples in that cell (approximately half of the average number of samples per cell).

To validate observed valleys, for two datasets (Lassa and Mumps), we performed additional phylodynamic analyses with topology fixed at each position along the valley-crossing path and compared posterior densities across these fixed topologies. Specifically, for each of these datasets, we ran four independent 25-million-iteration MCMC chains using BEAST v1.10.5 with BEAGLE v3.2.0 enabled, discarding the first 1 million iterations from each chain as the burnin. We used fewer and shorter chains than the original analyses as the sampling was much easier with fixed topology. Similar to the original analyses, we computed MCMC diagnostics to ensure satisfactory sampling for these fixed-topology analyses.

*Identify Problematic Sequences and Assess Their Impact*

The branch-specific MCMC diagnostics—in particular the branch-length ESS and PSRF—revealed that only a small fraction of tips are typically associated with convergence issues, despite the apparent lack of convergence indicated by the inferred disjoint distributions in tree space. For five of the datasets (including Lassa_n551_s7038, Mumps_n467_s15393, Rabies_n290_s11883, WNV_n801_s10302, and SARS2_n717_s29232) where a limited number of sequences fail to converge according to their associated MCMC diagnostics, we verified whether these identified sequences are the direct causes of the tree sampling problems by pruning them from each of the sampled trees and re-assessing the tree sampling performance. Specifically, we used TreePruner, a tool provided by BEAST that takes the inferred posterior trees as its input, to prune the identified problematic sequences from each posterior tree, and then re-computed the clade- and site-specific MCMC diagnostics conditioning on these pruned trees.

In addition, we reanalyzed two datasets (Lassa and Rabies) with BEAST after prospectively removing the identified problematic sequences from the alignments. For each of these datasets, we ran 10 independent MCMC chains with the other settings following the original 400M runs. We assessed MCMC performance, visualized the posterior tree space, and summarized phylodynamic parameter estimates for these prospective removal analyses, and compared the results with those from the original analyses.

For the datasets for which many more sequences are identified to be problematic (despite still being a small fraction, *e.g.*, $\approx$ 10% of all the tips), we were unable to determine the causal sequences to prune based upon MCMC diagnosis because a stable backbone topology cannot be established from the sampled trees. Instead, we attempted to identify sequences that should have been excluded from the phylodynamic analyses by applying the root-to-tip (RTT) regression test (44), a data-quality test that is commonly used prior to a phylodynamic analysis as a preliminary check to assess the temporal signal in the genetic data and identify outlier sequences whose sampling dates or genetic composition is strongly incongruent with the rest. The RTT regression effectively performs a linear regression between the genetic distance from each tip to the root and the sampling time of each tip, intended to be used as a qualitative exploration of the data instead of a formal statistical test (44) due to the partial correlation between the RTT genetic distances (resulting from the shared ancestry in the phylogeny).

To estimate the genetic distances, we first conducted maximum-likelihood (ML) phylogenetic inference of each dataset using IQ-TREE (45) to obtain a point estimate of the unrooted phylogram. We ran 20 replicate IQ-TREE analyses (with four perturbation strength values, including 0.025, 0.05, 0.1, 0.2, specified via the `-pers` argument and five replicates under each value) for each dataset and then took the inferred tree from the run whose ML score is the highest among all replicates as the ML unrooted phylogram. We then rooted the inferred ML tree by minimizing the root mean squared residuals (RMS), using `ape`'s (37) `rtt` function (with the `objective` argument of `rtt` set to "rms"). Finally, we performed a linear regression between the estimated RTT distances and the corresponding sampling times via the `lm` function in `R`. We also conducted the above last two steps of RTT regression, including rooting the phylogram and performing the linear regression, with the `TempEst` program (44), where the tree was rooted under the "residual mean squared" option. We confirmed that the results of these two approaches were virtually identical and proceeded to build our analysis pipeline with functions in `R` for easier downstream automation.

As the RTT regression is not a valid statistical hypothesis test (44), we identified outlier sequences based on commonly used rule-of-thumb cutoffs, including three times the root mean squared residuals (RMS) and four times the interquartile range (IQR; *i.e.*, the $25^{\text{th}}$ to $75^{\text{th}}$ percentile) as specified by the program `treetime` (46). Similar to how we treated the problematic sequences identified based on MCMC diagnosis, we pruned the identified outliers from the posterior trees inferred from each dataset and then re-computed the relevant MCMC diagnostics. We only conducted that RTT-outlier pruning to the datasets fitting the following three conditions: (1) at least one outlier is identified in the RTT test; (2) tree sampling problems have been identified; and (3) the RTT regression demonstrated temporal signal (*i.e.*, Pearson's r above 0.6 between the genetic divergence and sampling time). This resulted in nine empirical datasets to examine, including EBOV_n1610_s18992, H1N1_n2144_s1695, H3N2_n4006_s1698, B/Vic_n1999_s1755, Mumps_n467_s15393, Rabies_n290_s11883, WNV_n801_s10302, SARS2_n1046_s29409, and SARS2_n3241_s29409.

To explore the potential for recombination tests to detect the problematic sequences, we also analyzed each dataset with two recombination detection methods, `GARD` (47) and `3SEQ` (48). `GARD` is a computationally intensive likelihood-based model-selection method that involves repeated phylogenetic inferences under various recombination scenarios; we ran the `GARD` analyses with HyPhy v2.5 (49) under the default settings. For four of the 15 datasets, `GARD` was unable to run as these alignments are shorter than what `GARD` requires (and the minimum number of sites required by `GARD` increases with the number of sequences); for most other datasets, the `GARD` analyses took at least several days to two weeks to complete on an HPC cluster, thus rendering its computational costs similar to those of the actual Bayesian phylodynamic analysis, and the `GARD` analyses of two datasets failed to complete within the 30-day cluster walltime. As `GARD` is designed to test whether an alignment is recombination free and infer the optimal alignment-wide recombination breakpoints, it does not directly pinpoint which sequences are the potential recombinants. On the other hand, `3SEQ` examines all possible sequence triplets to identify potential recombinants and the corresponding breakpoints. We manually parallelized the `3SEQ` analyses so that the sequences could be evaluated concurrently, demonstrating the practicality of such a test for serving as a quick check prior to a phylodynamic analysis.

##### S2.2 Data and Code Availability

The sequence, sampling time and location data used in this study are maintained in the GitHub repository (https://github.com/jsigao/ssstree_supparchive) and archived in the Zenodo repository (https://doi.org/10.5281/zenodo.15574519). Our repositories also contain BEAST XML scripts used to perform the phylodynamic analyses and R scripts used to post process the analysis outputs and compute MCMC diagnostics. We provide major posterior outputs (with some [such as the posterior trees] excluded or subsampled due to the size limit) of the BEAST analyses in the GitHub repository, while more posterior output files that are more densely sampled are available in the Zenodo repository.

#### S2.3 Expanded Meta Summaries of Empirical Analyses

In this section, we provide additional summaries across all datasets, complementing the results presented in the main text and in Section S1.

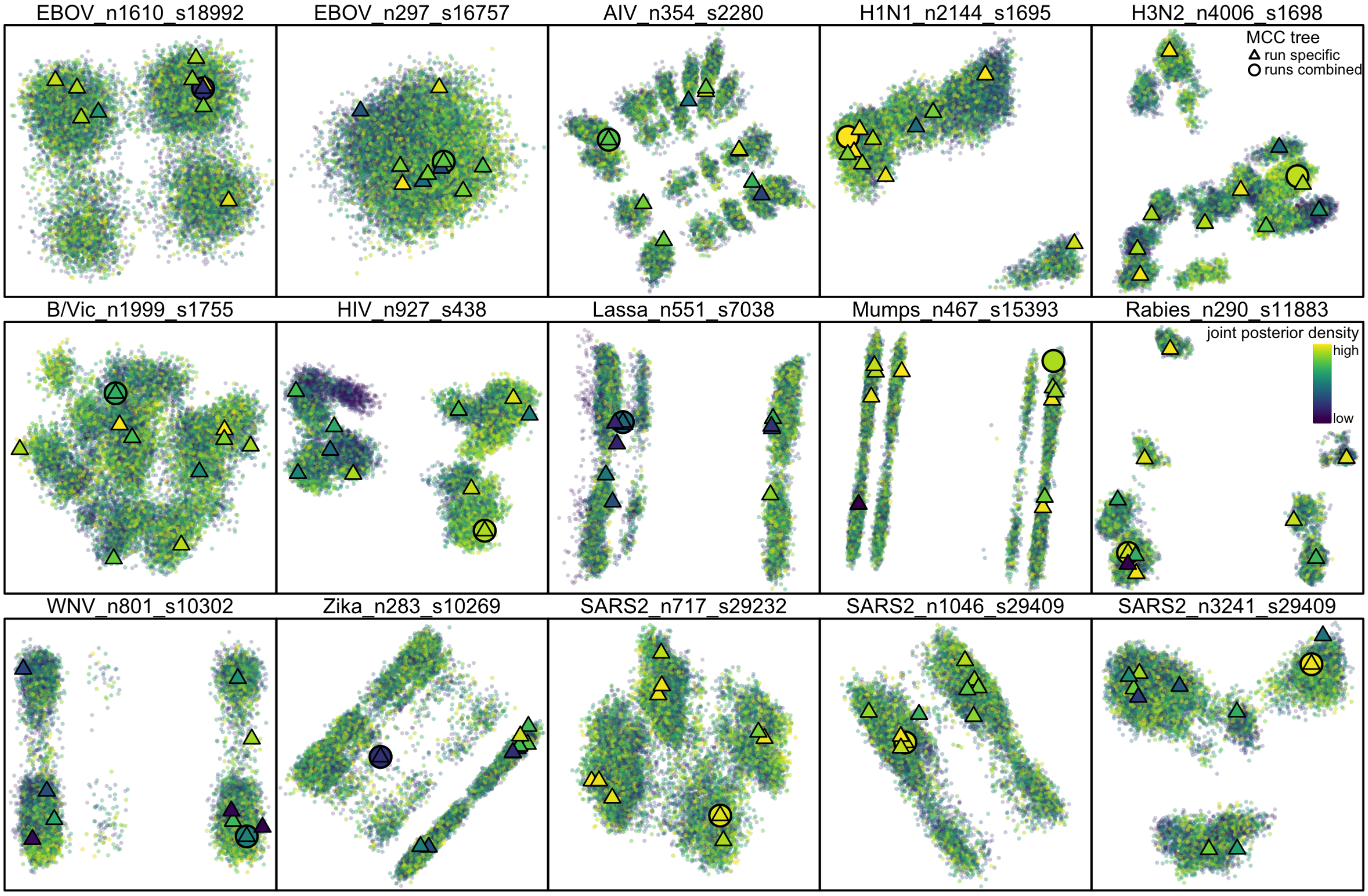


**Figure S40: Phylodynamic posterior tree space is diffuse, comprising many local peaks.** Each panel shows the MDS plot visualizing the posterior tree space of a dataset. A triangle represents the MCC tree summarized from an MCMC chain, while an open circle indicates the MCC tree summarized across all the replicates of that dataset. This figure is identical to Fig. 3 other than that each sampled tree here is colored by the corresponding joint posterior density value (yellow: high; purple: low). The color gradient does not exhibit spatial segregation for almost all the datasets, implying that the posterior tree space is diffuse but contains numerous local peaks with virtually the same height.

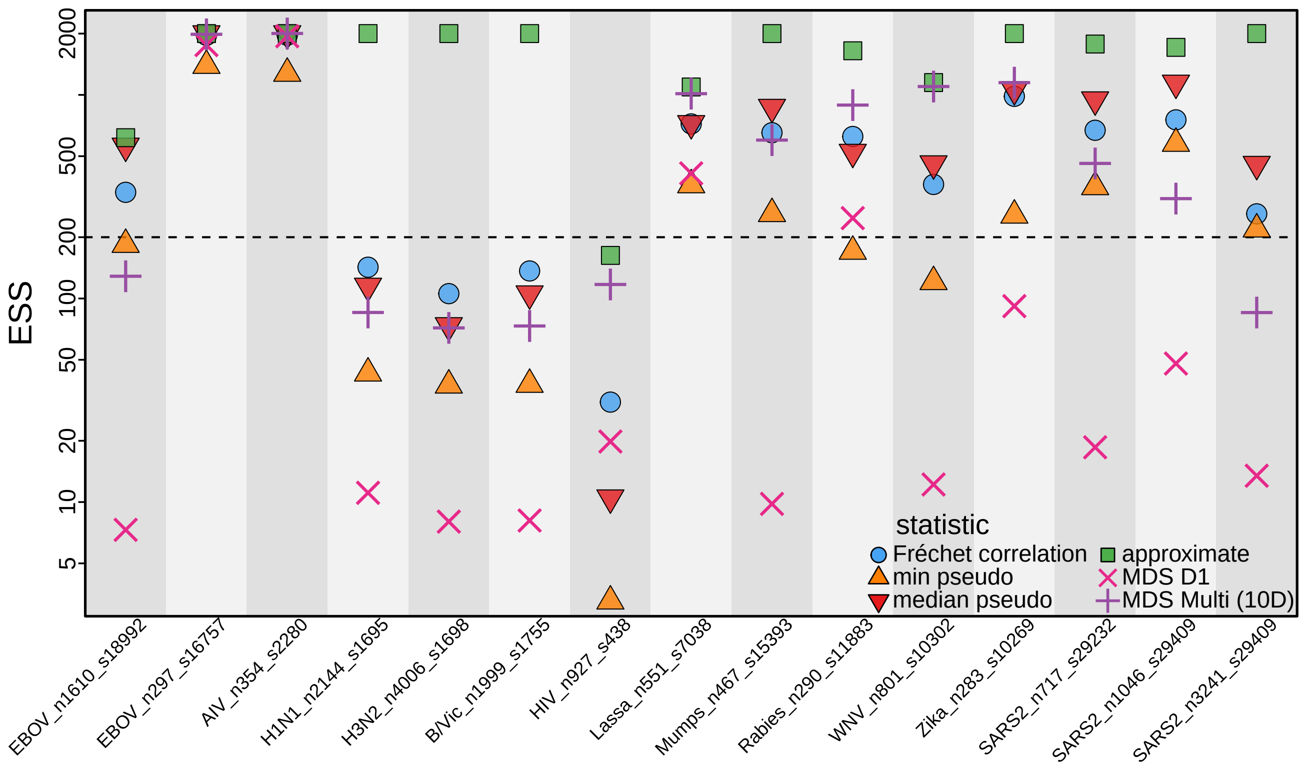


**Figure S41: Tree topology ESS diagnostics assess tree mixing performance within each MCMC chain.** Each column, indicated by the alternating gray/white background, corresponds to a dataset. We present various topology ESS diagnostics (computed using the pairwise RF distances), represented by the colored dots in each column. These diagnostics include the Fréchet correlation ESS (3), the minimum and median pseudo tree ESS, as well as the approximate tree ESS (42). We also include the MDS ESS suggested by Magee et al. (2024); specifically, we compute the MDS D1 ESS by treating the first dimension coordinate of the two-dimension scaling as a continuous variable, and the MDS Multi ESS by treating the MDS coordinates obtained under a higher number of dimensions (10 used here) as a multivariate variable. These ESS diagnostics are largely consistent among themselves (with the Fréchet correlation ESS included as the topology ESS presented in Fig. S5), except that the approximate tree ESS appears to be too liberal about the number of effective independent samples, likely a result of the diffuse tree space.

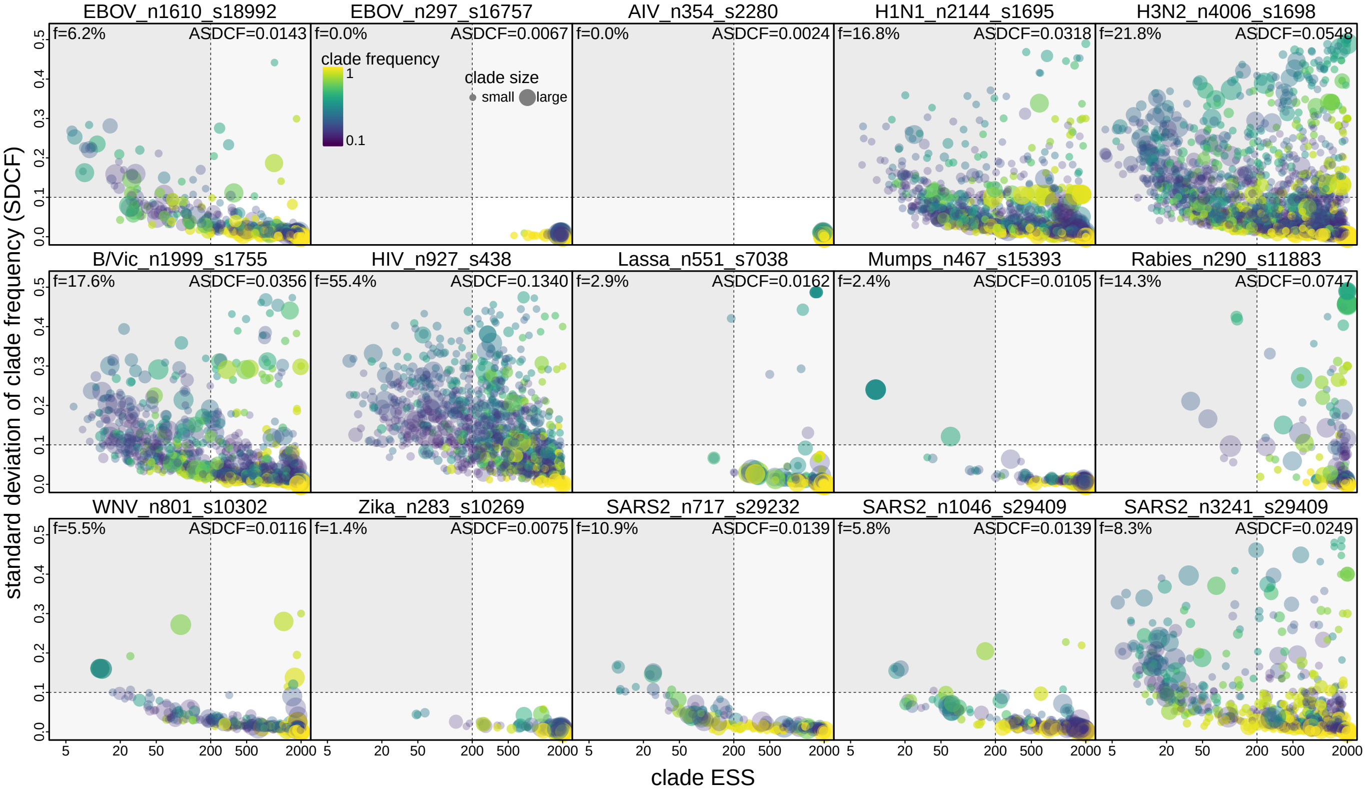


**Figure S42: For most datasets, a small fraction of the clades exhibit sampling problems.** Standard deviation of clade frequency (SDCF; $y$-axis) quantifies the between-chain convergence and clade ESS ($x$-axis) quantifies the within-chain mixing performance, respectively. Each panel corresponds to a dataset. Each dot represents a clade, whose color indicates the clade frequency (*i.e.*, posterior probability) averaged across all replicates (yellow: high; purple: low) and size is proportional to the number of descendant tips. Dashed lines indicate the thresholds typically used in phylogenetic studies; *i.e.*, SDCF above 0.1 (horizontal dashed line) indicates lack of convergence and ESS below 200 (vertical dashed line) indicates inadequate mixing. For most datasets, a small fraction ($f$) of the dots reside outside the bottom right quadrant surrounded by the dashed lines, indicating that the frequency of those clades failed to be estimated reliably.

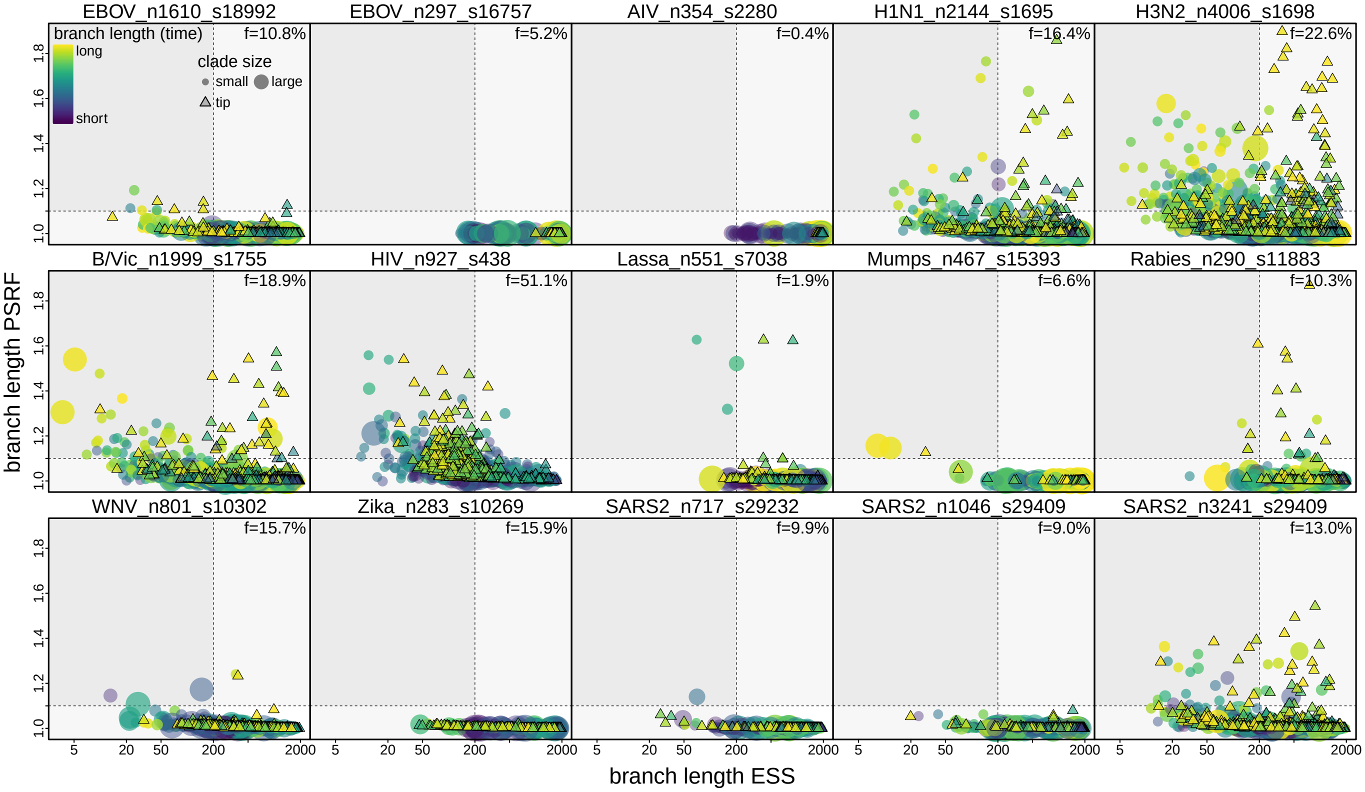


**Figure S43: For most datasets, a small fraction of the branch-length estimates exhibit sampling problems.** Each panel shows the MCMC diagnostics computed for the length (in unit of time) of each tree branch (with those whose subtended clade has a posterior probability smaller than 0.1 in more than five out of the ten replicate chains excluded) in a dataset. Potential scale reduction factor (PSRF; $y$-axis) quantifies convergence between the chains, and effective sample size (ESS; $x$-axis) quantifies mixing within each chain. Dashed lines indicate the thresholds typically used in phylogenetic studies; *i.e.*, PSRF above 1.1 (horizontal dashed line) indicates lack of convergence and ESS below 200 (vertical dashed line) indicates inadequate mixing. Circles and triangles represent internal and external branches, respectively, whose color indicates the inferred branch length value (yellow: high; purple: low) and size is proportional to the number of descendant tips. For most datasets, a small fraction ($f$) of the dots reside outside the bottom right quadrant surrounded by the dashed lines, indicating that those branch lengths failed to be estimated reliably.

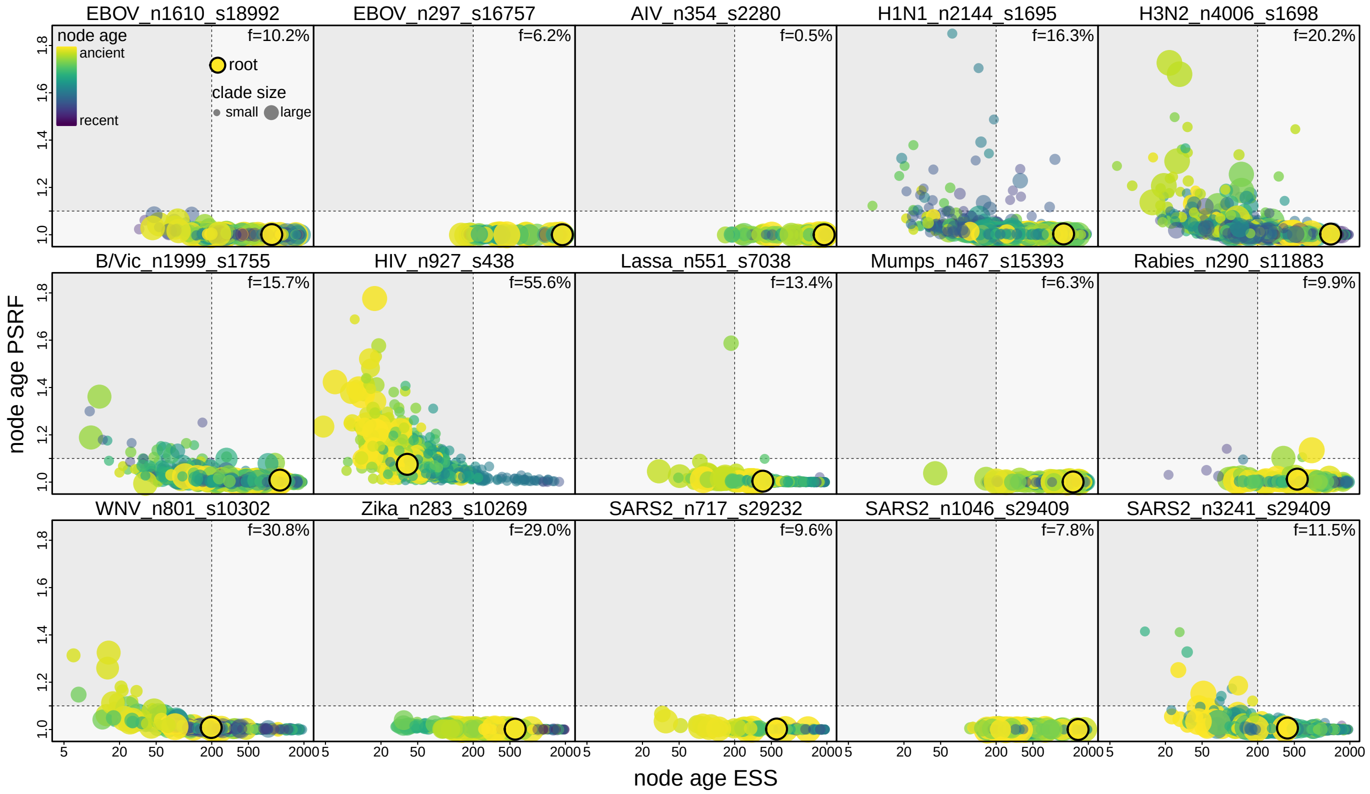


**Figure S44: For most datasets, a small fraction of the node-age estimates exhibit sampling problems.** Each panel shows the MCMC diagnostics computed for the age estimate of each internal node (*i.e.*, clade tMRCA; with those whose posterior probability below 0.1 in more than five out of the ten replicate chains excluded) in a dataset. PSRF ($y$-axis) quantifies convergence between the chains, and ESS ($x$-axis) quantifies mixing within each chain. Dashed lines indicate the thresholds typically used in phylogenetic studies; *i.e.*, PSRF above 1.1 (horizontal dashed line) indicates lack of convergence and ESS below 200 (vertical dashed line) indicates inadequate mixing. Each dot represents an internal node, whose color indicates the inferred age value (yellow: ancient; purple: recent) and size is proportional to the number of descendant tips; the root node is marked by a closed circle. For most datasets, a small fraction ($f$) of the dots reside outside the bottom right quadrant surrounded by the dashed lines, indicating that those node ages failed to be estimated reliably.

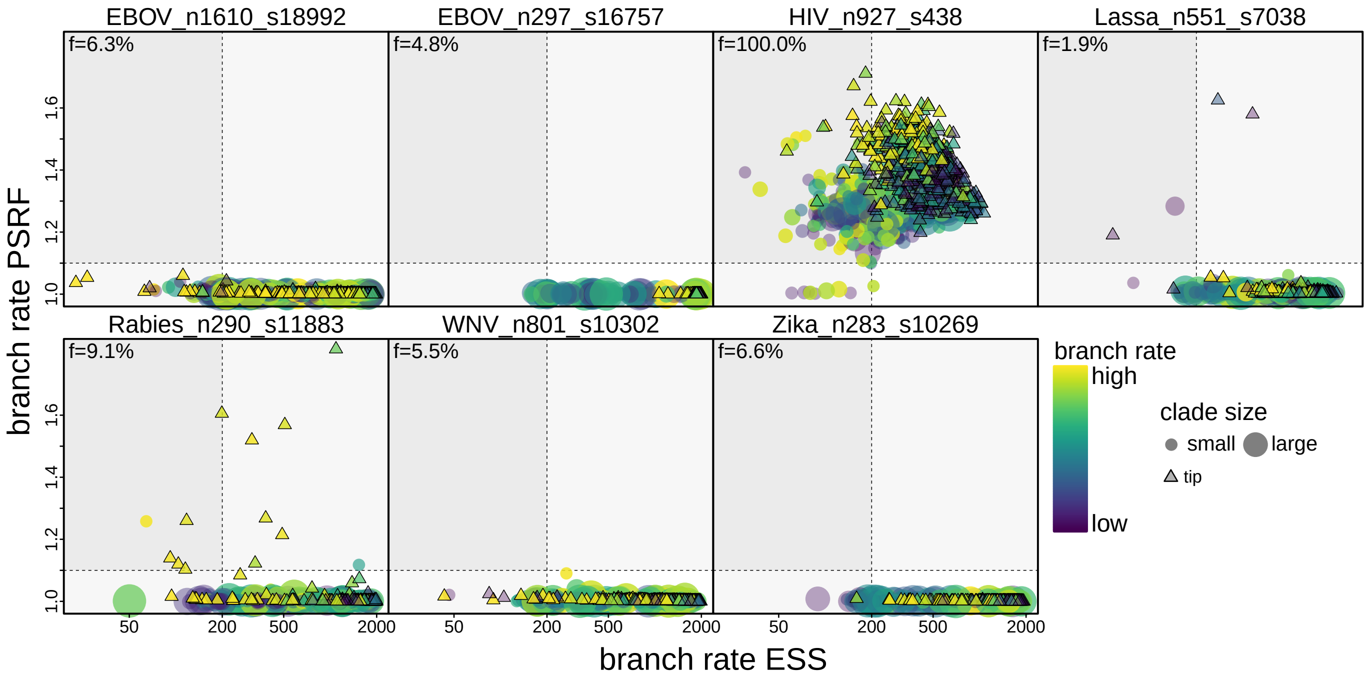


**Figure S45: For most datasets, a small fraction of the branch-rate estimates exhibit sampling problems.** Each panel shows the MCMC diagnostics computed for the evolutionary rate on each tree branch (with those whose subtended clade has a posterior probability smaller than 0.1 in more than five out of the ten replicate chains excluded) in a dataset. Here only the seven datasets whose analyses assume a relaxed-clock model are included. PSRF ($y$-axis) quantifies convergence between the chains, and ESS ($x$-axis) quantifies mixing within each chain. Dashed lines indicate the thresholds typically used in phylogenetic studies; *i.e.*, PSRF above 1.1 (horizontal dashed line) indicates lack of convergence and ESS below 200 (vertical dashed line) indicates inadequate mixing. Circles and triangles represent internal and external branches, respectively, whose color indicates the inferred branch rate value (yellow: high; purple: low) and size is proportional to the number of descendant tips. For most datasets, a small fraction ($f$) of the dots reside outside the bottom right quadrant surrounded by the dashed lines, indicating that those branch rates failed to be estimated reliably. The HIV dataset is an exception as almost no branch rate appears to be estimated reliably for that dataset, which is presumably due to the lack of convergence for the substitution model parameter estimates.

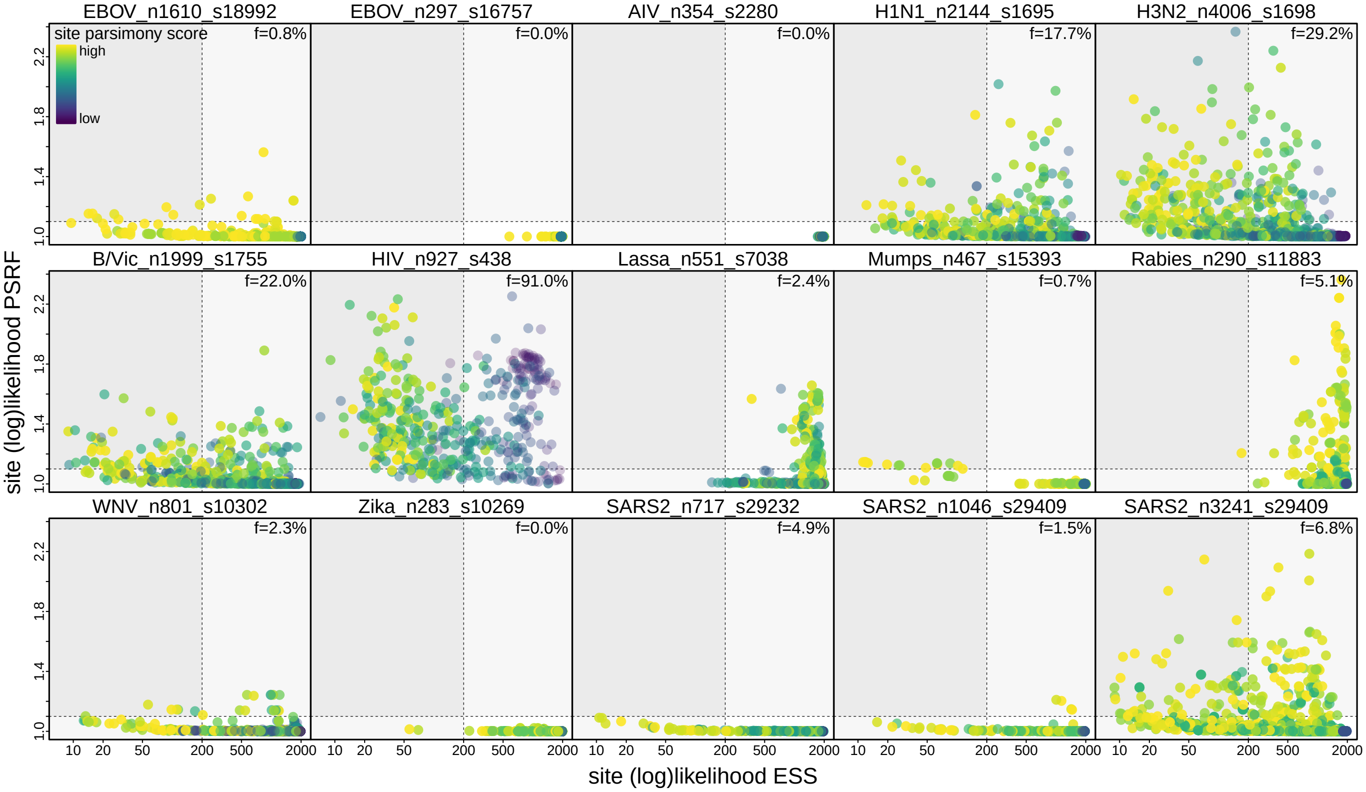


**Figure S46: For most datasets, a small fraction of the sites in the sequence alignment are associated with sampling problems.** Each panel shows the MCMC diagnostics computed for the likelihood of each site in a dataset. Each dot represents a site, whose color (yellow: high; purple: low) indicates the parsimony number of mutations at that site averaged over the posterior trees. PSRF ($y$-axis) quantifies convergence between the chains, and ESS ($x$-axis) quantifies mixing within each chain. Dashed lines indicate the thresholds typically used in phylogenetic studies; *i.e.*, PSRF above 1.1 (horizontal dashed line) indicates lack of convergence and ESS below 200 (vertical dashed line) indicates inadequate mixing. For most datasets, a small fraction ($f$) of the dots reside outside the bottom right quadrant surrounded by the dashed lines, indicating that those sites may have increased the difficulty of MCMC sampling.

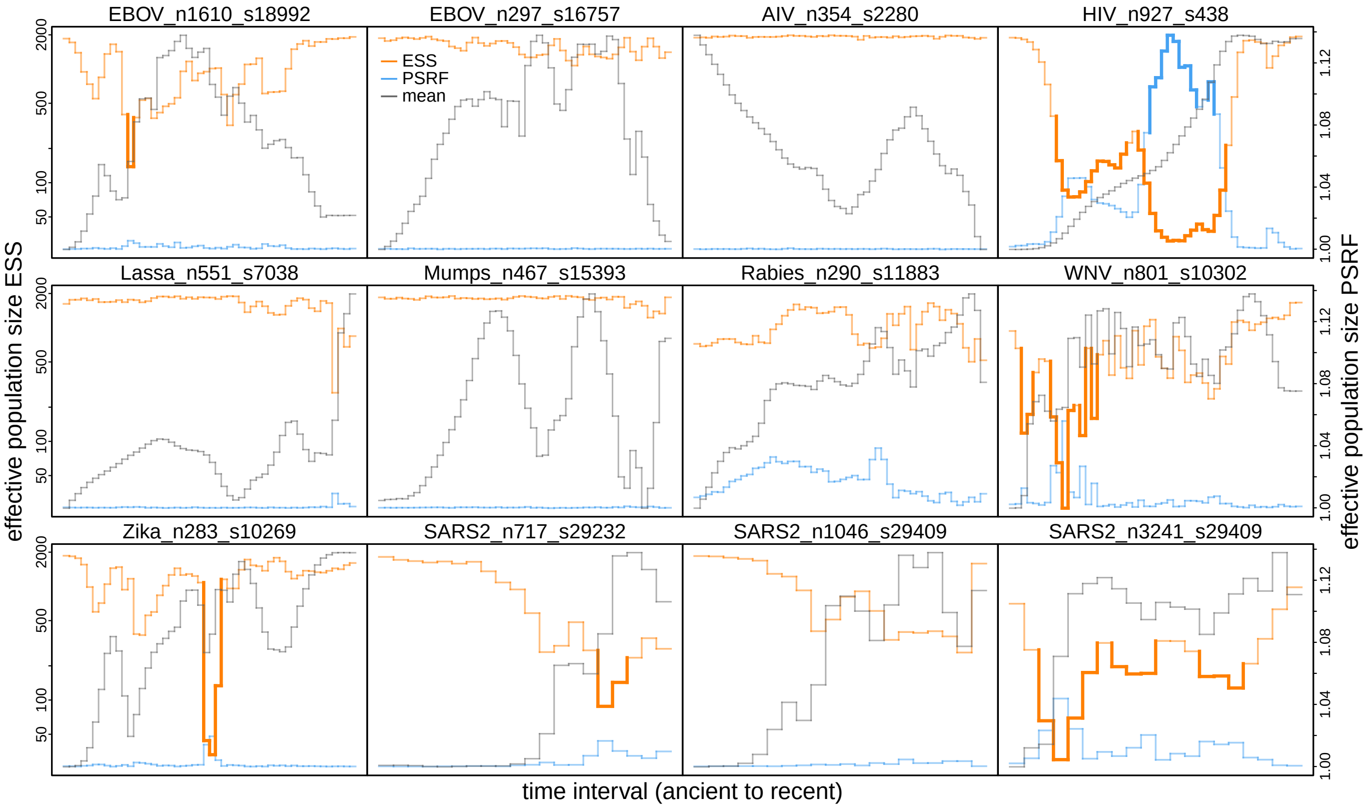


**Figure S47: Interval-specific diagnostics reveal that impact of tree sampling problems appears to be minor on effective population size estimates.** Each panel depicts MCMC diagnostics (ESS: left axis, orange; PSRF: right axis, blue) computed using the effective population size estimates of each time interval of a dataset (the three human influenza virus datasets that assumed a constant coalescent model in their analyses are excluded here). The segments are bolded with a darker color in certain intervals to indicate that the corresponding ESS is below the threshold (200) or PSRF is above the threshold (1.1) for that interval. It appears that most of the intervals with unsatisfactory MCMC performance coincide with rapid changes in the effective population size (gray) between neighboring intervals, consistent with the pattern shown in Fig. S26 that the inferred demographic temporal trajectories are shifted to be a slightly younger or older.

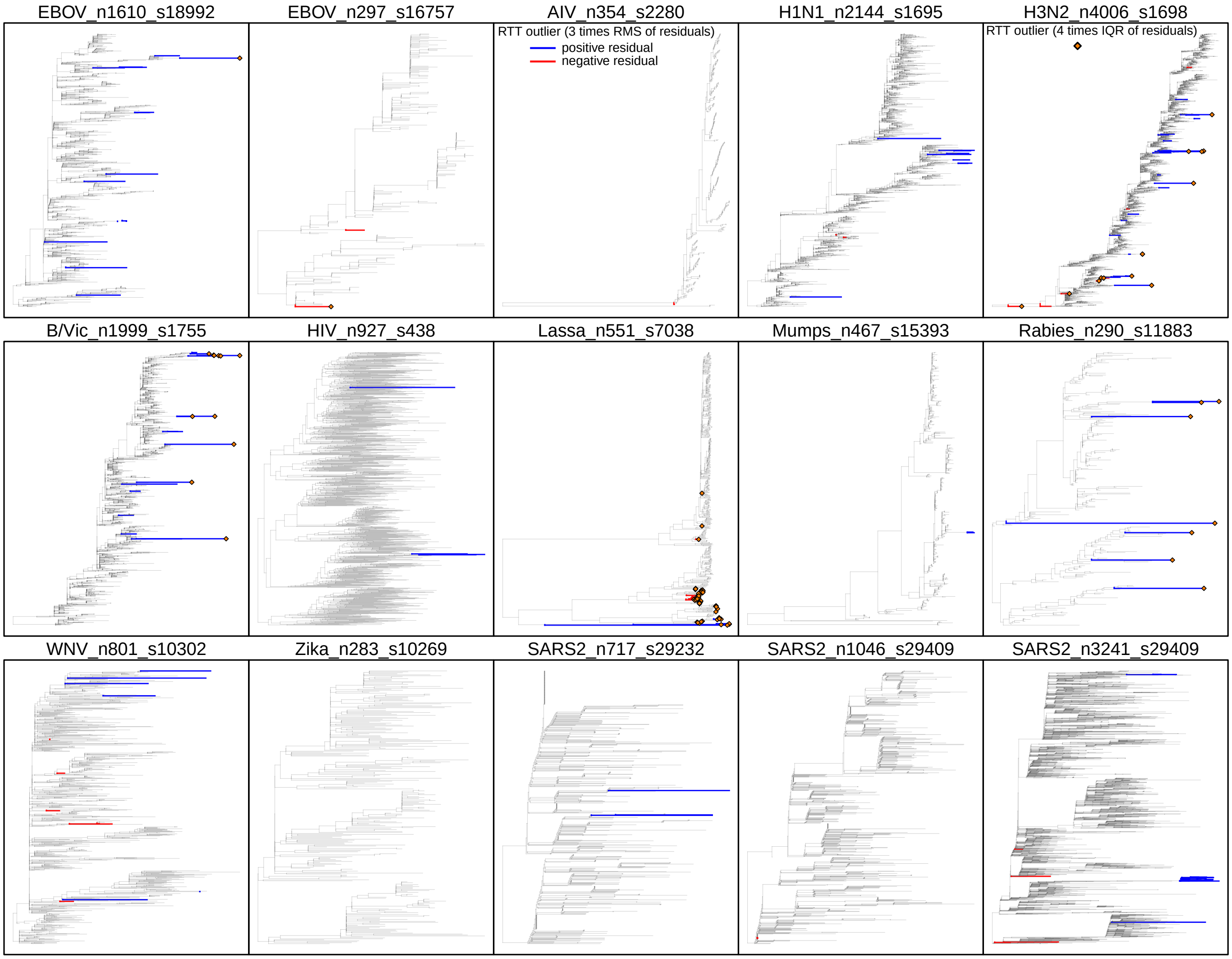


**Figure S48: Rooted ML tree and the identified RTT outliers.** Each panel presents the maximum-likelihood (ML) tree inferred with each empirical dataset. For each dataset, we first infer an unrooted ML phylogram using IQ-TREE and then root it using the RTT regression test to obtain the ML tree depicted here. The tips that fail the RTT regression test (*i.e.,* identified as "RTT outliers") are marked on each tree; blue and red external branches indicate the corresponding tip's residual is greater than three times the root mean square of the residuals (RMS) or smaller than three times the negative RMS, respectively, while orange diamonds indicate the tips whose absolute residual is greater than four times the interquartile range (IQR) of the residuals. The tips marked in the tree here correspond to the colored dots in the regression results shown in Fig. S25.

##### S2.4 Expanded Dataset-Specific Summaries of Empirical Analyses

In this section, we provide dataset-specific descriptions of phylodynamic analysis setup and result summaries that have been covered in the main text or sections above. These results reveal a highly consistent pattern across all datasets, further demonstrating the widespread tree sampling problems and their impacts on phylodynamic inferences.

###### S2.4.1 Ebola Virus West African Outbreak Dataset

Dudas et al. (10) investigated the dispersal and proliferation of Ebola virus during the 2013–2016 West African outbreak by analyzing viral genomes collected from over 5% of the known cases in that outbreak. This study contains a single sequence dataset comprised of 1610 Ebola virus genomes with 18992 nucleotide sites. One of the phylogenetic inferences of the original study was later shown to exhibit multimodality, with the different replicate analyses converging to different regions in tree space (50).

We directly acquired the BEAST XML scripts provided by the original study from its supplementary repository—containing both the sequence alignment and sampling time and location data—and specify our analyses accordingly. Specifically, following the original study, we divide the genomic alignment into four partitions, including three partitions that each corresponds to a codon position for the coding region and another partition for the non-coding intergenic region. For each partition, we specify an independent HKY+$\Gamma_4$ substitution model (51, 52) and a partition-specific rate multiplier whose mean is constrained to be one across the partitions. The phylodynamic model also includes an uncorrelated lognormal branch-rate prior model (53, 54) and a Skygrid coalescent node-age model (55).

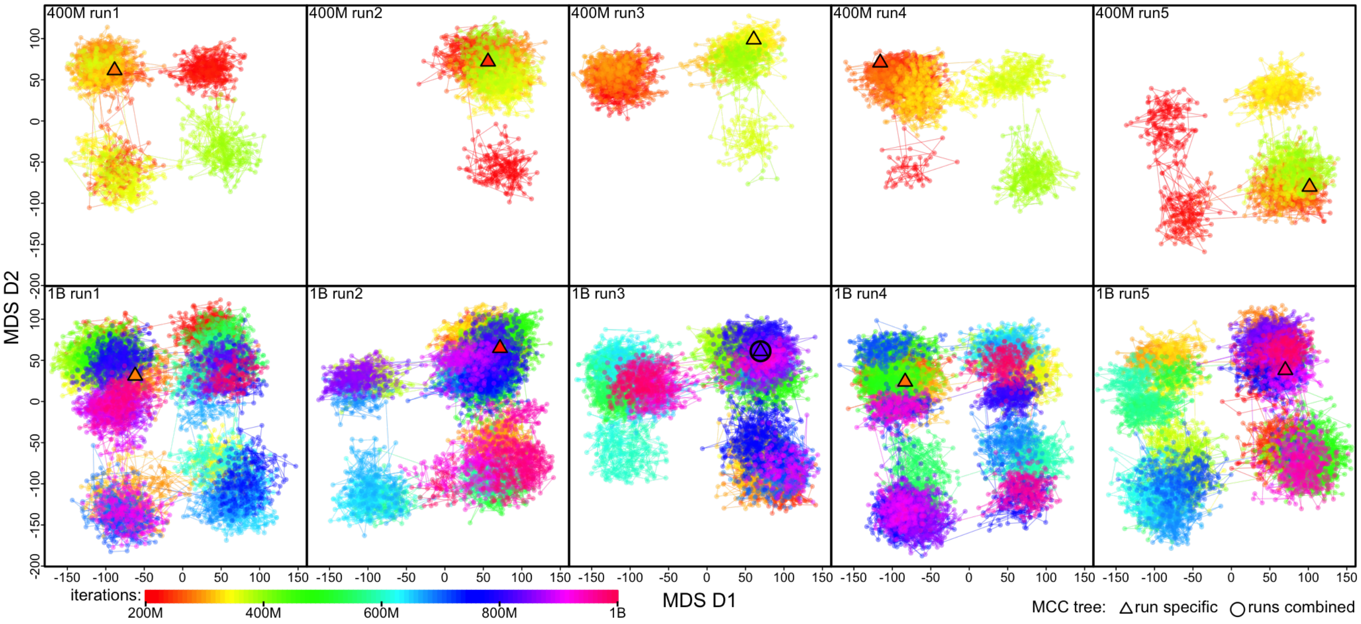


**Figure S49: MCMC sampler progression in tree space of the EBOV_n1610_s18992 dataset.** Each panel shows the MDS plot visualizing the posterior tree space inferred with one of the ten MCMC replicate chains. The MDS coordinates are calculated based on the pairwise RF distances computed over all sampled posterior trees. Each dot represents a tree sampled in the posterior whose rainbow color indicates the corresponding iteration index (see inset color gradient bar). The top and bottom rows present the five 400-million-iteration chains ("400M run") and the five 1-billion-iteration chains ("1B run"), respectively. A triangle represents the MCC tree summarized from an MCMC chain, while an open circle indicates the MCC tree summarized across all the replicates of that dataset. Dots are connected with lines to indicate that they are sampled consecutively in the chain.

##### S2.4.2 Ebola Virus DRC relapse Dataset

Mbala-Kingebeni et al. (20) investigated an Ebola virus transmission chain that is linked to a patient who was initially infected by Ebola virus during the 2018–2020 disease outbreak in North Kivu province in the Democratic Republic of Congo and then experienced a relapse half a year later. This study contains a single sequence dataset comprised of 297 Ebola virus genomes with 16757 nucleotide sites, collected from that focal patient, the initial outbreak, the later transmission chain, and other epidemiologically linked cases. We directly acquired the BEAST XML scripts provided by the original study from its supplementary repository—containing both the sequence alignment and sampling times—and specify our analyses accordingly.

Specifically, following the original study, we divide the genomic alignment into two partitions, where the first partition includes sites at the first and second codon positions, and the second partition includes sites at the third codon position. For each partition, we specify an independent HKY+$\Gamma_4$ substitution model (51, 52) and a partition-specific rate multiplier whose mean is constrained to be one across the partitions. The phylodynamic model also includes an uncorrelated lognormal branch-rate prior model (53, 54) and a Skygrid coalescent node-age model (55).

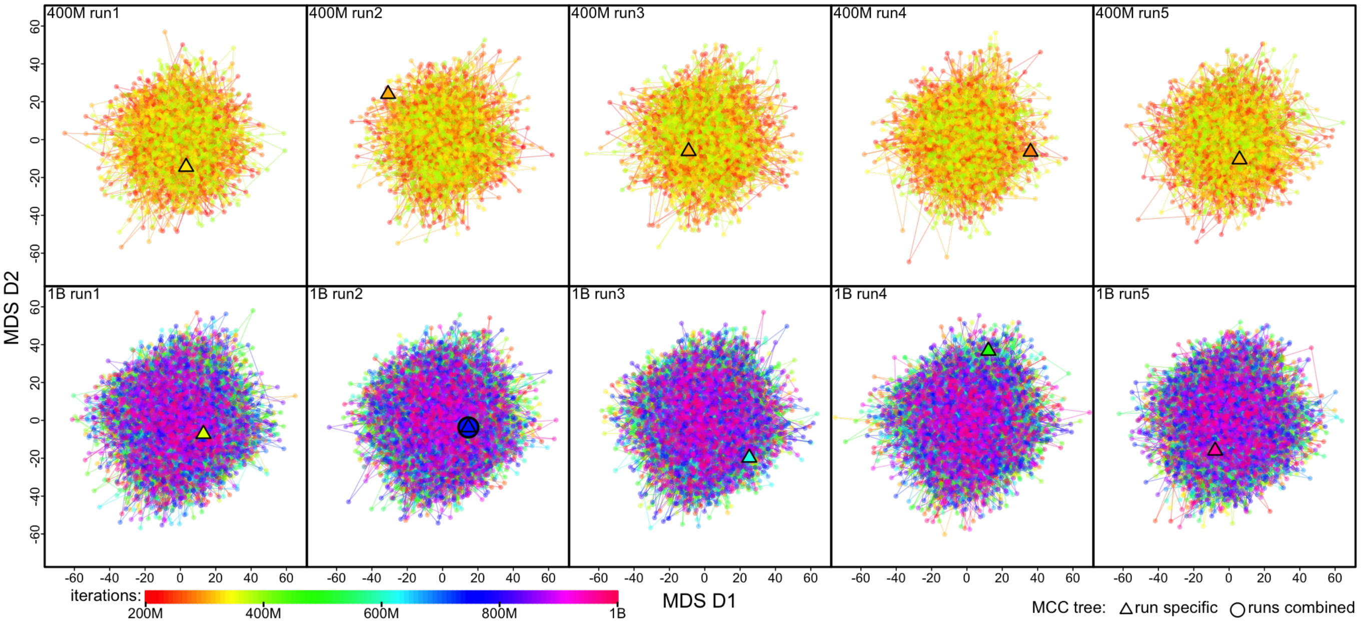


**Figure S50: MCMC sampler progression in tree space of the EBOV_n297_s16757 dataset.** Each panel shows the MDS plot visualizing the posterior tree space inferred with one of the ten MCMC replicate chains. The MDS coordinates are calculated based on the pairwise RF distances computed over all sampled posterior trees. Each dot represents a tree sampled in the posterior whose rainbow color indicates the corresponding iteration index (see inset color gradient bar). The top and bottom rows present the five 400-million-iteration chains ("400M run") and the five 1-billion-iteration chains ("1B run"), respectively. A triangle represents the MCC tree summarized from an MCMC chain, while an open circle indicates the MCC tree summarized across all the replicates of that dataset. Dots are connected with lines to indicate that they are sampled consecutively in the chain.

##### S2.4.3 Human Influenza Virus Datasets

Bedford et al. (11) inferred the geographic dynamics of the four most prevalent, globally circulating human seasonal influenza viruses (A/H3N2, A/H1N1, B/Victoria, and B/Yamagata). They collected complete sequences of the HA1 domain of the hemagglutinin (HA) gene for these influenza viruses, sampled across most of the major geographic areas between 2000–2012. We included three sequence datasets from that study, including (1) 2144 H1N1 sequences with 1695 nucleotide sites, (2) 4006 H3N2 sequences with 1698 nucleotide sites, and (3) 1999 B/Victoria sequences with 1755 nucleotide sites. We directly acquired the `BEAST` XML scripts provided by the original study from its supplementary repository—containing both the sequence alignment and sampling time and location data—and specify our analyses accordingly.

Specifically, for each of the three datasets, following the original study, we divide the genomic alignment into two partitions, where the first partition includes sites at the first and second codon positions, and the second partition includes sites at the third codon position. For each partition, we specify an independent HKY+$\Gamma_4$ substitution model (51, 52) and a partition-specific rate multiplier whose mean is constrained to be one across the partitions. The phylodynamic model also includes a strict molecular clock model that assumes a single mutation rate across all branches and a constant coalescent node-age model (56).

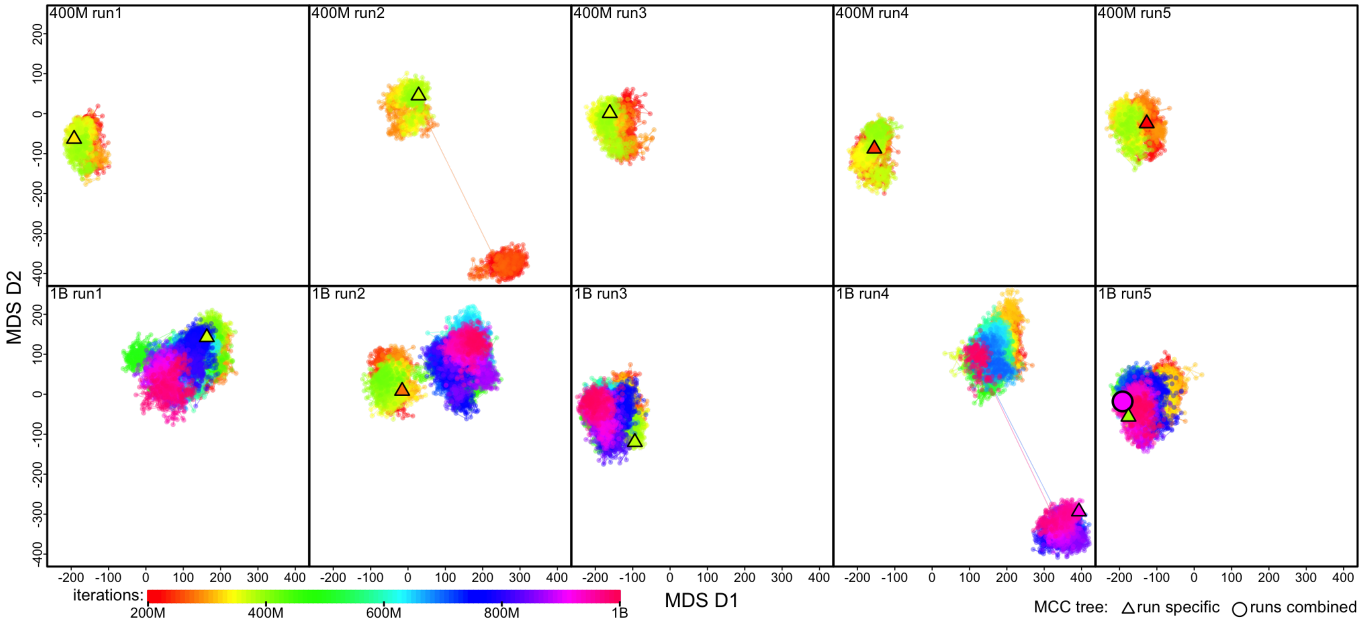


**Figure S51: MCMC sampler progression in tree space of the H1N1_n2144_s1695 dataset.** Each panel shows the MDS plot visualizing the posterior tree space inferred with one of the ten MCMC replicate chains. The MDS coordinates are calculated based on the pairwise RF distances computed over all sampled posterior trees. Each dot represents a tree sampled in the posterior whose rainbow color indicates the corresponding iteration index (see inset color gradient bar). The top and bottom rows present the five 400-million-iteration chains ("400M run") and the five 1-billion-iteration chains ("1B run"), respectively. A triangle represents the MCC tree summarized from an MCMC chain, while an open circle indicates the MCC tree summarized across all the replicates of that dataset. Dots are connected with lines to indicate that they are sampled consecutively in the chain.

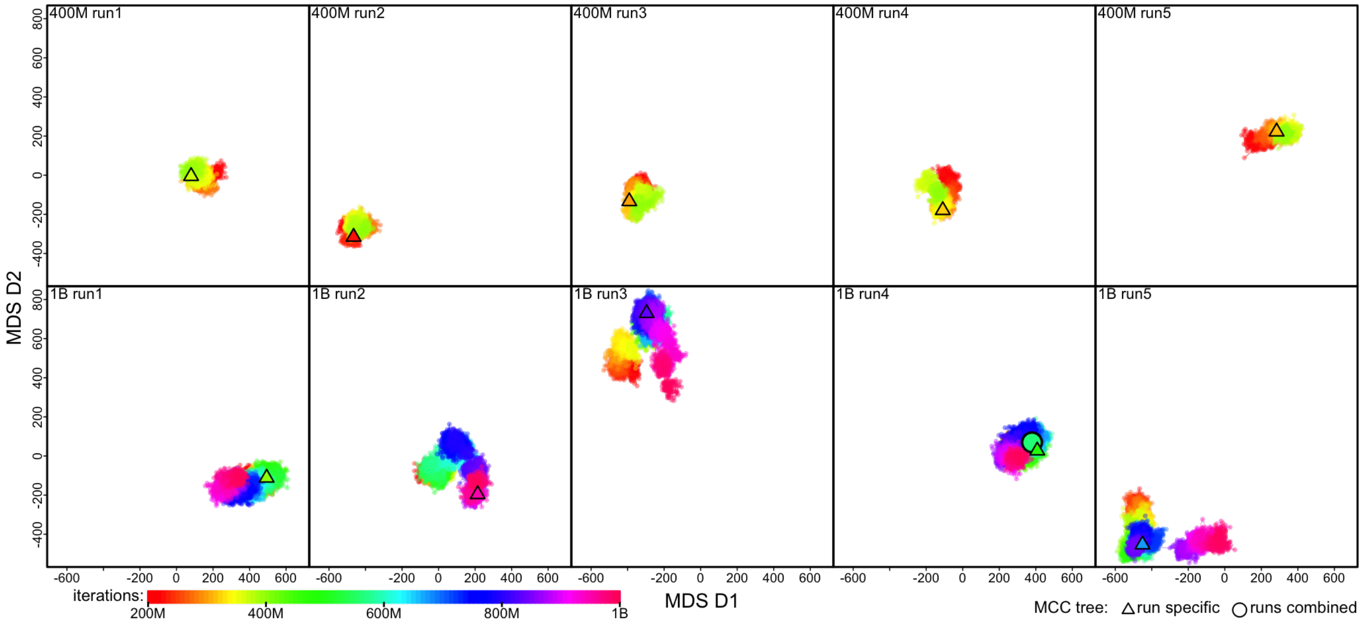


**Figure S52: MCMC sampler progression in tree space of the H3N2_n4006_s1698 dataset.** Each panel shows the MDS plot visualizing the posterior tree space inferred with one of the ten MCMC replicate chains. The MDS coordinates are calculated based on the pairwise RF distances computed over all sampled posterior trees. Each dot represents a tree sampled in the posterior whose rainbow color indicates the corresponding iteration index (see inset color gradient bar). The top and bottom rows present the five 400-million-iteration chains ("400M run") and the five 1-billion-iteration chains ("1B run"), respectively. A triangle represents the MCC tree summarized from an MCMC chain, while an open circle indicates the MCC tree summarized across all the replicates of that dataset. Dots are connected with lines to indicate that they are sampled consecutively in the chain.

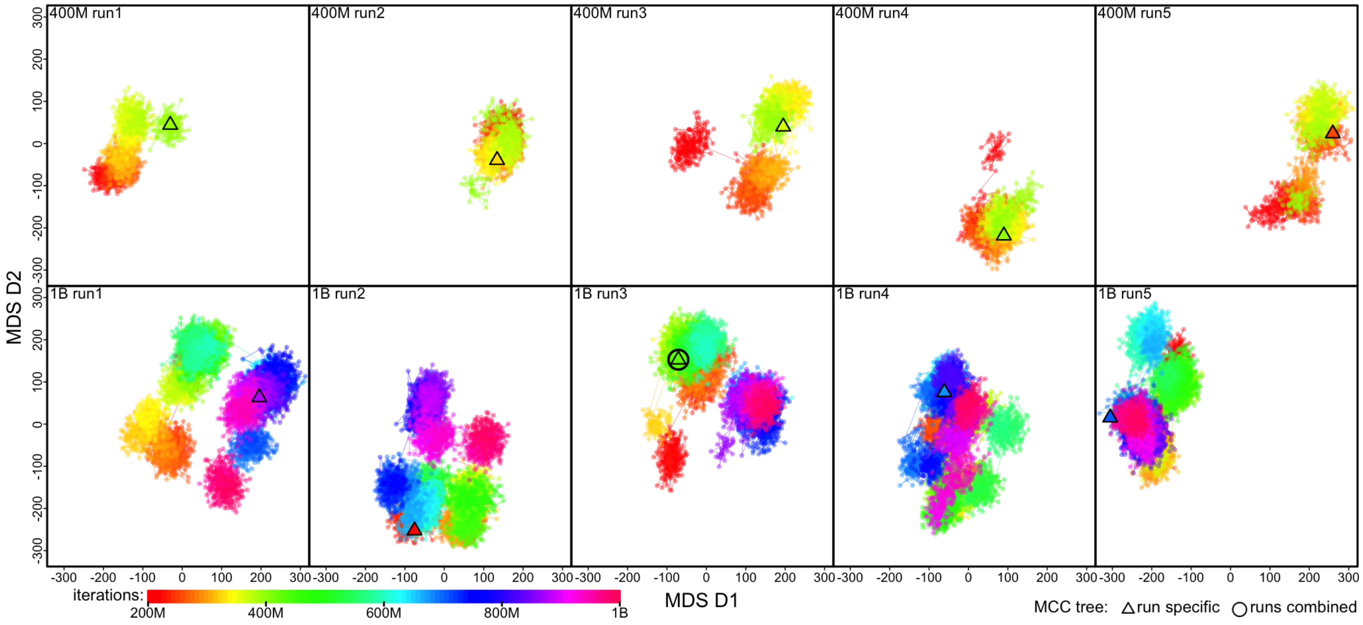


**Figure S53: MCMC sampler progression in tree space of the B/Vic_n1999_s1755 dataset.** Each panel shows the MDS plot visualizing the posterior tree space inferred with one of the ten MCMC replicate chains. The MDS coordinates are calculated based on the pairwise RF distances computed over all sampled posterior trees. Each dot represents a tree sampled in the posterior whose rainbow color indicates the corresponding iteration index (see inset color gradient bar). The top and bottom rows present the five 400-million-iteration chains ("400M run") and the five 1-billion-iteration chains ("1B run"), respectively. A triangle represents the MCC tree summarized from an MCMC chain, while an open circle indicates the MCC tree summarized across all the replicates of that dataset. Dots are connected with lines to indicate that they are sampled consecutively in the chain.

##### S2.4.4 Avian Influenza Virus Dataset

Worobey et al. (21) studied the molecular evolution pattern across various influenza A viruses and identified a global selective sweep in their recent evolutionary history. They demonstrate this synchronized global sweep by analyzing each influenza gene segment individually and revealing the consistent host-specific substitution rates across all the segments. We included one of the gene segments, PB2, as it has the most complete sequence data among the eight segments; the PB2 sequence dataset comprises 354 sequences with 2280 nucleotide sites. We directly acquired the `BEAST` XML scripts provided by the original study from its supplementary repository—containing both the sequence alignment and sampling times—and specify our analyses accordingly.

Specifically, following the original study, we divide the genomic alignment into two partitions, where the first partition includes sites at the first and second codon positions, and the second partition includes sites at the third codon position. For each partition, we specify an independent HKY+$\Gamma_4$ substitution model (51, 52) and a partition-specific rate multiplier whose mean is constrained to be one across the partitions. To capture the host-specific molecular evolution dynamics, a local clock model (57) is specified for each of the nine major clades in the tree defined by the host species. The phylodynamic model also includes a Skygrid coalescent node-age model component (55).

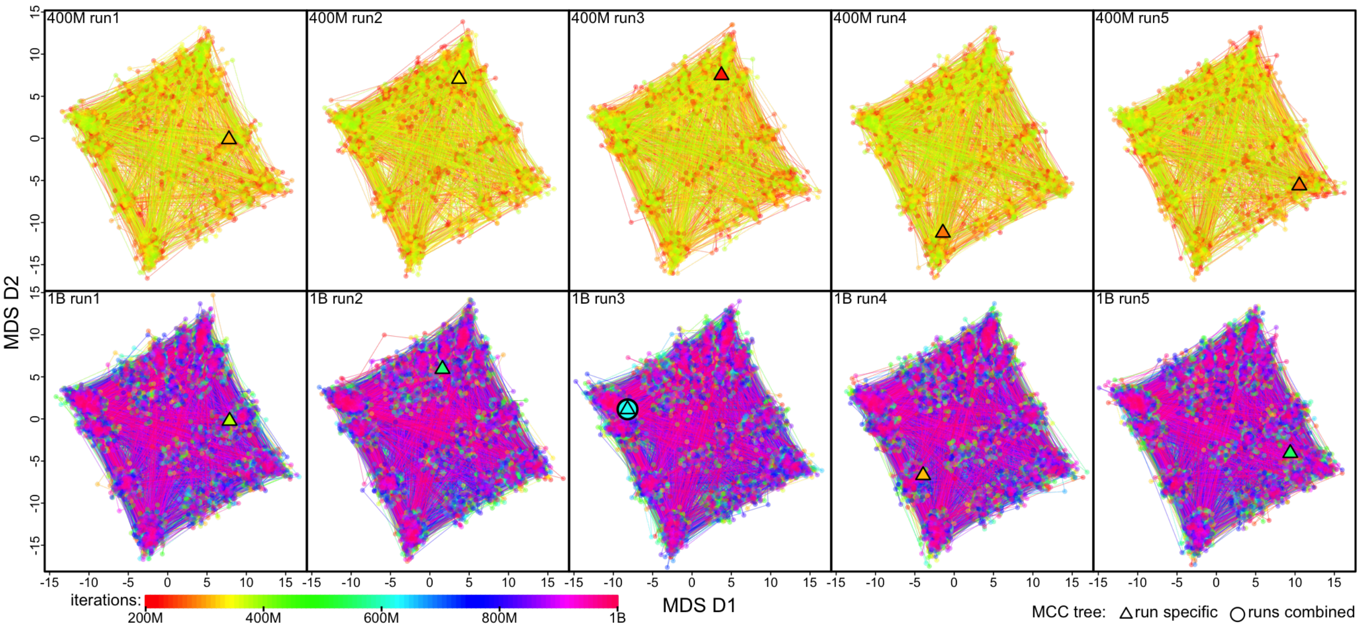


**Figure S54: MCMC sampler progression in tree space of the AIV_n354_s2280 dataset.** Each panel shows the MDS plot visualizing the posterior tree space inferred with one of the ten MCMC replicate chains. The MDS coordinates are calculated based on the pairwise RF distances computed over all sampled posterior trees. Each dot represents a tree sampled in the posterior whose rainbow color indicates the corresponding iteration index (see inset color gradient bar). The top and bottom rows present the five 400-million-iteration chains ("400M run") and the five 1-billion-iteration chains ("1B run"), respectively. A triangle represents the MCC tree summarized from an MCMC chain, while an open circle indicates the MCC tree summarized across all the replicates of that dataset. Dots are connected with lines to indicate that they are sampled consecutively in the chain.

##### S2.4.5 HIV Dataset

Faria et al. (14) explored the origin and early spread of HIV-1 in human populations by analyzing sequences collected from central and southeast Africa and America between the 1980s and early 2000s. We included the Dataset B from that study, containing 927 HIV envelope C2V3 sequences with 438 nucleotide sites, among which 792 sequences were collected between 1985–2004 from eight cities in the Democratic Republic of the Congo and the Republic of the Congo, 67 subtype C sequences were collected from southeast Africa (Zambia, Botswana, Tanzania, Kenya, Uganda, Burundi, Ethiopia and South Africa) sampled between 1986–2005, 67 sequences were collected from the Americas (Haiti, Trinidad and Tobago and the USA) sampled between 1978–1997, and the ZR59 isolate obtained in 1959 from blood collected in Kinshasa.

We directly acquired the BEAST XML scripts provided by the original study from its supplementary repository—containing both the sequence alignment and sampling time and location data—and specify our analyses accordingly. Specifically, following the original study, in our analyses we specify a phylodynamic model with the following components: (1) a GTR+$\Gamma_4$ substitution model (52, 58), (2) an uncorrelated lognormal branch-rate prior model (53, 54), and (3) a Skygrid coalescent node-age model (55).

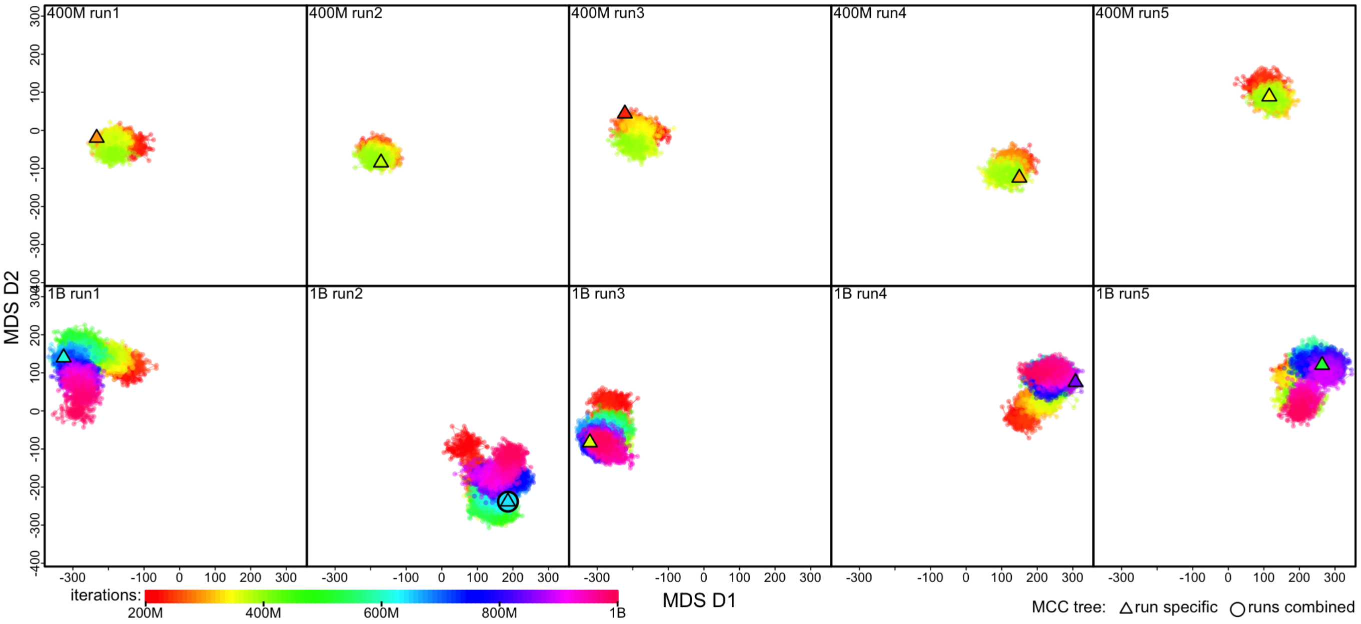


**Figure S55: MCMC sampler progression in tree space of the HIV_n927_s438 dataset.** Each panel shows the MDS plot visualizing the posterior tree space inferred with one of the ten MCMC replicate chains. The MDS coordinates are calculated based on the pairwise RF distances computed over all sampled posterior trees. Each dot represents a tree sampled in the posterior whose rainbow color indicates the corresponding iteration index (see inset color gradient bar). The top and bottom rows present the five 400-million-iteration chains ("400M run") and the five 1-billion-iteration chains ("1B run"), respectively. A triangle represents the MCC tree summarized from an MCMC chain, while an open circle indicates the MCC tree summarized across all the replicates of that dataset. Dots are connected with lines to indicate that they are sampled consecutively in the chain.

##### S2.4.6 Lassa Virus Dataset

Klitting et al. (22) performed a comprehensive phylodynamic and epidemiological study to identify the ecological and environmental factors that are suitable for the introduction and circulation of the Lassa virus (LASV) by analyzing all publicly available LASV sequences. We included the L gene segment dataset from that study, containing 551 LASV sequences with 7038 nucleotide sites.

We directly acquired the `BEAST` XML scripts provided by the original study from its supplementary repository—containing both the sequence alignment and sampling times—and specify our analyses accordingly. Specifically, following the original study, in our analyses we specify a phylodynamic model with the following components: (1) a GTR+$\Gamma_4$ substitution model (52, 58), (2) an uncorrelated lognormal branch-rate prior model (53, 54), and (3) a Skygrid coalescent node-age model (55).

**Table S2: Focal sequences in the Lassa dataset.**

| Short name | Full name | GenBank ID |
|---|---|---|
| 088 | LASV_NGA_2018_IRR_088_NGA-Anambra_Amansea_Hs_2018-02-14 | MK117925 |
| 091 | LASV_NGA_2018_IRR_091_NGA-Edo_Afowa_Hs_2018-02-15 | MK117926 |
| 036 | LASV_NGA_2018_IRR_036_NGA-Edo_Ekpoma_Hs_2018-01-26 | MK117894 |
| 046 | LASV_NGA_2018_IRR_046_NGA-Edo_Ekpoma_Hs_2018-02-05 | MK117898 |

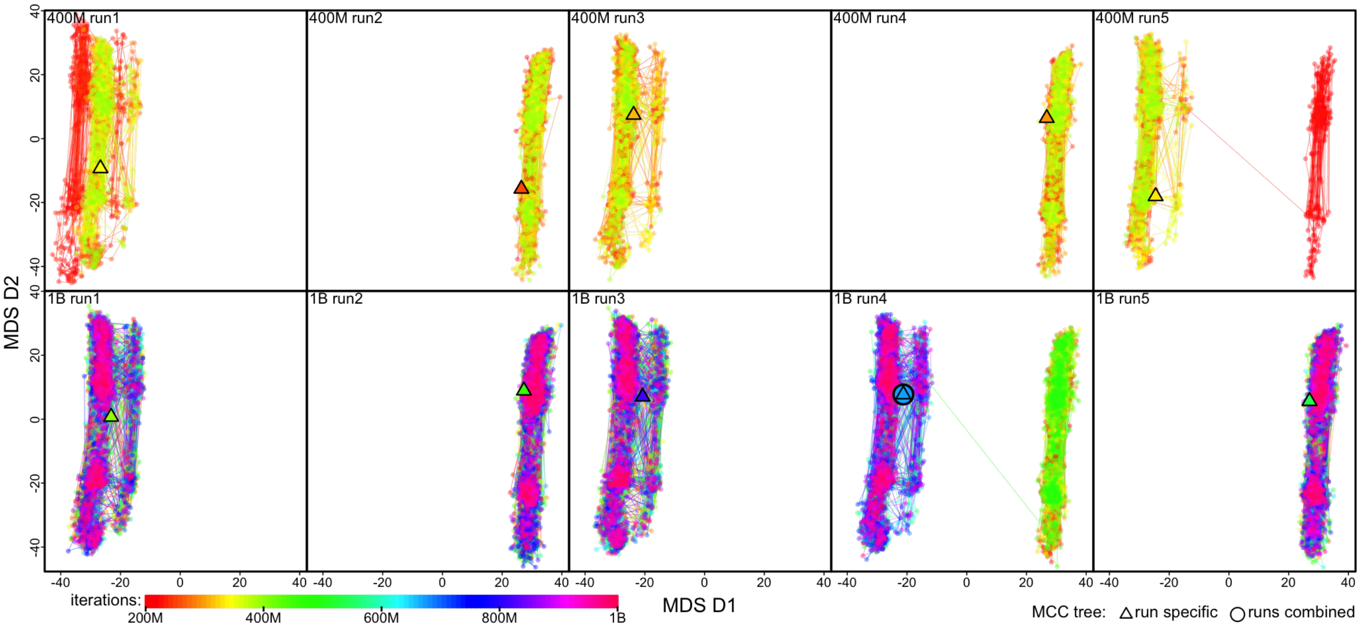


**Figure S56: MCMC sampler progression in tree space of the Lassa_n551_s7038 dataset.** Each panel shows the MDS plot visualizing the posterior tree space inferred with one of the ten MCMC replicate chains. The MDS coordinates are calculated based on the pairwise RF distances computed over all sampled posterior trees. Each dot represents a tree sampled in the posterior whose rainbow color indicates the corresponding iteration index (see inset color gradient bar). The top and bottom rows present the five 400-million-iteration chains ("400M run") and the five 1-billion-iteration chains ("1B run"), respectively. A triangle represents the MCC tree summarized from an MCMC chain, while an open circle indicates the MCC tree summarized across all the replicates of that dataset. Dots are connected with lines to indicate that they are sampled consecutively in the chain.

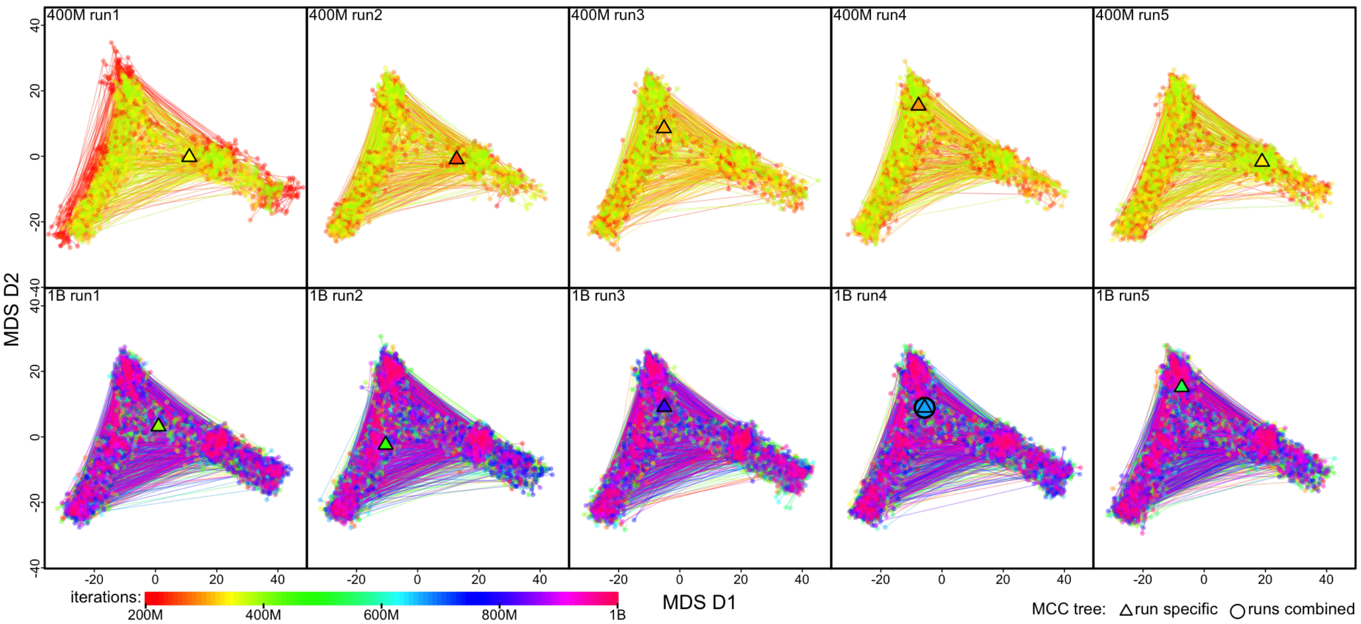


**Figure S57: Tree sampling problems with the Lassa_n551_s7038 dataset appears to be resolved after pruning two problematic sequences (088 and 091).** We re-generated the posterior tree space after pruning the two problematic sequences from each sampled posterior tree; *i.e.*, the distinct tree space here compared with Fig. S56 is likely to be caused solely by those two sequences. See Fig. 4 for details about these problematic sequences; Panel **E** of Fig. 4 shows a compact contrast between Fig. S56 and this figure.

##### S2.4.7 Mumps Virus Dataset

Moncla et al. (7) focus on a mumps virus outbreak in Washington State (WA) in 2016–2017, uncovering the geographic sources of mumps virus introduction to WA and revealing the dynamics of the transmission chains these introductions seeded. To achieve that, they sequenced 166 mumps virus genomes from this outbreak, combined it with all the other available mumps virus genomes collected from North America at the time of the study, and applied quality-control filtrations, resulting in a sequence dataset with 467 genomes and 15393 nucleotide sites with which they conducted the phylodynamic inference.

We directly acquired the BEAST XML scripts provided by the original study from its supplementary repository—containing both the sequence alignment and sampling time and location data—and specify our analyses accordingly. Specifically, following the original study, in our analyses we specify a phylodynamic model with the following components: (1) an HKY+$\Gamma_4$ substitution model (51, 52), (2) a strict molecular clock model, and (3) a Skygrid coalescent node-age model (55).

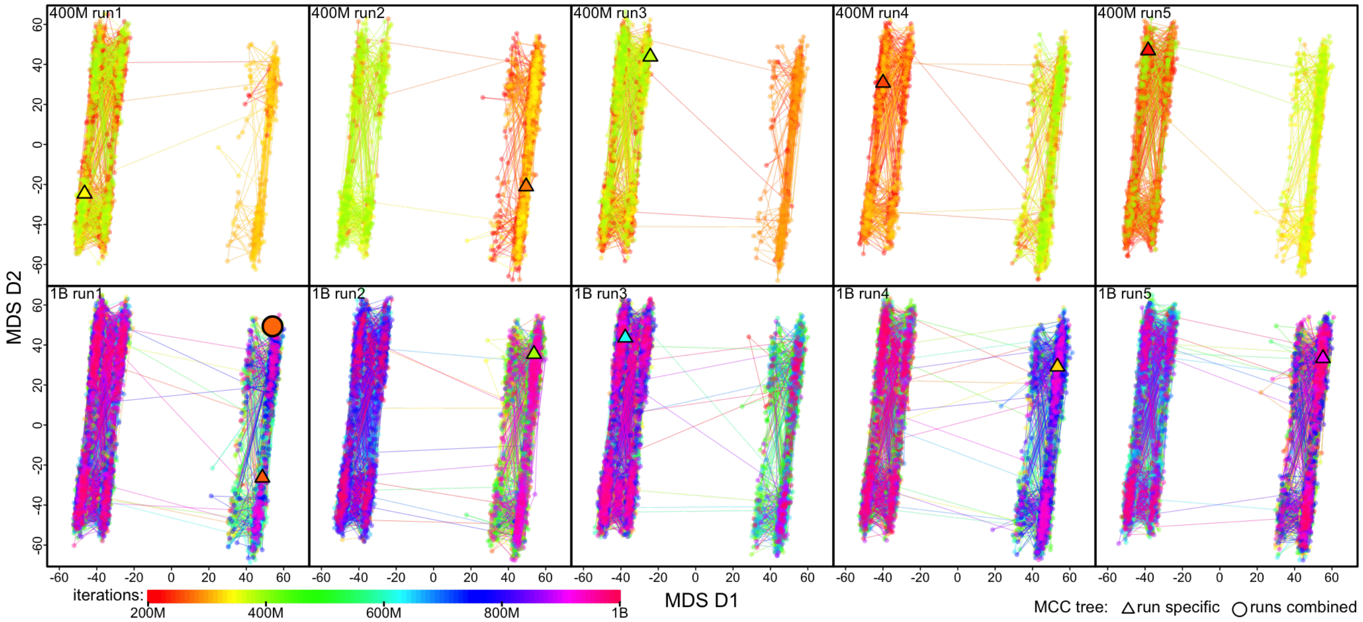


**Figure S58: MCMC sampler progression in tree space of the Mumps_n467_s15393 dataset.** Each panel shows the MDS plot visualizing the posterior tree space inferred with one of the ten MCMC replicate chains. The MDS coordinates are calculated based on the pairwise RF distances computed over all sampled posterior trees. Each dot represents a tree sampled in the posterior whose rainbow color indicates the corresponding iteration index (see inset color gradient bar). The top and bottom rows present the five 400-million-iteration chains ("400M run") and the five 1-billion-iteration chains ("1B run"), respectively. A triangle represents the MCC tree summarized from an MCMC chain, while an open circle indicates the MCC tree summarized across all the replicates of that dataset. Dots are connected with lines to indicate that they are sampled consecutively in the chain.

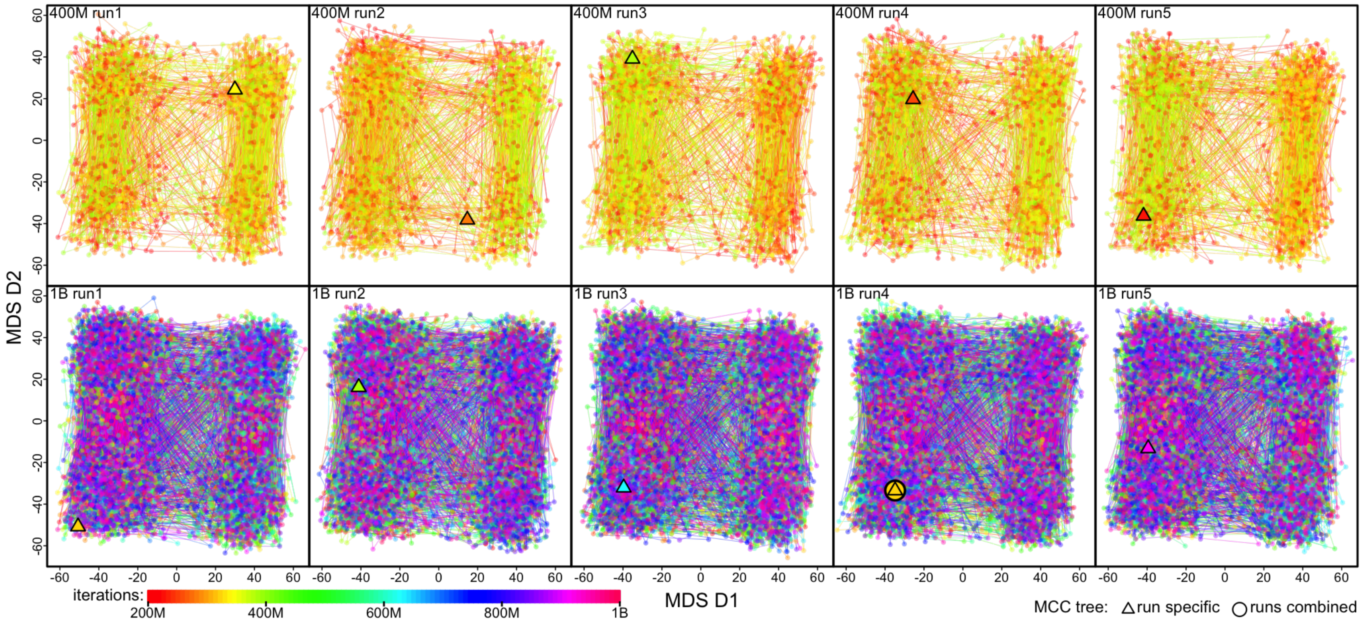


**Figure S59: The poor mixing in topological inference of the Mumps_n467_s15393 dataset appears to be resolved after pruning one sequence (20.16/2).** We re-generated the posterior tree space after pruning the problematic sequence from each sampled posterior tree; *i.e.*, the distinct tree space here compared with Fig. S58 is likely to be caused solely by that sequence. See Fig. S16 for details about the problematic sequence; Panel **F** (left and middle subpanels) of Fig. S16 shows a compact contrast between Fig. S58 and this figure.

##### S2.4.8 Rabies Virus Dataset

Viana et al. (8) explored the effects of culling vampire bat on the geographic dispersal dynamics of vampire bat rabies (VBR) virus by performing a geographic phylodynamic inference with a sequence dataset containing 290 VBR lineage 3 virus genomes (11883 nucleotide sites) collected between 1997 and 2016 in southern Peru. We directly acquired the BEAST XML scripts provided by the original study from its supplementary repository—containing both the sequence alignment and sampling times—and specify our analyses accordingly.

Specifically, following the original study, we divide the genomic alignment into two partitions, where the first partition includes sites at the first and second codon positions, and the second partition includes sites at the third codon position. For each partition, we specify an independent GTR+$\Gamma_4$ substitution model (52, 58) and a partition-specific rate multiplier whose mean is constrained to be one across the partitions. The phylodynamic model also includes an uncorrelated lognormal branch-rate prior model (53, 54) and a Skygrid coalescent node-age model (55).

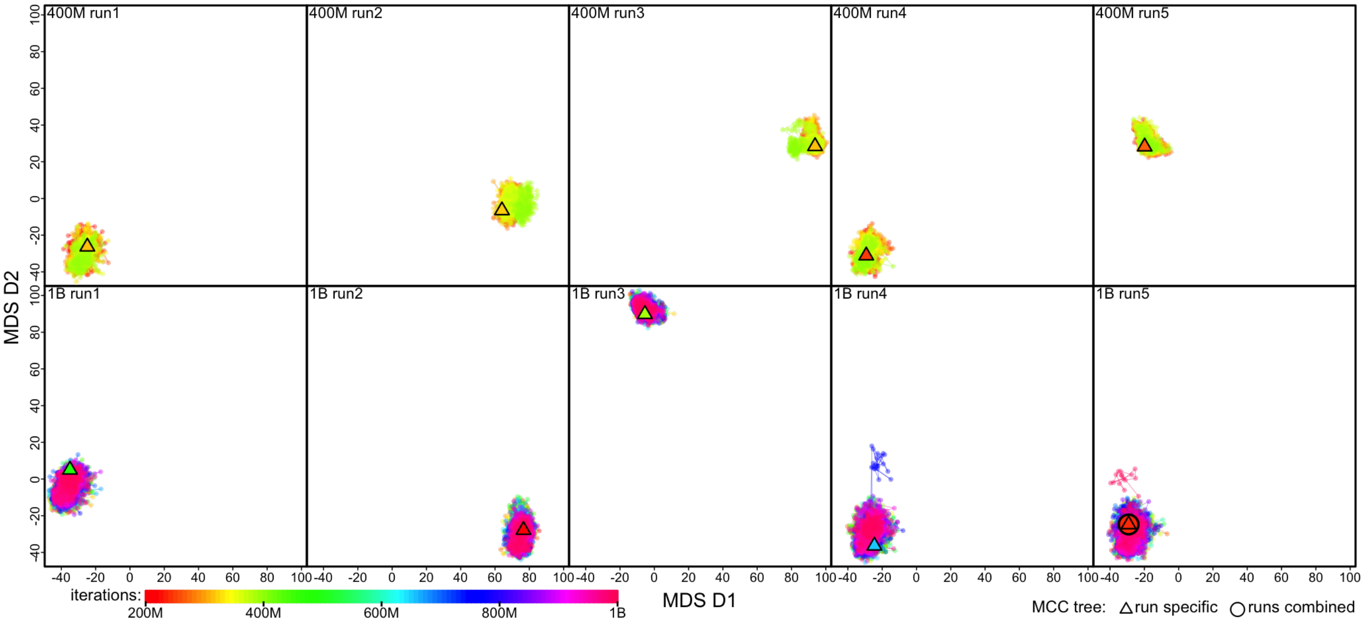


**Figure S60: MCMC sampler progression in tree space of the Rabies_n290_s11883 dataset.** Each panel shows the MDS plot visualizing the posterior tree space inferred with one of the ten MCMC replicate chains. The MDS coordinates are calculated based on the pairwise RF distances computed over all sampled posterior trees. Each dot represents a tree sampled in the posterior whose rainbow color indicates the corresponding iteration index (see inset color gradient bar). The top and bottom rows present the five 400-million-iteration chains (“400M run”) and the five 1-billion-iteration chains (“1B run”), respectively. A triangle represents the MCC tree summarized from an MCMC chain, while an open circle indicates the MCC tree summarized across all the replicates of that dataset. Dots are connected with lines to indicate that they are sampled consecutively in the chain.

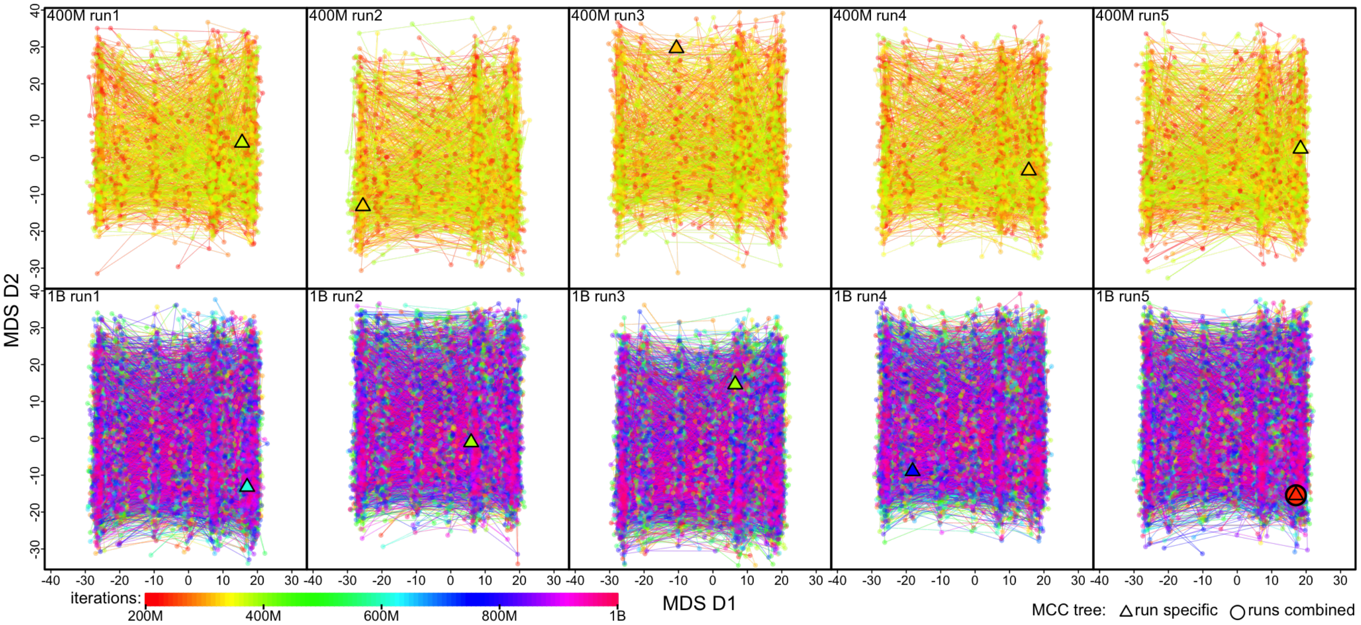


**Figure S61: Tree sampling problems with the Rabies_n290_s11883 dataset appears to be largely resolved after pruning seven problematic sequences.** We re-generated the posterior tree space after pruning the problematic sequences from each sampled posterior tree; *i.e.*, the distinct tree space here compared with Fig. S60 is likely to be caused almost entirely by those sequences. See Fig. S17 for details about the problematic sequences; Panel **C** (left and middle subpanels) of Fig. S17 shows a compact contrast between Fig. S60 and this figure.

##### S2.4.9 West Nile Virus (WNV) Dataset

Dellicour et al. (9) developed an analytical workflow to formally test epidemiological hypotheses via phylodynamic inference and illustrate this workflow by performing phylodynamic analyses with a comprehensive WNV sequence dataset. To construct that comprehensive dataset, they first collected the WNV sequences available on GenBank as of the time of that study (November 2017) and then filtered the sequences with multiple quality-control and subsampling steps, eventually obtaining a final alignment with 801 genomic sequences and 10302 nucleotide sites.

We directly acquired the BEAST XML scripts provided by the original study from its supplementary repository—containing both the sequence alignment and sampling times—and specify our analyses accordingly. Specifically, following the original study, in our analyses we specify a phylodynamic model with the following components: (1) a GTR+$\Gamma_4$ substitution model (52, 58), (2) an uncorrelated lognormal branch-rate prior model (53, 54), and (3) a Skygrid coalescent node-age model (55).

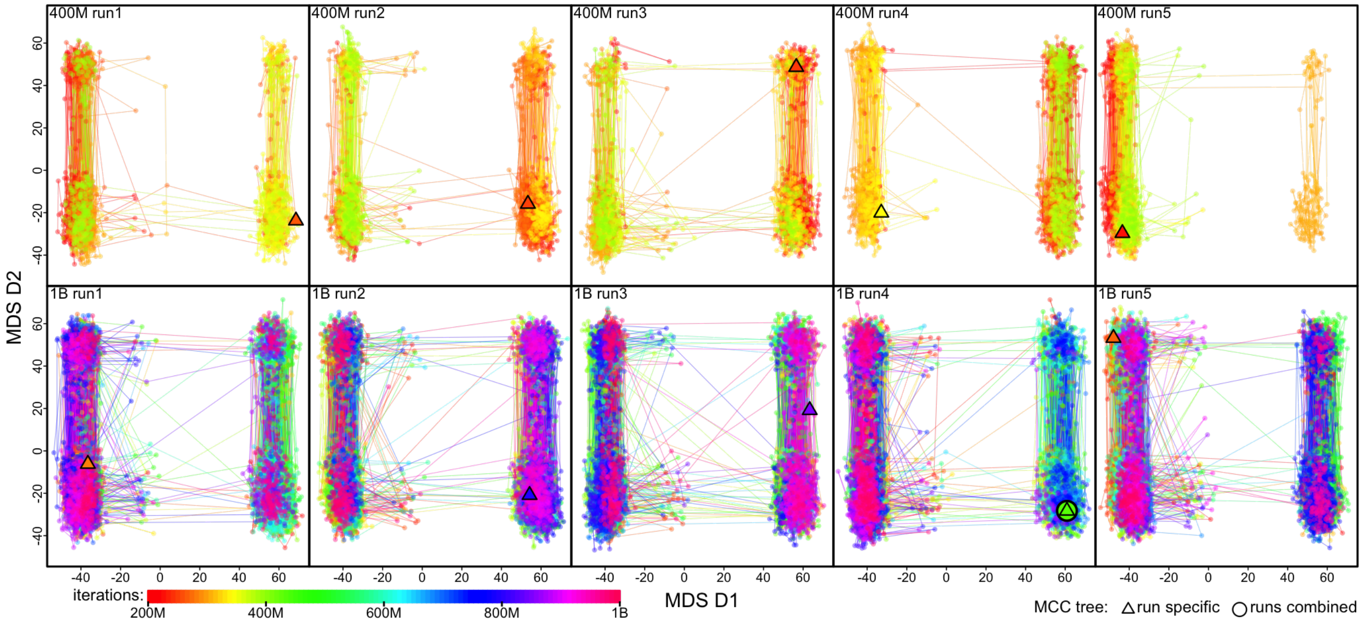


**Figure S62: MCMC sampler progression in tree space of the WNV_n801_s10302 dataset.** Each panel shows the MDS plot visualizing the posterior tree space inferred with one of the ten MCMC replicate chains. The MDS coordinates are calculated based on the pairwise RF distances computed over all sampled posterior trees. Each dot represents a tree sampled in the posterior whose rainbow color indicates the corresponding iteration index (see inset color gradient bar). The top and bottom rows present the five 400-million-iteration chains ("400M run") and the five 1-billion-iteration chains ("1B run"), respectively. A triangle represents the MCC tree summarized from an MCMC chain, while an open circle indicates the MCC tree summarized across all the replicates of that dataset. Dots are connected with lines to indicate that they are sampled consecutively in the chain.

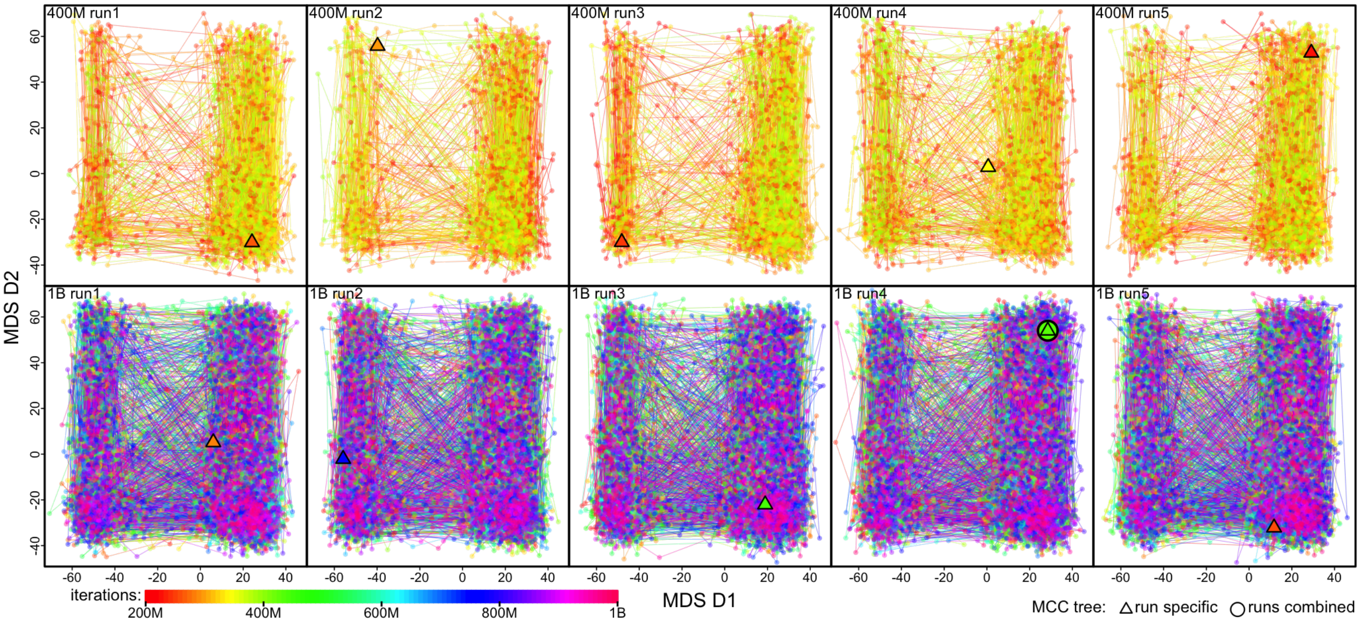


**Figure S63: The poor mixing in topological inference of the WNV_n801_s10302 dataset appears to be resolved after pruning two problematic sequences.** We re-generated the posterior tree space after pruning the problematic sequences from each sampled posterior tree; *i.e.*, the distinct tree space here compared with Fig. S62 is likely to be caused solely by those sequences. See Fig. S18 for details about the problematic sequences; Panel **D** (left and middle subpanels) of Fig. S18 shows a compact contrast between Fig. S62 and this figure.

##### S2.4.10 Zika Virus Dataset

Grubaugh et al. (23) uncovered a hidden Zika outbreak in Cuba by sequencing Zika virus from infected travelers arriving from Cuba during 2017–2018 and performing phylodynamic inference with a combined Zika virus sequence dataset containing the genomes collected in this study and all the publicly available Zika virus genomes of Asian genotype from the Pacific and Americas retrieved from GenBank as of August 2018. This combined dataset comprises 283 Zika virus genomes with 10269 nucleotide sites. We directly acquired the BEAST XML scripts provided by the original study from its supplementary repository—containing both the sequence alignment and sampling times—and specify our analyses accordingly.

Specifically, following the original study, we divide the genomic alignment into three partitions, each corresponding to a codon position. For each partition, we specify an independent HKY+$\Gamma_4$ substitution model (51, 52) and a partition-specific rate multiplier whose mean is constrained to be one across the partitions. The phylodynamic model also includes an uncorrelated lognormal branch-rate prior model (53, 54) and a Skygrid coalescent node-age model (55).

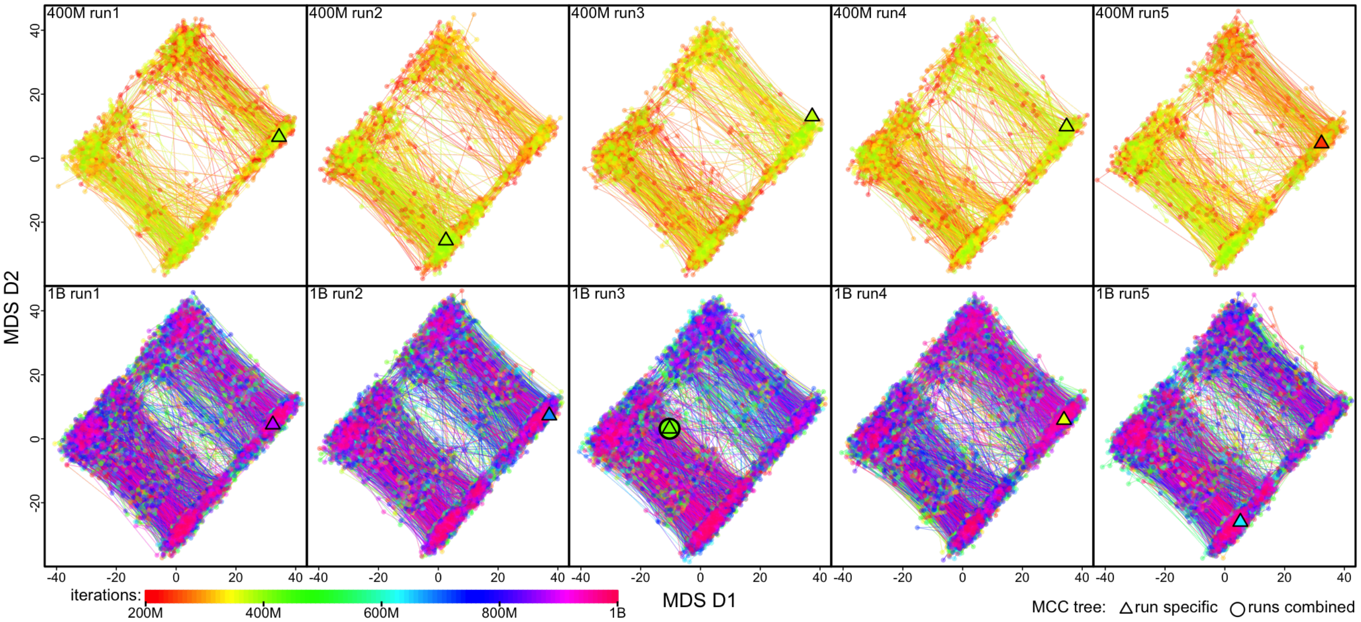


**Figure S64: MCMC sampler progression in tree space of the Zika_n283_s10269 dataset.** Each panel shows the MDS plot visualizing the posterior tree space inferred with one of the ten MCMC replicate chains. The MDS coordinates are calculated based on the pairwise RF distances computed over all sampled posterior trees. Each dot represents a tree sampled in the posterior whose rainbow color indicates the corresponding iteration index (see inset color gradient bar). The top and bottom rows present the five 400-million-iteration chains ("400M run") and the five 1-billion-iteration chains ("1B run"), respectively. A triangle represents the MCC tree summarized from an MCMC chain, while an open circle indicates the MCC tree summarized across all the replicates of that dataset. Dots are connected with lines to indicate that they are sampled consecutively in the chain.

##### S2.4.11 SARS-CoV-2 Origin Dataset

Pekar et al. (6) investigated the zoonotic origins of SARS-CoV-2 by performing comprehensive phylodynamic inference with viral genomic samples of SARS-CoV-2 lineages A and B collected in the early stage of the pandemic (by 14 February 2020). We downloaded the 787 SARS-CoV-2 genomes included in that study from GISAID according to the provided accession IDs, aligned them, and applied quality-control filtration (see Section S2.1 for details), obtaining a final alignment with 717 sequences and 29232 nucleotide sites.

We directly acquired the `BEAST` XML scripts provided by the original study from its supplementary repository—containing both the sequence alignment and sampling times—and specify our analyses accordingly. Specifically, following the original study, in our analyses we specify a phylodynamic model with the following components: (1) a GTR+I model (58, 59) substitution model, (2) a strict molecular clock model, and (3) a Skygrid coalescent node-age model (55).

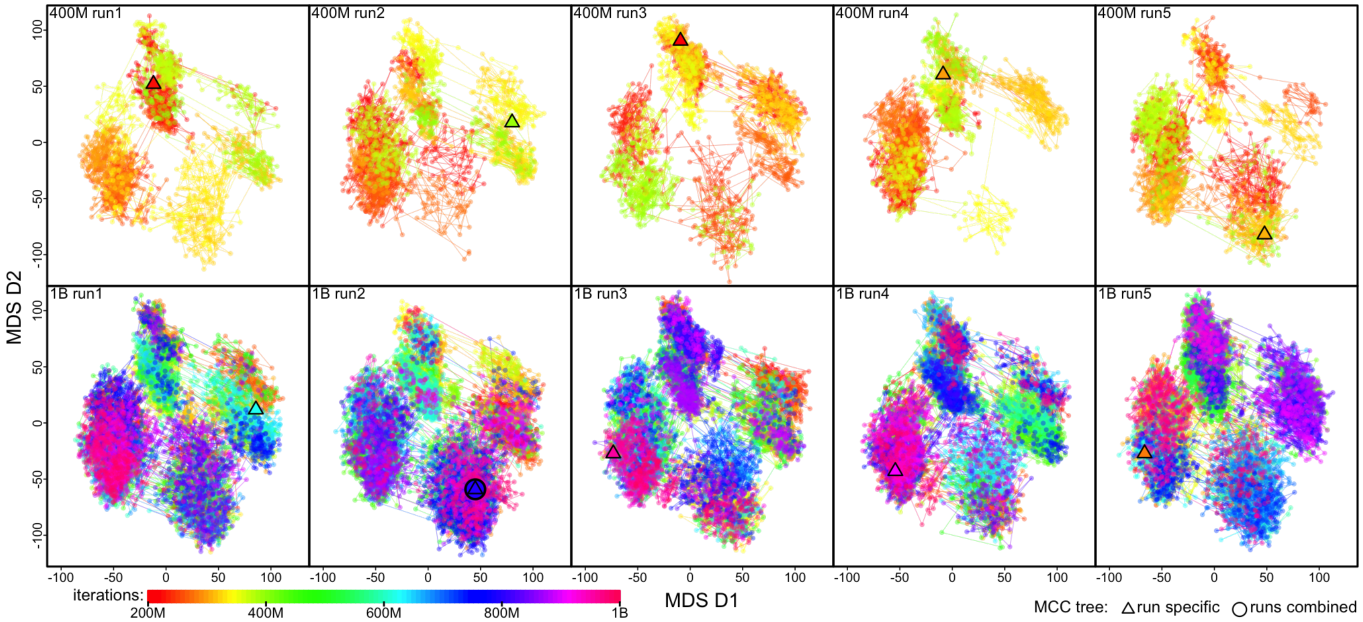


**Figure S65: MCMC sampler progression in tree space of the SARS2_n717_s29232 dataset.** Each panel shows the MDS plot visualizing the posterior tree space inferred with one of the ten MCMC replicate chains. The MDS coordinates are calculated based on the pairwise RF distances computed over all sampled posterior trees. Each dot represents a tree sampled in the posterior whose rainbow color indicates the corresponding iteration index (see inset color gradient bar). The top and bottom rows present the five 400-million-iteration chains ("400M run") and the five 1-billion-iteration chains ("1B run"), respectively. A triangle represents the MCC tree summarized from an MCMC chain, while an open circle indicates the MCC tree summarized across all the replicates of that dataset. Dots are connected with lines to indicate that they are sampled consecutively in the chain.

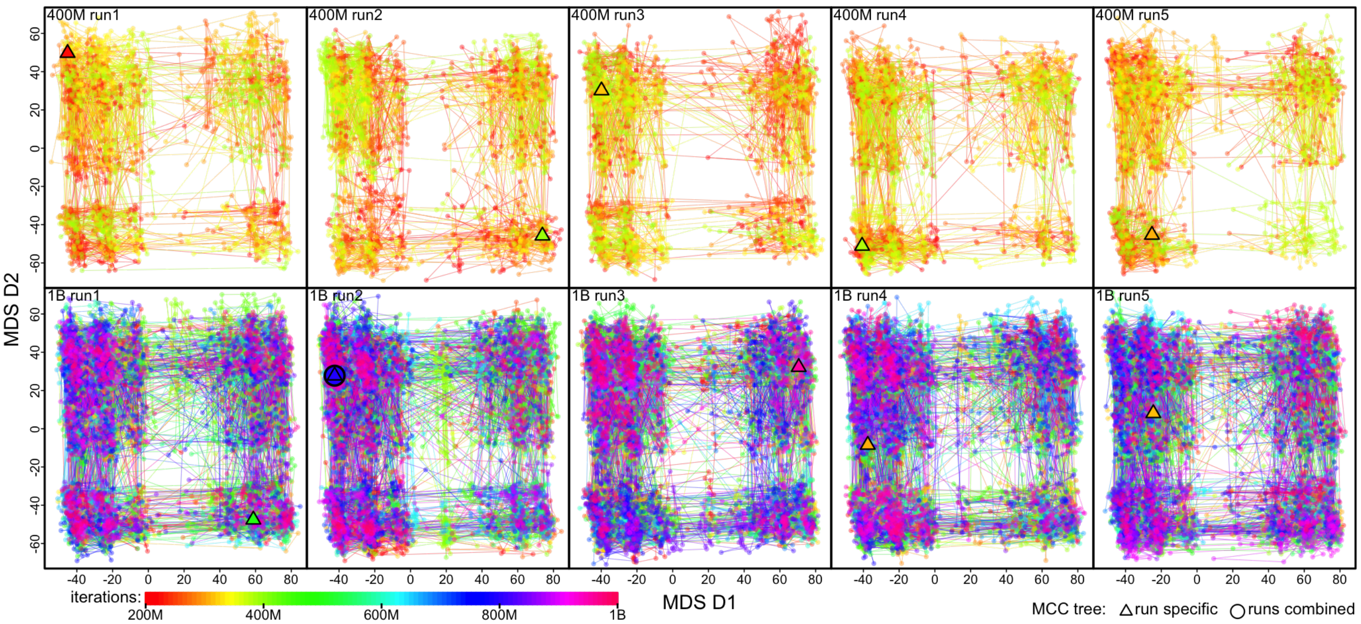


**Figure S66: The poor mixing in topological inference of the SARS2_n717_s29232 dataset appears to be resolved after pruning the problematic sequences.** We re-generated the posterior tree space after pruning the three problematic sequences from each sampled posterior tree; *i.e.*, the distinct tree space here compared with Fig. S65 is likely to be caused solely by those sequences. See Fig. S15 for details about the problematic sequences; Panel **C** of Fig. S15 shows a compact contrast between Fig. S65 and this figure.

##### S2.4.12 SARS-CoV-2 Brazil Dataset

Candido et al. (12) investigated the early spread of the SARS-CoV-2 epidemic in Brazil and the efficacy of mitigation measures on limiting that spread. They combined newly sequenced SARS-CoV-2 genomes sampled from Brazil with the genomes available on GISAID as of April 24, 2020 to produce a SARS-CoV-2 sequence dataset focusing on the epidemic in Brazil. This dataset contains 1182 SARS-CoV-2 genomes, including 490 sampled from Brazil and 692 subsampled from the sequences collected outside of Brazil. We downloaded these SARS-CoV-2 genomes from GISAID according to the provided accession IDs, aligned them, and applied quality-control filtration (see Section S2.1 for details), obtaining a final alignment with 1046 sequences and 29409 nucleotide sites.

We directly acquired the `BEAST` XML scripts provided by the original study from its supplementary repository—containing both the sequence alignment and sampling time and location data—and specify our analyses accordingly. Specifically, following the original study, in our analyses we specify a phylodynamic model with the following components: (1) an HKY+$\Gamma_4$ substitution model (51, 52), (2) a strict molecular clock model, and (3) a Skygrid coalescent node-age model (55).

The original study performed phylogeographic analyses under three different space discretization schemes; here, we used their scheme B—including five Brazilian regions ("Southeast", "Northeast", "North", "Centre-West", and "South") and five international regions (North America, Europe, Asia, Oceania, and Africa)—for our parsimony phylogeographic reconstruction.

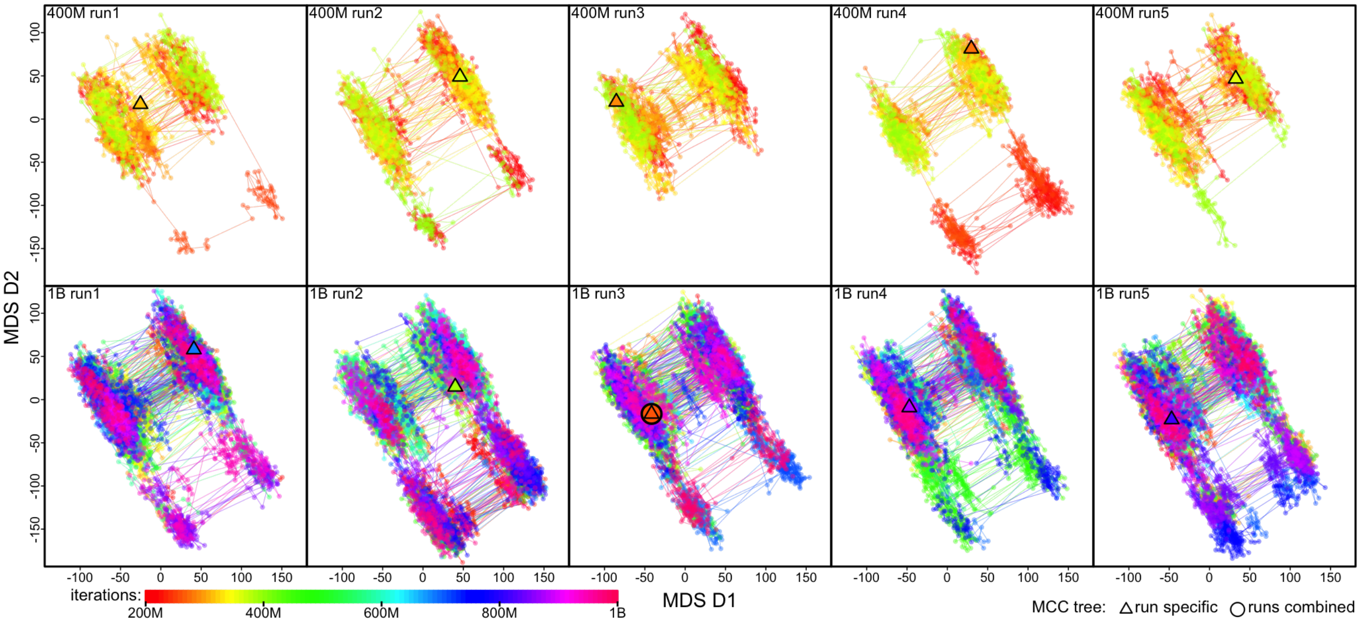


**Figure S67: MCMC sampler progression in tree space of the SARS2_n1046_s29409 dataset.** Each panel shows the MDS plot visualizing the posterior tree space inferred with one of the ten MCMC replicate chains. The MDS coordinates are calculated based on the pairwise RF distances computed over all sampled posterior trees. Each dot represents a tree sampled in the posterior whose rainbow color indicates the corresponding iteration index (see inset color gradient bar). The top and bottom rows present the five 400-million-iteration chains ("400M run") and the five 1-billion-iteration chains ("1B run"), respectively. A triangle represents the MCC tree summarized from an MCMC chain, while an open circle indicates the MCC tree summarized across all the replicates of that dataset. Dots are connected with lines to indicate that they are sampled consecutively in the chain.

##### S2.4.13 SARS-CoV-2 Europe Dataset

Lemey et al. (13) studied the relative contribution of newly introduced SARS-CoV-2 lineages versus the persistent lineages to the resurgence of COVID-19 in late summer 2020. They assembled a viral sequence dataset based on all available SARS-CoV-2 genomes from European countries in GISAID on 3 November 2020; after downsampling the dataset to mitigate spatiotemporal disparities in sequence data collection and applying quality-control filtration, their final dataset consisted of 3959 genomes. We downloaded these SARS-CoV-2 genomes from GISAID according to the provided accession IDs, aligned them, and applied additional quality-control filtration steps (see Section S2.1 for details), obtaining a final alignment with 3241 sequences and 29409 nucleotide sites.

We directly acquired the BEAST XML scripts provided by the original study from its supplementary repository—containing both the sequence alignment and sampling time and location data—and specify our analyses accordingly. Specifically, following the original study, in our analyses we specify a phylodynamic model with the following components: (1) an HKY+$\Gamma_4$ substitution model (51, 52), (2) a strict molecular clock model, and (3) a Skygrid coalescent node-age model (55).

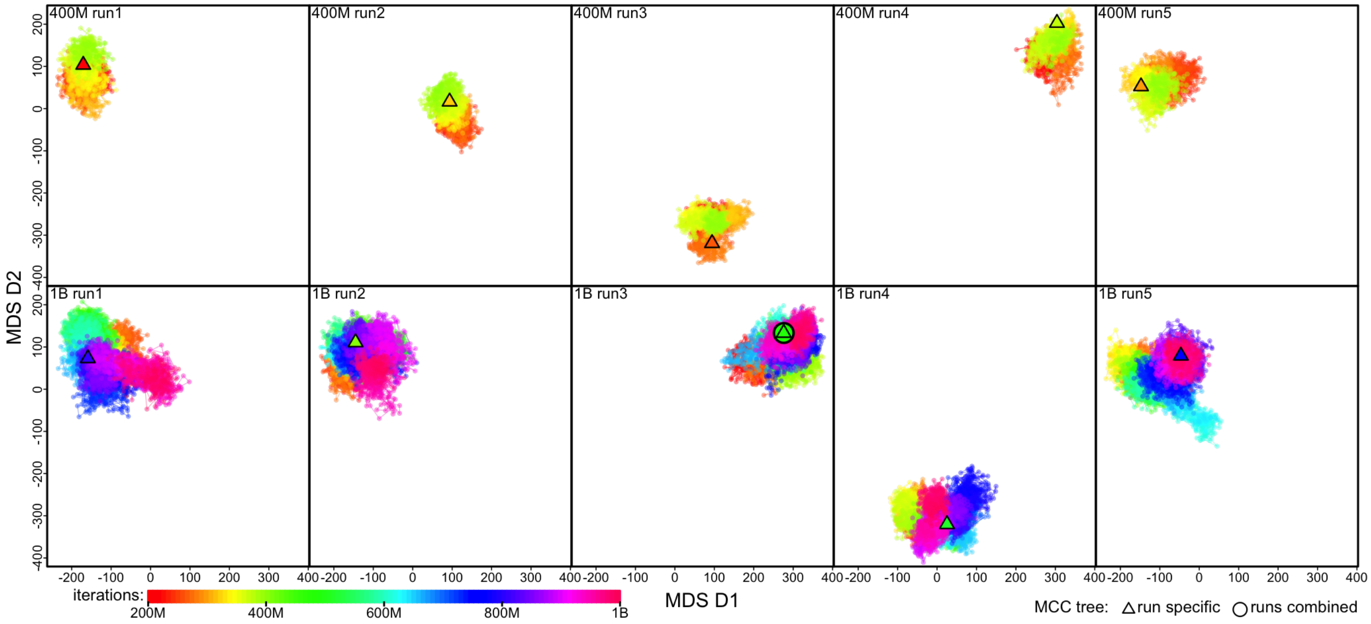


**Figure S68: MCMC sampler progression in tree space of the SARS2_n3241_s29409 dataset.** Each panel shows the MDS plot visualizing the posterior tree space inferred with one of the ten MCMC replicate chains. The MDS coordinates are calculated based on the pairwise RF distances computed over all sampled posterior trees. Each dot represents a tree sampled in the posterior whose rainbow color indicates the corresponding iteration index (see inset color gradient bar). The top and bottom rows present the five 400-million-iteration chains ("400M run") and the five 1-billion-iteration chains ("1B run"), respectively. A triangle represents the MCC tree summarized from an MCMC chain, while an open circle indicates the MCC tree summarized across all the replicates of that dataset. Dots are connected with lines to indicate that they are sampled consecutively in the chain.
